# Localization-dependent activation of the DEAD-box ATPase Vasa by eLOTUS domains

**DOI:** 10.64898/2026.03.28.715021

**Authors:** Asen Garbelyanski, David Hauser, Alberto Trolese, Prateek Kumar, Hin Hark Gan, Caroline de Almeida, Matthias P. Meyer, Maria Hondele, Teresa Carlomagno, Kristin C. Gunsalus, Mandy Jeske

## Abstract

DEAD-box RNA helicases remodel RNA structures in many cellular pathways, yet how their activity is spatially controlled in cells remains poorly understood. The *Drosophila* germline helicase Vasa functions in ovaries only when localized to cytoplasmic granules, a process mediated by eLOTUS-domain proteins. Here, we define the mechanism by which eLOTUS domains activate Vasa. Biochemical analyses reveal that Vasa alone is largely inactive. eLOTUS domains bind the open conformation of Vasa and promote formation of the closed RNA- and ATP-bound state by accelerating RNA engagement. This stimulation requires a positively charged intrinsically disordered sequence within eLOTUS that increases RNA association. Mutations in this element abolish Vasa stimulation while preserving binding. We demonstrate that Vasa activation is essential *in vivo*. Together, these findings reveal a localization-dependent mechanism for regulating the DEAD-box helicase Vasa, in which enzymatic activity is gated by a spatially restricted cofactor.

## INTRODUCTION

RNA helicases are enzymes that use ATP hydrolysis to remodel RNA and RNA-protein complexes, participating in virtually all aspects of RNA metabolism (Jarmoskaite & Russell, 2014; Bohnsack *et al*, 2023; Khemici & Linder, 2016). The DEAD-box family is the largest class of RNA helicases involved in nearly every stage of RNA life, from transcription and processing to translation and decay (Linder & Jankowsky, 2011). These proteins share a conserved core of two RecA-like domains connected by a flexible linker, with the N-terminal domain harboring the signature Asp-Glu-Ala-Asp motif essential for ATP hydrolysis (Linder *et al*, 1989; Jankowsky & Fairman, 2007; Fairman-Williams *et al*, 2010). DEAD-box proteins unwind short RNA duplexes non-processively, binding transiently exposed single-stranded regions and bending the RNA to destabilize base pairing (Yang *et al*, 2007; Sengoku *et al*, 2006). ATP binding promotes stable RNA engagement, whereas hydrolysis triggers release of the helicase from the substrate (Chen *et al*, 2008; Liu *et al*, 2008).

In cells, DEAD-box proteins often act within larger ribonucleoprotein (RNP) complexes where they can remodel RNA secondary structures, displace proteins from single-stranded RNA, or serve as regulated RNA-binding factors (Jarmoskaite & Russell, 2014; Leitão *et al*, 2015; Jankowsky, 2011). Many DEAD-box ATPases are intrinsically slow catalysts. In the absence of substrates, the RecA-like domains adopt a flexible open conformation, disfavoring efficient RNA and ATP binding (Caruthers *et al*, 2000; Bono *et al*, 2006; Andersen *et al*, 2006). Productive substrate engagement requires closure of the core around RNA and ATP, which limits the rate for substrate association (Ozgur *et al*, 2015). In many DEAD-box proteins, phosphate release following ATP hydrolysis constitutes the rate-limiting step in the catalytic cycle (Henn *et al*, 2008; Hilbert *et al*, 2011; Cao *et al*, 2011; Wong *et al*, 2016).

To overcome these kinetic limitations, DEAD-box ATPases often rely on activating cofactors, which can modulate distinct steps of the reaction cycle, including RNA binding, conformational transitions, or product release (Sloan & Bohnsack, 2018; Donsbach & Klostermeier, 2021). For example, MIF4G domains accelerate the cycle by binding to both RecA-like domains on the side opposite the ATP-binding cleft, restricting the conformational space and preventing adoption of fully inactive orientations (Oberer *et al*, 2005; Schütz *et al*, 2008; Montpetit *et al*, 2011; Chen *et al*, 2014; Mathys *et al*, 2014; Pühringer *et al*, 2020; Schuller *et al*, 2020). In contrast to these well-characterized examples, the molecular mechanisms by which other factors stimulate DEAD-box helicases remain poorly understood. Understanding how binding partners modulate DEAD-box activity is essential for linking biochemical to cellular and developmental function.

Vasa (DDX4) is among the first DEAD-box proteins identified (Lasko & Ashburner, 1988; Linder *et al*, 1989; Hay *et al*, 1988), and the crystal structure of the RNA- and nucleotide-bound Vasa core was pivotal in establishing a general model for local duplex unwinding by DEAD-box ATPases (Sengoku *et al*, 2006). Vasa is highly conserved across metazoans and predominantly expressed in the germline (Lasko, 2013; Raz, 2000). Studies in *Drosophila* have established Vasa as a key regulator of oogenesis, germ cell development, and embryonic patterning (Schüpbach & Wieschaus, 1986; Dehghani & Lasko, 2017). At the cellular level, Vasa contributes to translation regulation and piRNA-mediated transposon repression, with the latter being essential for genome defense in gonads (Liu *et al*, 2009; Carrera *et al*, 2000; Johnstone & Lasko, 2004; Xiol *et al*, 2014; Nishida *et al*, 2015; Kuramochi-Miyagawa *et al*, 2010; Wenda *et al*, 2017). Given its central roles in multiple germline processes, Vasa activity must be precisely regulated in space and time, but the mechanisms by which cofactors control this remains poorly understood.

Previous work identified the extended LOTUS (eLOTUS) domain as an activator of Vasa (Jeske *et al*, 2017). LOTUS domains are helix-turn-helix motifs highly conserved across all domains of life (Callebaut & Mornon, 2010; Anantharaman *et al*, 2010). In animals, LOTUS domain proteins play prominent roles during gametogenesis and early embryogenesis (Kubíková *et al*, 2020; Cipriani *et al*, 2021; Price *et al*, 2021). Vasa binding requires an α-helical C-terminal extension that is present in eLOTUS, but not in minimal LOTUS (mLOTUS) domains (Jeske *et al*, 2017). The crystal structure of the eLOTUS-Vasa complex revealed that the eLOTUS domain binds the C-terminal RecA-like domain (CTD) of Vasa (Jeske *et al*, 2017) in a mode distinct from previously characterized DEAD-box protein activators, yet how this binding translates into stimulation of ATPase activity remains unknown.

In *Drosophila,* the proteins Oskar, Tejas (TDRD5), and Tapas (TDRD7) contain eLOTUS domains, all of which can stimulate Vasa *in vitro* (Jeske *et al*, 2017), but act in distinct biological contexts and in distinct subcellular sites. At the posterior pole of the oocyte, Oskar cooperates with Vasa to assemble the germ plasm, required for germ cell specification and embryonic patterning (Ephrussi & Lehmann, 1992; Ephrussi *et al*, 1991). In nurse cells, Tejas and Vasa localize to perinuclear nuage and are essential for piRNA-mediated transposon silencing, which preserves genome integrity in gonads (Patil & Kai, 2010; Handler *et al*, 2011; Ozata *et al*, 2019; Malone *et al*, 2009). Tapas shares a similar domain organization with Tejas but plays a non-essential, non-redundant role in the piRNA pathway (Handler *et al*, 2011; Patil *et al*, 2014). Interaction with eLOTUS domains is critical for Vasa recruitment to germ plasm and nuage (Jeske *et al*, 2017). The distinct subcellular localization of Vasa-eLOTUS complexes raises the possibility that eLOTUS-mediated stimulation of Vasa could have context-specific functional consequences.

Here, we combine *in vitro* biochemical assays, structural modeling, and *in vivo* genetic studies to dissect the mechanism by which eLOTUS domains activate Vasa. We show that Vasa alone has very poor ATPase activity, even in the presence of RNA. The eLOTUS domain specifically binds the open conformation of Vasa and promotes transition to the closed, RNA- and ATP-bound state. This stimulation requires an unstructured, positively charged stretch within eLOTUS that enhances RNA engagement. Mutation of this unstructured region abolishes stimulation without affecting Vasa binding, revealing the functional importance of Vasa activation uncoupled from localization *in vivo*. Swapping the eLOTUS domains of Oskar and Tejas leads to severe defects *in vivo,* consistent with their stimulatory roles in distinct cellular pathways. Together, these findings reveal a previously unknown strategy by which a regulatory domain uses an intrinsically disordered element to activate a DEAD-box protein, a modular activation mechanism that is tightly linked to subcellular localization with broad implications for RNA metabolism and early animal development.

## RESULTS

### The Tejas-eLOTUS - Vasa interaction is low affinity yet critical for germline function *in vivo*

To understand how eLOTUS domains regulate Vasa, we focused on the Tejas-eLOTUS domain. This domain is monomeric in solution (**Supplementary Figure 1A**), making the mechanistic interpretation of the data more straightforward than with the dimeric Oskar-eLOTUS domain (**Supplementary Figure 1B**) (Jeske *et al*, 2017). We first characterized the interaction between Tejas and Vasa (**Figure 1A)** *in vitro* to guide mutant design for subsequent studies. Isothermal titration calorimetry revealed that Tejas-eLOTUS binds Vasa-CTD with a dissociation constant of ∼20 µM (**Figure 1B**), comparable to the ∼10 µM reported for Oskar-eLOTUS (Jeske *et al*, 2017). Based on the Oskar-eLOTUS - Vasa-CTD crystal structure, we built a homology model of the Tejas complex and identified two residues in Tejas-eLOTUS (S16 and I85) that correspond to residues of Oskar-eLOTUS essential for Vasa binding (**Figure 1C**). Consistently, mutation of these residues (S16E/I85E; hereafter SI) abolished binding in GST pull-down assays (**Figure 1D**) and strongly reduced stimulation of ATP hydrolysis and dsRNA unwinding by Vasa (**Figures 1E and 1F**).

**FIGURE 1.**
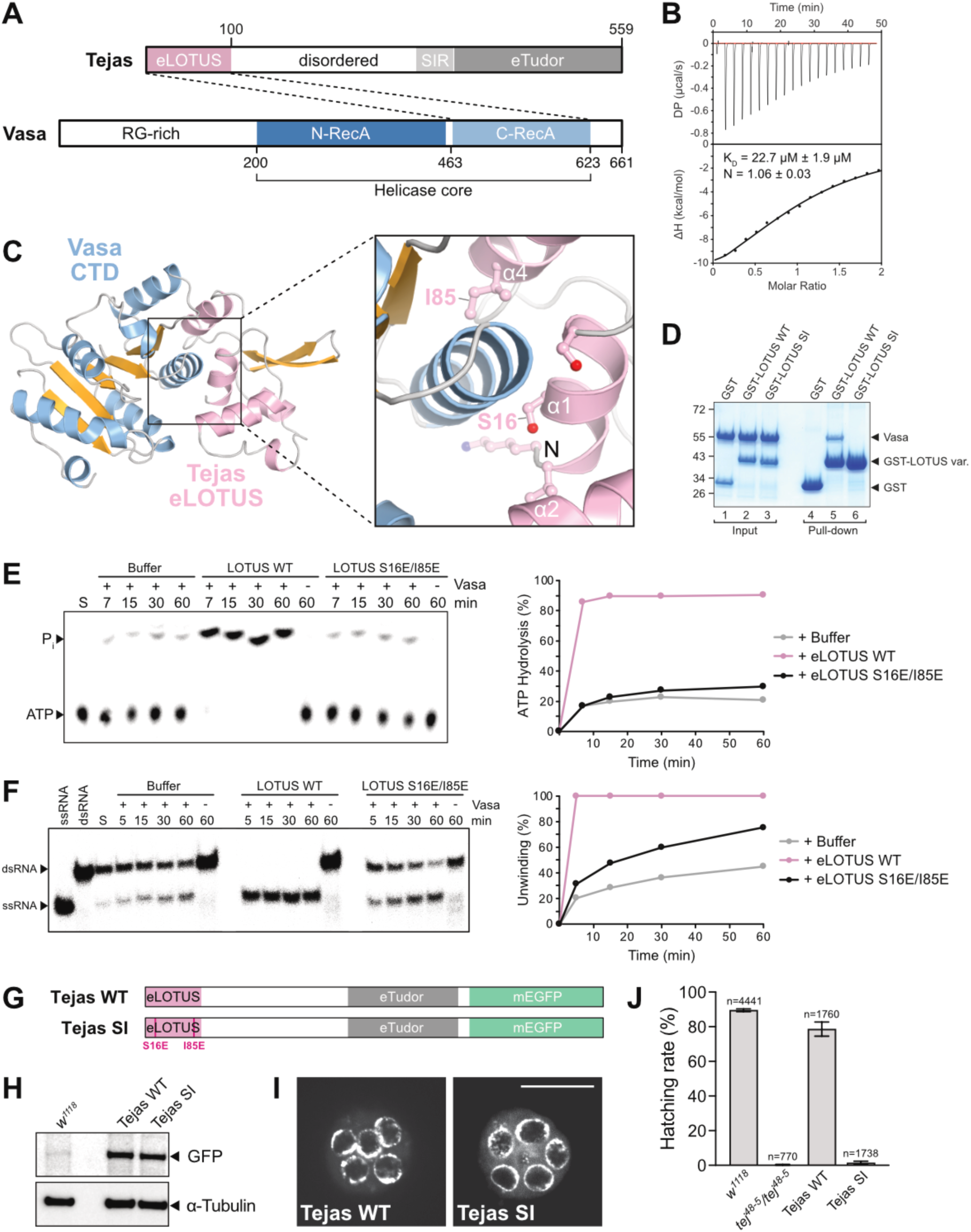
The Tejas-eLOTUS - Vasa interaction is low affinity yet critical for germline function *in vivo*. (A) Domain organization of Tejas and Vasa. The eLOTUS domain of Tejas interacts with the C-RecA-like domain of Vasa. SIR, Spindle E-interacting region; RG-rich, arginine/glycine-rich. (B) ITC analysis of the complex formed by His-Tejas-eLOTUS (aa 1-100) and His-Vasa-CTD (aa 463-661). (C) SWISS-MODEL-generated (Waterhouse et al. 2018) homology model of the complex between Tejas-eLOTUS and Vasa-CTD using the Oskar-eLOTUS - Vasa-CTD crystal structure (PDB ID 5NT7) as a template. The S16 and I85 residues shown in ball-and-stick representation were mutated (SI) in subsequent studies to disrupt Vasa binding. (D) GST pull-down assay using 10 µM GST-Tejas-eLOTUS WT or SI mutant protein and 20 µM His-Vasa 200-623. Molecular weight marker (in kDa) is shown on the left. (E) ATPase assay using 2.5 µM of the His-Vasa 200-661*, 10 µM R26 ssRNA oligo, 8 nM [γ-^32^P] ATP, in the presence of protein buffer, 100 µM wildtype or mutant Tejas-eLOTUS domain. (F) dsRNA unwinding assay in the presence of 3 mM ATP, 25 nM labeled R13 dsRNA oligo, and 500 nM unlabeled competitor R13 ssRNA oligo using 10 µM His-Vasa 200-623 ("+ Vasa") and 35 µM of the wildtype or mutant Tejas-eLOTUS domain. (G) Scheme of the Tejas-WT and Tejas-SI transgenes. Transgenes were analyzed in the *tej^48-5^/tej^48-5^*null background. (H) Western blot analysis of ovary samples using anti-GFP or anti-α-Tubulin antibodies. (I) Live confocal fluorescence microscopy of stage 3 egg chambers showing the localization of the Tejas-mEGFP transgenes indicated. Scale bar is 25 µm. (J) Hatching rate of eggs laid by females with the genotype as indicated. n indicates the number of eggs scored.

To test the biological relevance of this interaction, we generated a wildtype Tejas transgene with a C-terminal mEGFP tag (Tejas-WT) under the control of a UAS promotor and a variant carrying the SI mutation (Tejas-SI) (**Figure 1G**), and expressed them using the germline-specific *oskar*-GAL4 driver in *tej^48-5^/tej^48-5^*null ovaries. Western blot analysis confirmed comparable expression of Tejas-WT and Tejas-SI, indicating that the SI mutation does not destabilize the protein (**Figure 1H**). Immunofluorescence showed proper localization of both transgenes to nuage (**Figures 1I**), as well as normal Vasa localization (**Supplementary Figure 1C**), consistent with previous observations that loss of Tejas does not mislocalize Vasa, likely due to compensation by the related nuage-localized eLOTUS-domain protein Tapas (Patil & Kai, 2010; Patil *et al*, 2014). Phenotypic analyses revealed that egg-laying by *tej^48-5^/tej^48-5^*females was reduced ∼5-fold relative to *w^1118^* controls and that hatching was abolished, consistent with prior *tejas* studies (Patil & Kai, 2010; Patil *et al*, 2014; Handler *et al*, 2011) (**Figure 1J, Supplementary Figure 1D**). Both Tejas-WT and Tejas-SI partially rescued egg-laying (∼50% of *w^1118^*), whereas only Tejas-WT restored hatching (**Figure 1J, Supplementary Figure 1D**). These results demonstrate that the Tejas-eLOTUS-Vasa interaction is essential for germline function *in vivo* and required beyond sole recruitment of Vasa to the nuage.

### The eLOTUS domain binds the open conformation of Vasa and promotes closure

DEAD-box proteins adopt distinct open and closed conformations depending on ATP and RNA binding (Linder & Jankowsky, 2011). To experimentally stabilize these states in Vasa, we used two well-characterized mutations: K295N in the Walker A motif, which prevents ATP binding thus keeping Vasa in an open conformation (Vasa-open), and E400Q in the Walker B motif (DEAD), which allows ATP and RNA binding but blocks product release, stabilizing the closed state (Vasa-closed) (Xiol *et al*, 2014; Walker *et al*, 1982; Gorbalenya *et al*, 1988) (**Figure 2A**).

**FIGURE 2.**
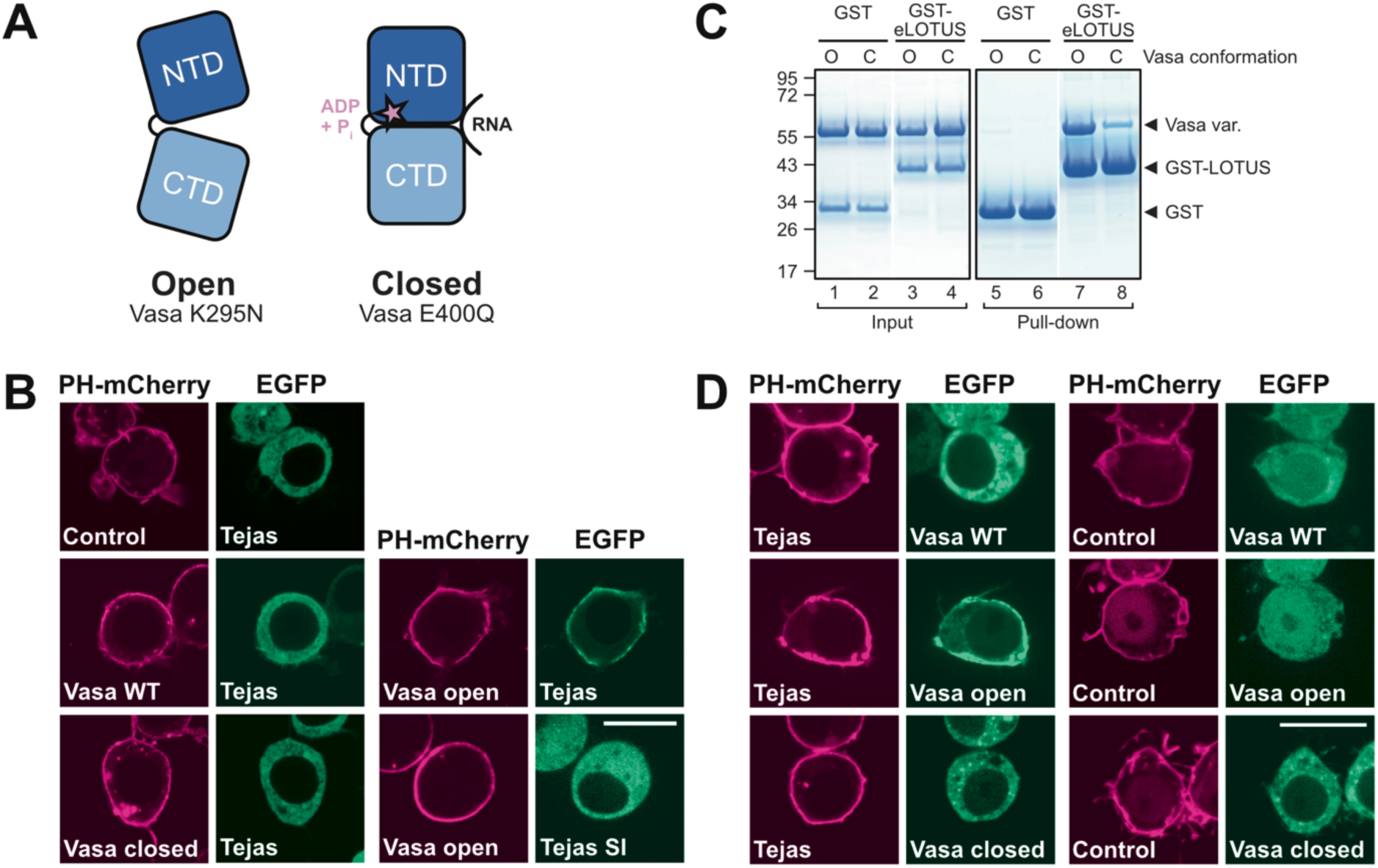
eLOTUS binds the open conformation of Vasa and promotes closure. (A) Pictorial representation of Vasa N- and C-terminal RecA-like domains (NTD and CTD, respectively) in the open and closed conformations. Vasa-open and Vasa-closed carry the K295N or E400Q mutations, respectively. Adapted from Figure 1 of Salgania et al. (Salgania et al. 2024). (B) Tejas interacts with Vasa-open but not with Vasa-WT or Vasa-closed in ReLo assays. Proteins were fused with PH-mCherry or EGFP as indicated and transiently coexpressed in S2R+ cells. After two days, subcellular protein localization was examined by confocal live fluorescence microscopy. ‘Control’ indicates a plasmid expressing PH-mCherry only. Scale bar is 10 µm. (C) GST pull-down assay using 1 nmol GST or GST-Tejas-eLOTUS and 2 nmol His-Vasa 200-623. For the open conformation (’O’), the K295N mutant variant was used. For the closed conformation (’C’), the WT core was incubated with 40 µM of the R13 ssRNA oligo and 2 mM ATP prior to incubation with the LOTUS domain. Molecular weight marker (in kDa) is indicated at the left. Input and pull-down samples originated from one experiment but were loaded onto two gels. Lanes with irrelevant data were removed. (D) Coexpression of Tejas changes the localization of Vasa-WT in ReLo assays from nuclear and cytoplasmic (no Tejas) to cytoplasmic only (with Tejas). ‘Control’ indicates a plasmid expressing PH-mCherry only. In most of the cells, Tejas does not relocalize to the plasma membrane in the presence of Vasa WT (**Supplementary Figure 2C**). Scale bar is 10 µm.

To test whether Tejas-eLOTUS discriminates between these conformations in cells, we employed the ReLo assay, which reports direct protein-protein interactions through relocalization of one fluorescently tagged protein to another anchored at the plasma membrane via a PH domain (Salgania *et al*, 2024). For ReLo, *Drosophila* S2R+ cells are used, which do not express Vasa and Tejas endogenously. Upon cotransfection, EGFP-Tejas robustly relocalized with PH-mCherry-Vasa-open but not with Vasa-closed, demonstrating a clear preference for the open conformation (**Figure 2B**). This behavior aligns with previous cell-based observations for Tejas and Oskar (Salgania *et al*, 2025; Jeske *et al*, 2017). Vasa-WT did not show detectable Tejas interaction under these conditions, likely reflecting a predominantly closed RNA-bound state in Tejas-expressing cells.

We next tested this conformational preference biochemically. In GST pull-down assays using purified GST-Tejas-eLOTUS and recombinant Vasa helicase cores, the open Vasa mutant bound efficiently (**Figure 2C**). Because the closed mutant remained persistently associated with nucleic acids, preventing its purification to homogeneity, we recapitulated this state by incubating Vasa-WT with ATP and a R13 ssRNA oligonucleotide. Under these conditions, Vasa bound stably to the ssR13 in EMSAs (**Supplementary Figure 2A**), and Tejas-eLOTUS binding was markedly diminished compared to the open mutant in the pull-down experiments (**Figure 2C**). Similar results were obtained with Oskar-eLOTUS (**Supplementary Figure 2B**), indicating that preferential recognition of the open state is a shared property of eLOTUS domains.

Finally, we perfromed a reciprocal ReLo assay, in which Tejas was anchored at the plasma membrane and Vasa was unanchored. Unanchored Vasa showed different subcellular localization depending on the mutation: EGFP-Vasa-open localized throughout the cytoplasm and nucleus, whereas EGFP-Vasa-closed remained restricted to the cytoplasm (**Figure 2D**). Under these conditions, Vasa-WT resembled the open mutant. Upon coexpression of Tejas, Vasa-open was efficiently recruited to the membrane, whereas Vasa-closed and Vasa-WT were not, consistent with the first ReLo experiment. Strikingly, in the presence of Tejas, Vasa-WT shifted from a nucleo-cytoplasmic localization resembling the open mutant to a predominantly cytoplasmic distribution resembling the closed mutant (**Figure 2D**). This finding suggests that in cells Tejas binding drives Vasa-WT from an open into a relatively long-lived closed state before dissociating.

Together, these data show that the eLOTUS domain of Tejas recognizes the open conformation of the Vasa helicase core, promotes its transition to the closed state, and dissociates thereafter.

### eLOTUS promotes Vasa activity primarily by enhancing RNA binding

To dissect the mechanism of Vasa activation by eLOTUS domains, we measured the kinetics of ATP hydrolysis using purified proteins and an NADH-coupled assay with an ATP-regenerating system to maintain nearly constant ATP concentrations throughout the measurement (Guo & Pyle, 2022) (**Supplementary Figure 3A**).

We first titrated a ssRNA oligo (R26) in the presence of 5 mM ATP. Vasa alone displayed very low basal ATPase activity even in the presence of RNA (**Figure 3A**). Addition of saturating Tejas-eLOTUS strongly stimulated Vasa, increasing its catalytic efficiency (*k_cat_*/K_M_) by 465-fold. While *k_cat_* increased ∼2.4-fold, the apparent K_M_ for RNA decreased by ∼190-fold, indicating that eLOTUS primarily promotes RNA engagement.

**FIGURE 3.**
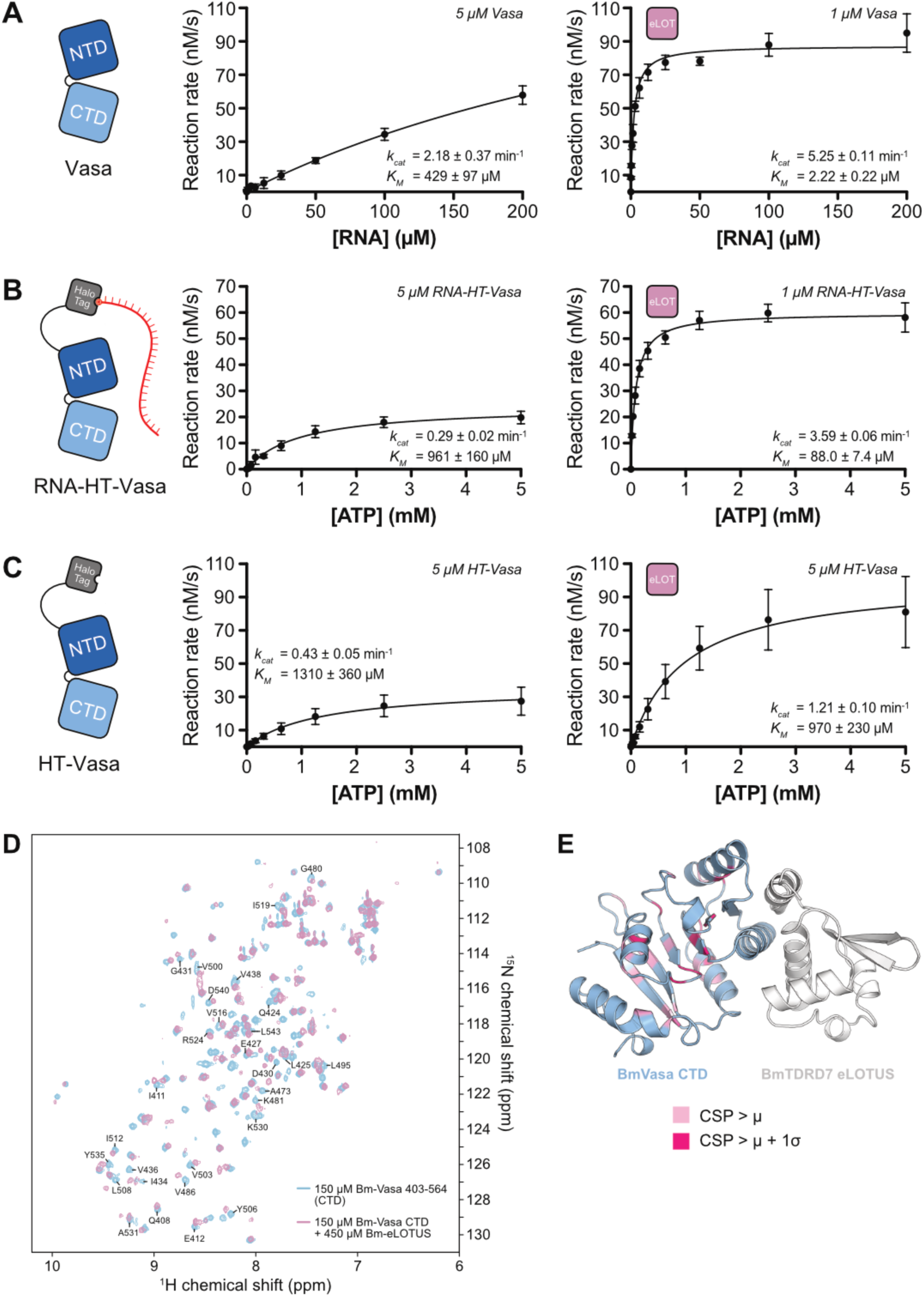
eLOTUS promotes Vasa activity primarily by enhancing RNA binding. (A) NADH-coupled ATPase assay using His-Vasa 200-661 with concentrations indicated, 5 mM ATP, and increasing concentrations of the R26 ssRNA oligo in the absence (left panel) or presence (right panel) of 100 µM Tejas-eLOTUS. (B) NADH-coupled ATPase assay using HT-Vasa 200-661 covalently linked to R26 ssRNA (RNA-HT-Vasa) with concentrations indicated, and increasing concentrations of ATP in the absence (left panel) or presence (right panel) of 100 µM His-Tejas-eLOTUS. (C) NADH-coupled ATPase assay in the absence of RNA using 5 µM HT-Vasa 200-661 and increasing concentrations of ATP in the absence (left panel) or presence (right panel) of 100 µM His-Tejas-eLOTUS. All NADH-coupled ATPase experiments (A-C) were performed in three replicates. Standard deviations are indicated. (D) Overlay of ^1^H-^15^N HSQC spectra of 150 µM *Bombyx* Vasa-CTD in the absence (light blue) and presence (pink) of 450 µM *Bombyx* TDRD7 eLOTUS domain, showing chemical shift perturbations (CSPs) upon binding. Residues with CSPs above the mean are labeled. See also **Supplementary Figures 4E and 4F**. (E) AlphaFold3-generated structural model of the *Bombyx mori* complex composed of the Vasa-CTD and the TDRD7-eLOTUS domain. The PAE plot is shown in **Supplementary Figure 4H**. Residues exhibiting CSPs above the mean (µ) are shown in light pink, whereas residues with CSPs exceeding the mean plus one standard deviation (μ + 1σ) are shown in dark pink. Residues in the CTD α-helix (aa 441-454) contacting eLOTUS are either not assigned or their peaks disappeared during titration (**Supplementary Figure 4E**) and are therefore not colored.

Next, we sought to examine ATP-dependent kinetics, which requires RNA-saturating conditions. However, the maximal RNA concentration of 200 µM was insufficient to saturate Vasa in the absence of eLOTUS (**Figure 3A**), and higher RNA concentrations caused high viscosity and inhibition of enzymatic turnover (**Supplementary Figure 3B**). To overcome this limitation, we mimicked RNA saturating conditions by covalently linking Vasa to its RNA substrate using the HaloTag approach (Los *et al*, 2008), generating a HaloTag-Vasa protein fusion (HT-Vasa) coupled to a chloroalkylated R26 ssRNA (RNA-HT-Vasa; **Supplementary Figure 3C**). ATP titrations under these RNA-saturating conditions revealed that Tejas-eLOTUS increased the catalytic efficiency of RNA-HT-Vasa by ∼135-fold, with *k_cat_* rising ∼12.3-fold and the apparent K_M_ for ATP decreasing ∼10.9-fold. The similar magnitudes of these eLOTUS-induced effects suggest tight coupling between RNA binding and ATP hydrolysis in the presence of eLOTUS.

To investigate whether eLOTUS can influence Vasa independently of RNA, we measured ATPase activity in the absence of RNA. Tejas-eLOTUS produced only a modest 2.8-fold enhancement of HT-Vasa activity (**Figure 3C**), which was insensitive to RNase treatment (**Supplementary Figures 3D and 3E**), suggesting a minor RNA-independent, potentially allosteric effect.

To probe an allosteric contribution directly, we performed NMR experiments using *Bombyx mori* Vasa (Bm-Vasa), previously characterized by NMR (Codutti *et al*, 2024). Because *Drosophila* Tejas-eLOTUS did not stimulate Bm-Vasa (**Supplementary Figure 4A**), we used the eLOTUS domain of Bm-TDRD7. Bm-eLOTUS efficiently stimulated Bm-Vasa ATPase activity (**Supplementary Figure 4B**), similar to prior observations (Jeske et al., 2017), whereas Bm-eLOTUS MUT (S13E/I81E) containing interface mutations equivalent to Tejas SI did not (**Supplementary Figure 4B**). Notably, Bm-eLOTUS also stimulated *Drosophila* Vasa efficiently (**Supplementary Figures 4C and 4D**), indicating functional conservation. Despite poor eLOTUS sequence conservation (Jeske *et al*, 2017), these data, together with the structural data on the Oskar-Vasa complex (Jeske *et al*, 2017), and the mutation analysis of the Tejas-Vasa interaction (**Figure 1**), suggest a functionally conserved binding interface for the eLOTUS-Vasa complex.

**FIGURE 4.**
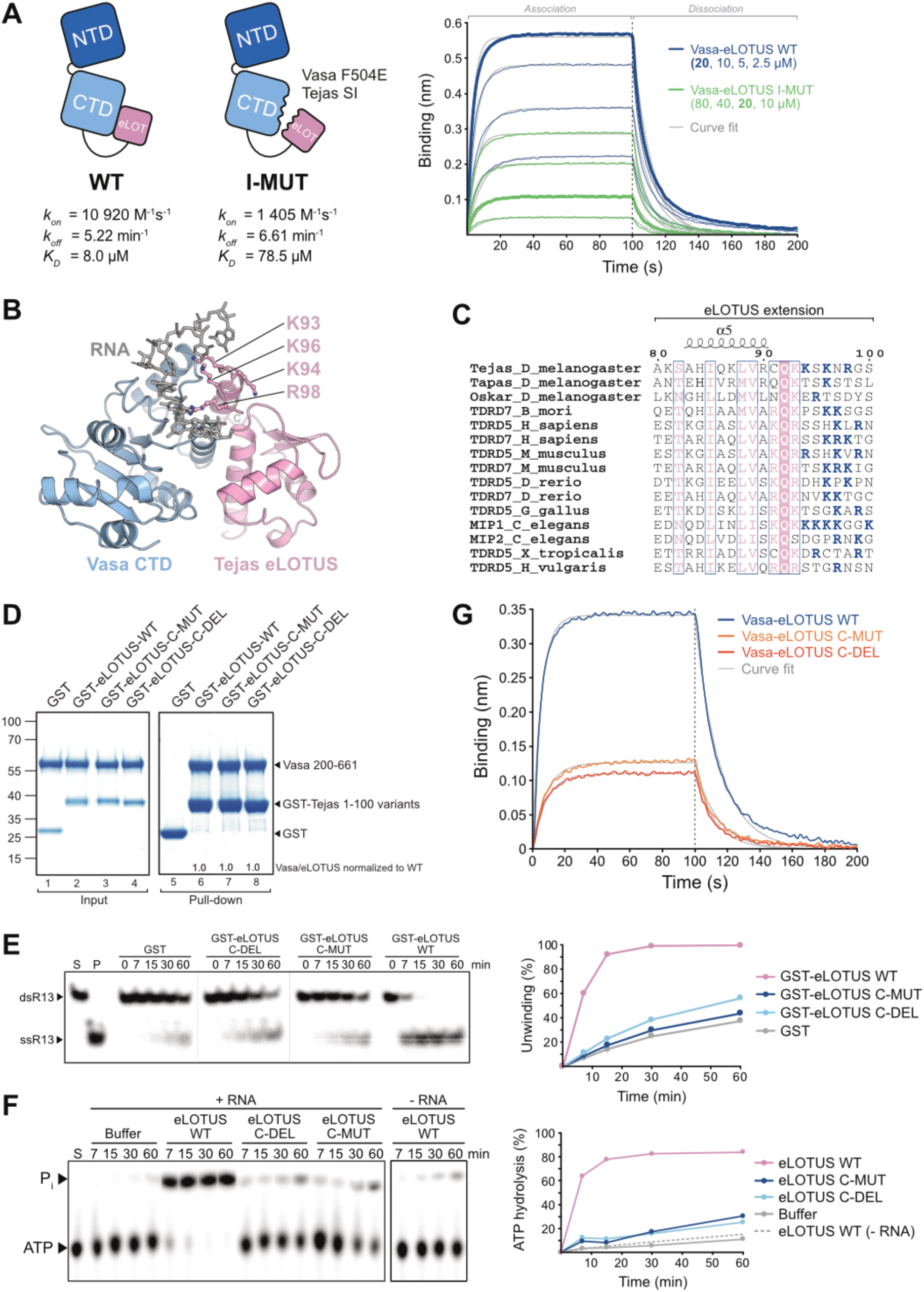
eLOTUS enhances Vasa RNA binding via a basic unstructured tail. (A) Scheme of Vasa core - Tejas-eLOTUS WT and I-MUT fusion proteins (left). Mutations in the Vasa - eLOTUS interface (I-MUT) are indicated. Biolayer interferometry analysis (right) of the fusion proteins was performed in the presence of 2 mM ATP and an ATP-regenerating system. Streptavidin biosensors were loaded in the presence of 5 nM biotinylated R13 ssRNA oligo. For reference, thick lines indicate 20 µM protein concentration. (B) MD-generated structural model of the complex composed of the Vasa-CTD, the Tejas-eLOTUS domain and a short ssRNA oligo. The frame shown corresponds to 6.125 ns. Charged residues in the unstructured C-terminal tail of the eLOTUS domain are highlighted in ball-and-stick representation. (C) Multiple sequence alignment of the C-terminus of the eLOTUS domains from various animals with the positively charged residues in the very C-terminus highlighted in blue. (D) GST pull-down assay using 1 nmol GST, GST-Tejas-eLOTUS WT, C-MUT (K94A/K96A/R98A), or C-DEL (Δ94-100) and 2 nmol His-Vasa 200-661*. Molecular weight marker (in kDa) is indicated at the left. Input and pull-down samples originated from one experiment but were loaded onto two gels. Lanes with irrelevant data were removed. His-Vasa 200-661* contains a short GS linker at its C-terminus not affecting Vasa function. (E) dsRNA unwinding assay in the presence of 3 mM ATP and an ATP-regenerating system, 5 nM labeled dsR13 oligo, and 500 nM unlabeled competitor R13 ssRNA oligo using 7.5 µM of the His-Vasa 200-661* and 100 µM GST or wildtype or mutant GST-Tejas-eLOTUS domain. All samples originated from one experiment but were loaded onto two gels. Lanes with irrelevant mutants were removed. The use of similar amounts of the eLOTUS domain variants was verified by PAGE (**Supplementary Figure 6D**). His-Vasa 200-661* contains a short GS linker at its C-terminus not affecting Vasa function. (F) ATPase assay using 1 µM of His-Vasa 200-661*, 10 µM R26 ssRNA oligo, 8 nM [ψ-^32^P] ATP, in the presence of buffer, 100 µM wildtype or mutant His-Tejas-eLOTUS domains. All samples originated from one experiment but were loaded onto two thin layer plates. The use of similar amounts of the eLOTUS domain variants was verified by PAGE (**Supplementary Figure 6D**). His-Vasa 200-661* contains a short GS linker at its C-terminus not affecting Vasa function. (G) Biolayer interferometry analysis of Vasa-eLOTUS fusion proteins as indicated in the presence of 2 mM ATP and an ATP-regenerating system. Streptavidin biosensors were loaded in the presence of 5 nM biotinylated R13 ssRNA. Association and dissociation curves obtained with 20 µM protein are shown. Full titration series and quantification are provided in **Supplementary Figure 6E**.

NMR titration experiments revealed that Bm-eLOTUS binds exclusively to the CTD of Vasa, with extensive chemical shift perturbations (CSPs) across the domain, suggesting conformational changes within the CTD (**Figures 3D and 3E, Supplementary Figure 4E and 4F**). In contrast, no binding was detected with the NTD (**Supplementary Figure 4G**), which contains the ATP-binding site, suggesting that the minor RNA-independent stimulation likely reflects an indirect allosteric effect of eLOTUS on the Vasa-CTD rather than a direct influence on ATP binding.

Together, these results indicate that the eLOTUS domain primarily promotes RNA engagement by Vasa, thereby stabilizing the catalytically active closed conformation and enhancing ATP hydrolysis, with only a minor RNA-independent allosteric contribution.

### The eLOTUS domain enhances Vasa RNA binding via a basic, unstructured tail

To directly assess the effect of the eLOTUS domain on RNA binding by Vasa, we performed biolayer interferometry (BLI) using the R13 ssRNA oligo. RNA binding by Vasa alone depended on the presence of ATP (**Supplementary Figure 5A**). As adding free eLOTUS to the Vasa-RNA system cannot be interpreted unambiguously in a ternary setup, we created a binary fusion in which the Vasa core was linked to Tejas-eLOTUS (Vasa-eLOTUS WT) via the unstructured C-terminal extension of Vasa (aa 624-661) (**Figure 4A**). As a control, we generated a mutant fusion (Vasa-eLOTUS I-MUT) in which both the Vasa (F504E) and eLOTUS (SI) interfaces were disrupted, preventing complex formation while maintaining identical composition and molecular weight. Importantly, the F504E mutation does not impair helicase activity (Jeske *et al*, 2017). The fusion constructs recapitulated expected performance in ATPase and dsRNA unwinding assays: Vasa-eLOTUS WT reproduced stimulation observed with free eLOTUS, whereas Vasa-eLOTUS I-MUT behaved like Vasa alone (**Supplementary Figures 5B and 5C**).

**FIGURE 5.**
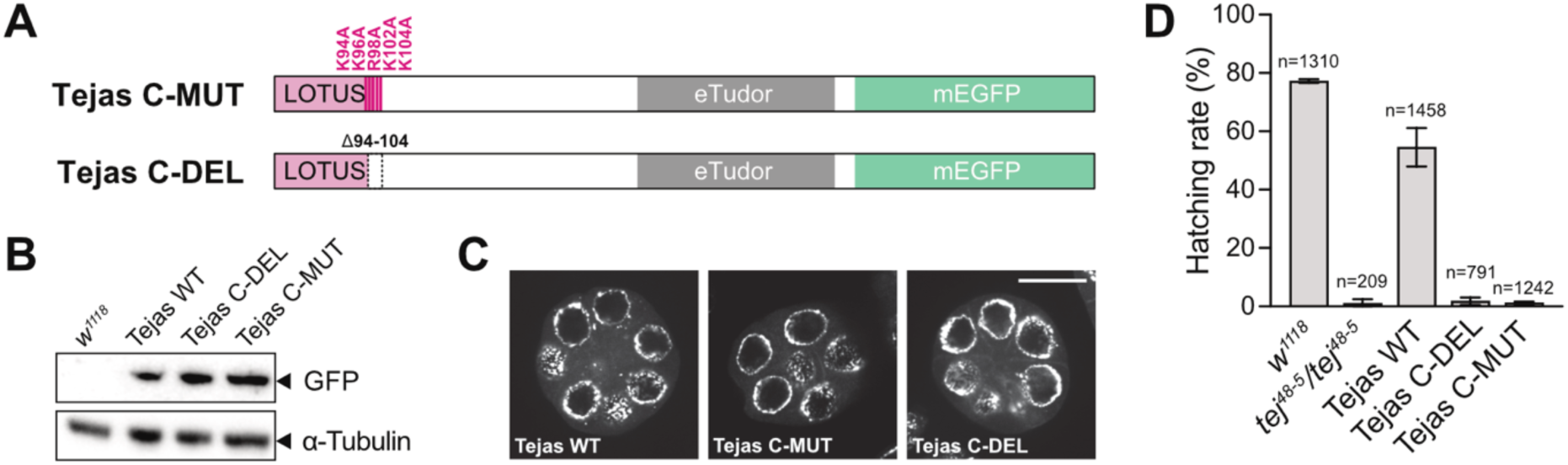
Activation of Vasa is essential *in vivo*. (A) Scheme of the Tejas C-MUT and Tejas C-DEL transgenes. Transgenes were analyzed in the *tej^48-5^/tej^48-5^* null background. (B) Western blot analysis of ovary samples using anti-GFP or anti-α-Tubulin antibodies. (C) Live confocal fluorescence microscopy of stage 3 egg chambers showing the localization of the Tejas-mEGFP transgenes indicated. Scale bar is 20 µm. (D) Hatching rate of eggs laid by females with the genotypes indicated. n indicates the number of eggs scored. Egg laying is shown in **Supplementary Figure 7A**.

BLI measurements in the presence of an ATP-regenerating system revealed that the RNA-binding on-rate of Vasa-eLOTUS WT was approximately eightfold higher than that of Vasa-eLOTUS I-MUT, while the off-rates were similar (**Figure 4A**). The eLOTUS domain also stimulated RNA binding when fused to only the CTD of Vasa (**Supplementary Figure 5D**). Consistently, in EMSAs, eLOTUS enhanced RNA binding by a Vasa variant carrying a mutation at the NTD-CTD interface (Q333A), previously shown to uncouple ATP hydrolysis from dsRNA unwinding (Sengoku *et al*, 2006), suggesting that the mutation perturbs proper domain closure (**Supplementary Figures 5E and 5F**). Together, these results, along with the preference of eLOTUS for the open conformation of Vasa, indicate that eLOTUS primarily facilitates an early step of the helicase cycle by promoting RNA engagement and subsequent closure, without altering the lifetime of the closed state.

Despite this enhancement, the eLOTUS domain alone did not bind either the ssRNA or dsRNA substrates used in this study, even at millimolar concentrations (**Supplementary Figures 5G and 5H**). This indicates that eLOTUS contributes to RNA binding only when complexed with Vasa, rather than through high intrinsic RNA affinity.

To identify residues responsible for this effect, we performed molecular dynamics (MD) simulation of the Vasa CTD in complex with Tejas-eLOTUS and a short ssRNA (**Figure 4B**). We also generated a high-confidence (ipTM=0.85) AlphaFold3 model of a ternary complex containing the Vasa core (**Supplementary Figure 6A**). Although the AlphaFold model incorrectly depicts eLOTUS bound to a closed Vasa conformation, both this model and the MD simulation revealed a positively charged candidate RNA-contact region in the C-terminal stretch of eLOTUS (Lys93, Lys94, Lys96, Arg98), positioned toward the RNA. This stretch is unresolved in the RNA-free Oskar-eLOTUS - Vasa-CTD structure (Jeske *et al*, 2017), consistent with high flexibility and no direct Vasa contact, making it a promising candidate for a stimulation-specific function.

Sequence alignment of eLOTUS domains shows that Lys93 is highly conserved as either lysine or arginine (**Figure 4C**). Mutation of the equivalent residue in Oskar-eLOTUS (K236A) affected both Vasa binding and stimulation (**Supplementary Figures 6B and 6C**), preventing clear interpretation. We therefore targeted residues C-terminal to Lys93, in a short stretch not strictly conserved, but enriched in positively charged residues across eLOTUS domains. Deletion (1′94-100; C-DEL) or mutation (K94A/K96A/R98A; C-MUT) of this stretch in Tejas-eLOTUS did not impair Vasa binding in GST pull-down assays (**Figure 4D**), but both dsRNA unwinding and ATPase stimulation were essentially abolished (**Figures 4E and 4F**). Consistently, BLI of Vasa-eLOTUS fusions with these mutations showed decreased RNA on-rates while off-rates remained unchanged (**Figure 4G, Supplementary Figure 6E**). Interestingly, the small RNA-independent effect of the eLOTUS domain on the ATPase activity of Vasa (**Figure 3C**) was also observed with the eLOTUS C-MUT variant (**Supplementary Figure 6F**), indicating that this effect is independent of the C-terminal stretch and again most consistent with an allosteric contribution upon Vasa binding.

**FIGURE 6.**
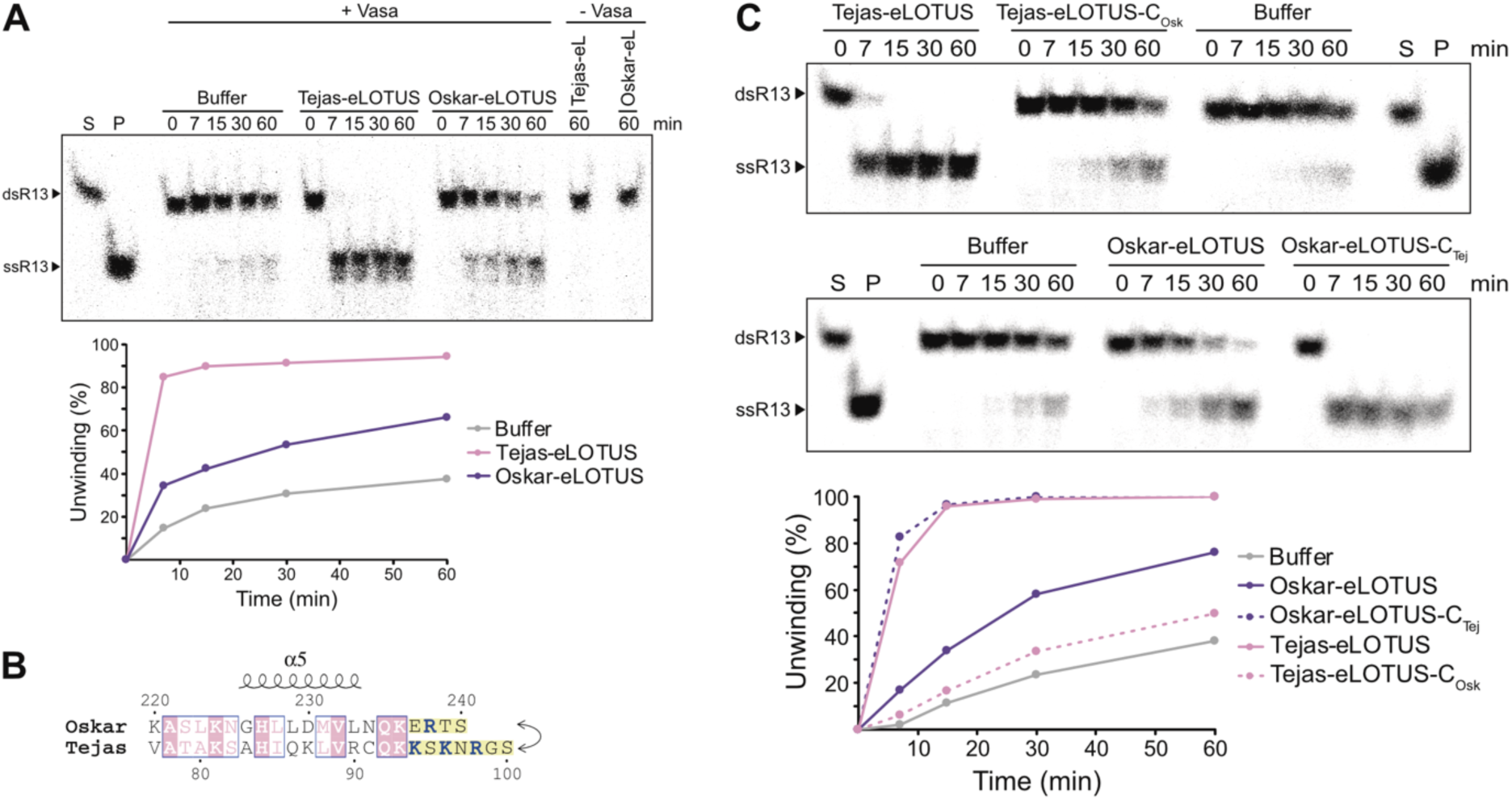
Stimulatory properties of Oskar and Tejas eLOTUS domains can be swapped *in vitro*. (A) dsRNA unwinding assay in the presence of 5 mM ATP and an ATP-regenerating system, 2.5 nM labeled dsR13 oligo, and 500 nM unlabeled competitor R13 ssRNA oligo and using 10 µM of the His-Vasa 200-661* in the absence ("Buffer") or presence of 25 µM of the His-eLOTUS domain of Tejas or Oskar as indicated. His-Vasa 200-661* contains a short GS linker at its C-terminus not affecting Vasa function. (B) Scheme indicating the C-terminal ends that were swapped between the eLOTUS domains of Oskar and Tejas. (C) dsRNA unwinding assay in the presence of 5 mM ATP and an ATP-regenerating system, 2.5 nM labeled dsR13 oligo, and 500 nM unlabeled competitor R13 ssRNA oligo and using 10 µM of the His-Vasa 200-661* in the absence ("Buffer") or presence of 25 µM of the His-eLOTUS domain variant as indicated. The use of similar amounts of the eLOTUS domain variants was verified by PAGE (**Supplementary Figure 7B**). His-Vasa 200-661* contains a short GS linker at its C-terminus not affecting Vasa function.

### eLOTUS-mediated Vasa activation is essential *in vivo*

We showed that the Tejas-eLOTUS - Vasa interaction is required for *in vivo* function (**Figure 1**). To test whether the stimulation of Vasa activity by Tejas-eLOTUS is required *in vivo*, we introduced the corresponding C-DEL or C-MUT mutations into mEGFP-tagged Tejas transgenes (**Figure 5A**). To avoid any potential compensation of a defect, the C-MUT transgene additionally contained two point mutations of positively charged residues extending beyond the structured eLOTUS domain into the disordered region (K102A/K104A) and the C-DEL transgenes a slightly longer deletion (1′94-104). Both Tejas C-DEL and Tejas C-MUT were expressed in ovaries at levels comparable to Tejas WT (**Figure 5B**) and correctly localized to nuage (**Figure 5C**). Strikingly, however, neither mutant rescued the *tejas* null phenotype, as eggs laid by females expressing C-DEL or C-MUT failed to hatch (**Figure 5D, Supplementary Figure 7A**). These data demonstrate that eLOTUS-mediated Vasa activation is critical for germline function *in vivo*.

### Stimulatory properties of Tejas and Oskar eLOTUS can be swapped

Although eLOTUS domain sequences from different proteins and organisms are poorly conserved, binding and stimulation of Vasa by eLOTUS domains appears widespread (Jeske *et al*, 2017). Interestingly, in ATPase (Jeske *et al*, 2017) and dsRNA unwinding assays (**Figure 6A**), Vasa is less efficiently stimulated by Oskar-eLOTUS than by Tejas-eLOTUS, despite similar affinities for the Vasa-CTD. Given that the C-terminal stretch is required for stimulation, we compared this region between Tejas and Oskar eLOTUS and found that Tejas-eLOTUS contains more positively charged residues than Oskar-eLOTUS (**Figure 4C**).

To directly test whether this difference explains the disparity in activity, we generated chimeric eLOTUS domains in which the C-terminal stretch was swapped: Tejas-eLOTUS containing the C-terminal stretch of Oskar (Tejas-eLOTUS-C_Osk_) and Oskar-eLOTUS containing the corresponding region from Tejas (Oskar-eLOTUS-C_Tej_) (**Figure 6B**). Indeed, Tejas-eLOTUS-C_Osk_ displayed strongly reduced Vasa stimulation, whereas Oskar-eLOTUS-C_Tej_ gained stimulatory activity relative to its Tejas wild-type counterpart (**Figure 6C**).

Together, these results demonstrate that the Vasa-stimulatory capacity of eLOTUS relies on the sequence of this short, basic, unstructured C-terminal stretch, which accelerates RNA association without directly contributing to Vasa binding.

### Oskar and Tejas eLOTUS domains are functionally non-interchangeable *in vivo*

The *in vitro* experiments revealed that the eLOTUS domains of Tejas and Oskar differ in their capacity to stimulate Vasa, despite comparable binding to the helicase. These differences in stimulatory potential suggest that the eLOTUS domains might not be functionally interchangeable *in vivo*. To test this, we generated swap transgenes: a genomic *oskar* transgene carrying the Tejas eLOTUS domain (Oskar-TL) and a *tejas* transgene carrying the Oskar eLOTUS domain (Tejas-OL), expressed in their respective null backgrounds (**Figure 7A and Supplementary Figure 7C**). Oskar-WT and Tejas-WT transgenes served as controls.

**FIGURE 7.**
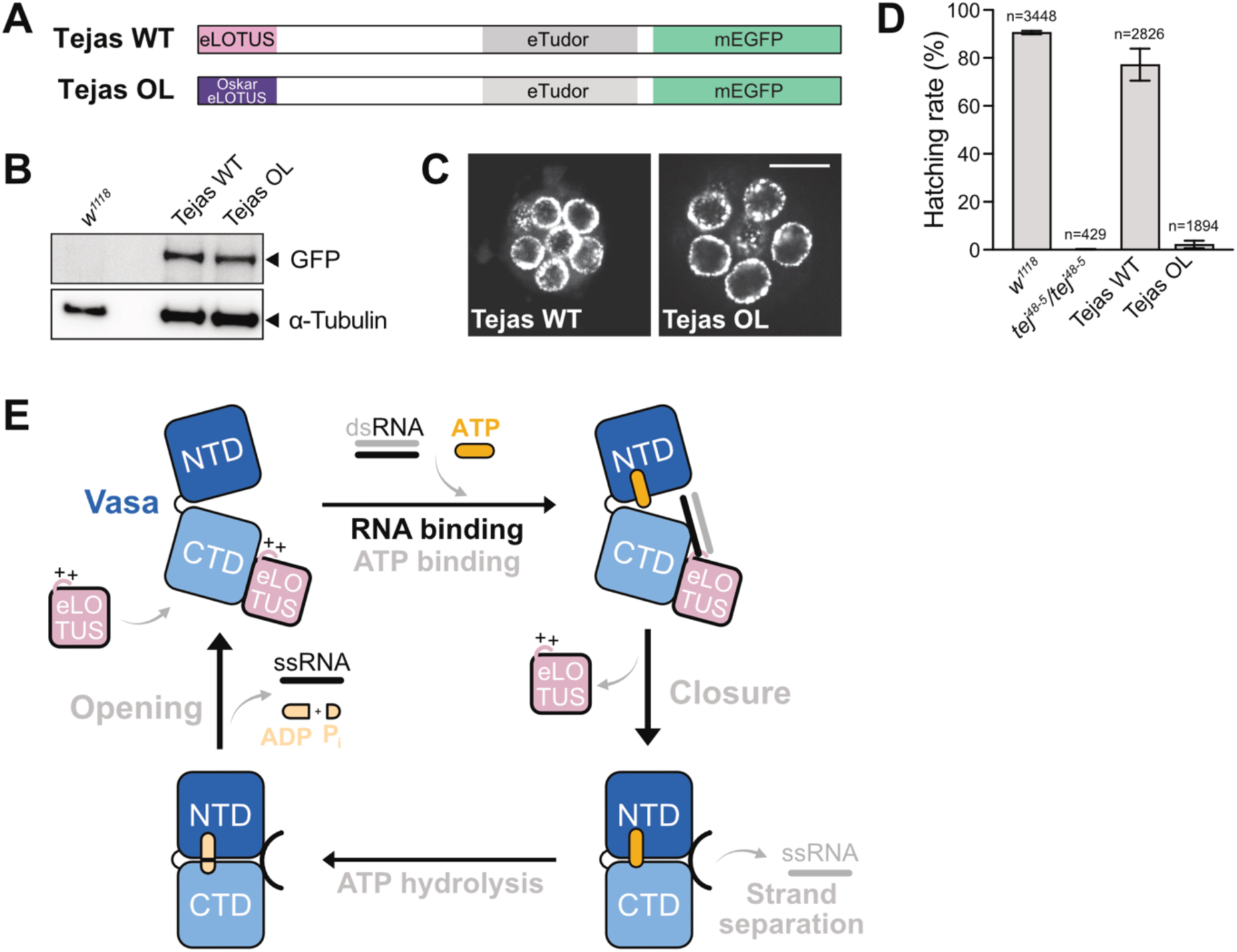
Oskar and Tejas eLOTUS domains are functionally non-interchangeable *in vivo*. (A) Scheme of the Tejas-WT and Tejas-OL transgenes. Transgenes were analyzed in the *tej^48-5^/tej^48-5^* null background. (B) Western blot analysis of ovary samples using anti-GFP or anti-α-Tubulin antibodies. (C) Live confocal fluorescence microscopy of stage 3 egg chambers showing the localization of the Tejas-mEGFP transgenes indicated. Scale bar is 25 µm. (D) Hatching rates of eggs laid by females with the genotype as indicated. n indicates the number of eggs scored. Egg laying is shown in **Supplementary Figure 7F**. (E) Model for the enzymatic cycle of Vasa and its stimulation by eLOTUS domains. The Vasa helicase core interacts with the eLOTUS domain in the open conformation. In this state, Vasa binds RNA and ATP cooperatively. While bound to Vasa through its structured domain, eLOTUS contacts the Vasa-bound RNA strand via its C-terminal unstructured tail, promoting RNA association with the helicase core. Substrate binding induces closure of the helicase core and formation of the catalytically competent conformation. Closure leads to eLOTUS dissociation. In case of dsRNA, core closure also leads to local strand separation. ATP hydrolysis and product release result in reopening of the helicase core, dissociation of the bound RNA, and rebinding of eLOTUS.

Expression analysis revealed that Tejas-OL accumulated at levels comparable to Tejas-WT in ovaries and localized correctly to nuage (**Figures 7B and 7C**), whereas Oskar-TL was undetectable and thus did not rescue, indicating poor protein stability *in vivo* (**Supplementary Figures 7D and 7E**). Despite proper expression and localization, Tejas-OL failed to rescue the hatching defects of *tej^48-5^/tej^48-5^* null embryos, although it partially restored egg-laying, reminiscent to the Tejas C-DEL and C-MUT mutants (**Figure 7D, Supplementary Figure 7F**).

Taken together, these results demonstrate that the eLOTUS domains of Oskar and Tejas are not functionally interchangeable *in vivo*. This specificity is likely influenced by differences in their capacity to stimulate Vasa, but may also reflect the distinct oligomeric properties or additional, context-dependent eLOTUS interactions.

## DISCUSSION

Here we demonstrate the first kinetic characterization of the DEAD-box protein Vasa and dissect the molecular mechanism by which it is activated by eLOTUS domains. Despite the presence of RNA and ATP, the basal ATPase activity of Vasa alone is extremely low and with k_cat_/K_M_ in the range of 4 to 84 M⁻¹ s⁻¹ several orders of magnitude lower than for other characterized DEAD-box proteins (**Supplementary Table 1**). At a physiologically relevant RNA concentrations of 1 µM, a single Vasa molecule would hydrolyze only 4 to 84 x 10^-6^ ATP molecules per second, indicating that the enzyme is essentially inactive. This low intrinsic activity highlights the need for activation by accessory proteins, and the only known regulator of Vasa is the eLOTUS domain.

Based on our mechanistic analyses, we propose a model by which the eLOTUS domain acts on an early step of the catalytic cycle of Vasa (**Figure 7E**). The structured portion of the eLOTUS domain anchors to the CTD of Vasa, while a positively charged, intrinsically disordered region is positioned near the RNA binding site. This basic segment likely engages the RNA backbone through electrostatic interactions, increasing the lifetime of encounter complexes and thereby accelerating RNA association. By promoting rapid RNA engagement, eLOTUS increases the probability that Vasa transitions into its closed, catalytically competent conformation. Formation of the closed, RNA-bound state induces bending of the RNA and, for double-stranded substrates, local strand separation. Transition to the closed state is accompanied by dissociation of the eLOTUS domain, likely due to conformational changes in the CTD that further weaken its already low intrinsic affinity for eLOTUS. Thus, the eLOTUS domain does not stabilize the RNA-bound complex but instead promotes RNA engagement, thereby enhancing the apparent RNA affinity of Vasa.

The mechanism of eLOTUS-mediated activation is fundamentally different from previously described DEAD-box protein activators. Most known activators bind both RecA-like domains of the helicase core (e.g., MIF4G domains (Schütz *et al*, 2008), Barentz (Andersen *et al*, 2006; Bono *et al*, 2006), eIF4H (Marintchev *et al*, 2009; Sun *et al*, 2014)), and typically stimulate ATP binding as well as the often rate-limiting product (P_i_) release (e.g., MIF4G (Hilbert *et al*, 2011; Marintchev *et al*, 2009; Gray *et al*, 2022)) or ATP binding alone (e.g., Esf2 (Granneman *et al*, 2006)) (**Supplementary Table 2**). In contrast, eLOTUS domains act primarily by accelerating RNA association to the helicase, without substantially altering ATP binding or product release. This represents a fundamentally different regulatory strategy: rather than modulating the catalytic steps directly, eLOTUS domains prime Vasa for RNA engagement, facilitating formation of the catalytically competent closed state.

Positively charged disordered regions within diverse RNA- and DNA binding proteins are known to influence nucleic acid binding and folding, for example in RNA processing, DNA repair, and transcription (Vuzman & Levy, 2011; Rajkowitsch *et al*, 2007; Dunker *et al*, 2002; Pontius, 1993; Doetsch *et al*, 2011). The Vasa-eLOTUS complex is reminiscent of the fungal DEAD-box proteins Mss116p and CYT-19. In these proteins, a C-terminal auxiliary domain (CTE) contacts the CTD of the helicase core and is followed by a positively charged disordered tail that binds nonspecifically to structured RNA extensions, thereby increasing RNA binding affinity and tethering the helicase core to unwind nearby RNA helices (Busa *et al*, 2017; Grohman *et al*, 2007; Mohr *et al*, 2008; Mallam *et al*, 2011; Del Campo & Lambowitz, 2009). Despite this compositional similarity, the mechanism of Vasa activation by eLOTUS differs in key aspects. The Mss116p/CYT-19 CTE binds the CTD in the closed helicase core, whereas eLOTUS binds a different interface of the CTD and preferentially the open conformation of Vasa. Furthermore, in Mss116p/CYT-19, the structured CTE contributes strongly to activation, while the unstructured tail has only a minor effect. In contrast, in the Vasa-eLOTUS complex, the structured portion of eLOTUS exerts only a minor allosteric effect, whereas the unstructured C-terminal tail is the main driver of RNA association. Importantly, in Mss116p, deletion of the unstructured tail does not impair *in vivo* function (Del Campo & Lambowitz, 2009), whereas the C-terminal disordered stretch of Tejas-eLOTUS is essential for Vasa activation *in vivo*. These comparisons highlight a unique regulatory mechanism of the eLOTUS-dependent activation of Vasa to control RNA engagement.

Compared to Mss116p/CYT-19, in which the activating component is intrinsic, the Vasa-eLOTUS complex functions as a modular two-component system. This modularity likely enables spatiotemporal control of Vasa activity in the ovary, restricting enzymatic activation to sites where eLOTUS-domain proteins are present. In the ovary, Vasa localization is restricted to the nuage in nurse cells and to the germ plasm at the posterior tip of the oocyte (Liang *et al*, 1994; Hay *et al*, 1988). This localization is critical for Vasa’s biological function and depends on interactions with eLOTUS-domain proteins (Hay *et al*, 1990; Breitwieser *et al*, 1996; Patil *et al*, 2014; Jeske *et al*, 2017). Given Vasa’s low basal catalytic efficiency, unlocalized Vasa lacking eLOTUS contact is essentially inactive and likely not bound to RNA, ensuring that RNA binding, unwinding, or remodeling occur only at the correct subcellular site.

In the *Drosophila* oocyte, Vasa is recruited to the posterior pole through interaction with Oskar (Hay *et al*, 1990; Breitwieser *et al*, 1996). Both proteins are critical for germ cell specification and embryonic patterning (Ephrussi *et al*, 1991; Ephrussi & Lehmann, 1992; Schüpbach & Wieschaus, 1986; Dehghani & Lasko, 2017). They have also been implicated in the translational activation of *nanos* mRNA at the posterior pole (Zaessinger *et al*, 2006; Jeske *et al*, 2011; Kubíková *et al*, 2023; Dahanukar *et al*, 1999), a step critical for posterior embryonic patterning (Wang & Lehmann, 1991; Ephrussi *et al*, 1991; Gavis & Lehmann, 1994). Despite this, the molecular function of Vasa at the posterior remains unclear. An Oskar mutant in the eLOTUS C-terminal tail that uncouples Vasa recruitment from activation could provide a means to directly test the role of Vasa activation in germ plasm function in future studies.

In the context of the piRNA pathway, Vasa is essential for piRNA amplification via the ping-pong cycle (Malone *et al*, 2009). While the precise molecular function of *Drosophila* Vasa in this pathway is not fully understood, studies in *Bombyx mori* suggest that Bm-Vasa is required for the handover of transposon RNA fragments from Siwi to Ago3, protecting the RNA from degradation and enabling proper piRNA amplification (Xiol *et al*, 2014). ATPase activity appears necessary for releasing transposon RNA fragments from Siwi-piRNA complexes after cleavage (Nishida *et al*, 2015). Although no eLOTUS-domain protein has yet been studied in the *Bombyx* piRNA pathway, interaction with an eLOTUS-domain partner may be required to promote RNA engagement. Future studies will be needed to test whether a Vasa-eLOTUS module is critical in the *Bombyx* pathway.

In *Drosophila,* Vasa was proposed to associate with piRNA precursors at the cytoplasmic face of the nuclear pore upon export, promoting their handover to the processing machinery in the nuage (Zhang *et al*, 2012). The core of DEAD-box proteins contacts the sugar-phosphate backbone of a single RNA strand (Sengoku *et al*, 2006; Andersen *et al*, 2006; Bono *et al*, 2006; Collins *et al*, 2009; Del Campo & Lambowitz, 2009; Von Moeller *et al*, 2009), and thus does not inherently recognize specific RNA sequences. Auxiliary domains or interaction partners are therefore necessary for sequence-specific RNA targeting. In the case of Vasa, eLOTUS may contribute to RNA selectivity. While we did not observe binding to the RNA substrates used in this and a previous study (Jeske *et al*, 2015), LOTUS domains have been shown to bind RNA in other contexts. For example, an array of eight mLOTUS domains in mouse MARF1 binds both ssRNA and dsRNA *in vitro* (Yao *et al*, 2018). In addition, the mouse eLOTUS-domain protein TDRD5 preferentially interacts with G-rich RNA *in vivo*, and both mLOTUS and eLOTUS domains, including those of *Drosophila* Tejas and Oskar, recognize RNA G-quadruplexes (G4) *in vitro* (Ding *et al*, 2020). G4 is a stable G-rich secondary structure (Sahayasheela & Sugiyama, 2024), increasingly implicated in the piRNA pathway (Vourekas *et al*, 2015; Zhang *et al*, 2019; Ding *et al*, 2020; Balaratnam *et al*, 2019). AlphaFold3 modeling of G4 complexes with the LOTUS domains from *Drosophila*, mouse, human, plant, and bacterial proteins predicted, a conserved G4-binding surface opposite to the Vasa interface (**Supplementary Figure 8A**). *In vitro* experiments confirmed that the Tejas-eLOTUS domain binds G4, and mutational analysis of predicted interface residues reduced binding ∼4-32-fold (**Supplementary Figures 8B, 8C, 8D**). These data suggest that eLOTUS domains can engage G4 and Vasa simultaneously. Given the enrichment of G4 motifs in *Drosophila* piRNA sequences (Balaratnam *et al*, 2019), Tejas may selectively recognize G-rich piRNA precursors, analogous to the sequence preference observed for its mouse ortholog TDRD5 (Ding *et al*, 2020), and thereby potentially guide Vasa to these RNAs. Importantly, the Tejas G4-binding mutants may provide a tool to dissect the potential role of G4 recognition in *Drosophila* piRNA biogenesis *in vivo* in future studies. Similarly, whether G4 recognition by the Oskar eLOTUS domain contributes to germ plasm function remains to be investigated.

Both germ plasm and nuage are cytosolic granular structures (Chiappetta *et al*, 2022). DEAD-box proteins, including Vasa orthologs from human, worm, and silk moth, have been shown to form condensates depending on their N- and C-terminal unstructured extensions often requiring the presence of RNA and ATP (Nott *et al*, 2015; Chen *et al*, 2020; Hondele *et al*, 2019; Weis & Hondele, 2022; Yamazaki *et al*, 2023). Using *in vitro* condensation assays, we found that *Drosophila* Vasa also forms condensates dependent on its extensions but without added RNA (**Supplementary Figure 9**). The eLOTUS domain of Tejas did not enhance Vasa condensation, but was capable of forming condensates in the presence of RNA. Furthermore, the Vasa core lacking the unstructured extensions, which did not condense alone, was recruited to condensates formed by eLOTUS variants capable of interacting with Vasa. These observations suggest that eLOTUS domains may contribute to nuage and germ granule formation through protein-RNA scaffolding, rather than by directly promoting Vasa self-condensation.

In summary, our study reveals a unique mechanism of DEAD-box regulation in which the eLOTUS domain acts as a modular, RNA-engagement accelerator for Vasa that couples enzymatic activation to subcellular localization.

## METHODS

### DNA constructs

Detailed information on all plasmids used in this study is provided in **Supplementary Table 3**. For cloning, T4 DNA ligation reactions were assembled in 10 µl volumes containing T4 DNA ligase (Thermo Fisher Scientific), 50 ng of vector DNA, and a 5- to 20-fold molar excess of PCR-amplified DNA insert. Reactions were incubated for 1 - 2 h at room temperature. To prevent vector self-ligation, 0.25 µl of the same blunt-end restriction enzyme also used for vector linearization was added to the ligation reaction, unless the restriction site was also present in the insert. Positive clones were screened by colony PCR using one primer annealing to the vector and a second primer specific to the insert. Constructs generated by Gibson assembly were prepared using the Gibson Assembly® Master Mix (New England Biolabs) according to the manufacturer’s instructions, with reaction volumes reduced to half. All plasmids were verified by Sanger sequencing.

### Fly strains and generation of transgenes and transgenic fly lines

The *w^1118^* (Hazelrigg *et al*, 1984), *osk*-GAL4 (Telley *et al*, 2012), and *tej^48-5^* (Patil & Kai, 2010) fly lines were described before. The Tejas WT subclone (pBSK-Tejas-WT) was assembled from four DNA fragments that were ligated in a single reaction. The first fragment comprising the 5’ UTR and the coding sequence of Tejas was amplified from *Drosophila* ovary cDNA introducing 5’ XhoI and 3’ KpnI sites, the second fragment consisting of a linker and the mEGFP sequence was amplified from the Blunt II-TOPO-g-osk mEGFP vector (RR116) introducing 5’ KpnI and 3’ NotI sites, and the third fragment coding for the *tejas* 3’UTR was amplified from *Drosophila* ovary cDNA introducing 5’ NotI and 3’ XhoI sites. All PCR products were double-digested with the respective restriction enzymes and ligated into the XhoI-digested pBluescript SK (pBSK). To obtain the Tejas-SI subclone (pBSK-Tejas-SI), the pBSK-Tejas-WT vector was subjected to two successive site-directed mutagenesis reactions introducing the S16E/I85E double mutation. To generate the pBSK-Tejas-OL subclone, the pBSK-Tejas-WT vector was opened by PCR amplification omitting the Tejas-LOTUS domain (aa 1-100). The Oskar-eLOTUS domain (aa 139-240) sequence was amplified from pAc5.1-mCherry-Short Oskar (Jeske *et al*, 2017) and then inserted via blunt end ligation into the opened pBSK-Tejas-WT vector. All pBSK-Tejas subclones were digested with AgeI and BgIII and ligated into the AgeI and BamHI double digested pUAS-attB-g-*osk* plasmid (K1-MJ; digestion removes the genomic *oskar* region) to obtain attB-pUAS-Tejas WT, attB-pUAS-Tejas SI, and attB-pUAS-Tejas OL transgenes.

To generate attB-pUAS-Tejas C-MUT, a fragment encoding Tejas 71-104 containing the mutated eLOTUS C-terminus was amplified from pMJ-His-Tejas 1-100 C-MUT (ANG6) using a forward primer with an overhang matching the region upstream of the PstI site in pBSK-Tejas-WT (RR239) and a reverse primer overlapping a second Tejas fragment (aa 99-267). This second fragment was amplified from pBSK-Tejas-WT (RR239) using a forward primer overlapping the mutated 71-104 fragment and a reverse primer with an overhang corresponding to the EcoRI site in attB-pUAS-Tejas-WT (RR108). The attB-pUAS-Tejas-WT vector (RR108) was digested with PstI and EcoRI, and both PCR fragments were assembled into the vector by Gibson assembly.

To generate attB-pUAS-Tejas-C-DEL, a pAc5.1-mCherry Tejas Δ94-104 subclone was generated from pAc5.1-mCherry Tejas (H33-MJ) by deletion PCR. Next, a fragment covering Tejas 71-267 was amplified from this pAc5.1-mCherry Tejas Δ94-104 vector using primers matching the flanking regions of attB-pUAS-Tejas-WT (RR108) digested with PstI and EcoRI. The PCR product was then inserted into the digested attB-pUAS-Tejas-WT vector using Gibson assembly.

The attB-pUAS-Tejas transgenes were integrated into the VK33 fly strain at position 65B2 (3R) using ΦC31 integrase using the *Drosophila* injection service of the University of Cambridge (Tejas WT and Tejas SI) or Bestgene Inc. (USA) (Tejas WT, Tejas OL, Tejas C-MUT, and Tejas C-DEL).

### Egg laying and hatching assays

Five to ten virgin females were mated with three *w^1118^* males for 24 h at 25°C to allow egg laying. The flies were then transferred to apple juice agar plates (22.5g/L agar (Millipore), 25% (v/v) apple juice, 2,5% (w/v) sucrose (Roth), 0.15% (w/v) nipagin (Sigma Aldrich)), for egg laying, with the plates being replaced periodically over approximately five days. Each Plate was kept at 25°C for another 24 h to allow hatching. Then, the number of unhatched eggs and empty egg shells was counted to determine egg laying and hatching rates. Experiments were repeated in three (Tejas-TG) or two (Oskar-TG) independent replicates.

### Ovary imaging

For the Tejas-mEGFP protein fluorescence, ovaries from two-days-old flies mated with *w^1118^* males were dissected in PBS and placed and spread on cover glass. Live images were taken immediately using 40 x objective at a Nikon ECLIPSE VoxTi3 confocal microscope or a Nikon Plan Apo λ 60x Oil NA 1.4 objective and a Nikon Ti2-W1 spinning disk confocal fluorescence microscope. The images were processed with Fiji (Schindelin *et al*, 2012).

### Protein extraction from fly ovaries and western blot analysis

Twenty pairs of ovaries from 2-days-old transgenic flies mated with *w^1118^* males were homogenized using a pestle in 200 µl ice-cold NB lysis buffer (Prudêncio & Guilgur, 2015) containing 150 mM NaCl, 50 mM Tris-HCl (pH 7.5), 2 mM EDTA, and 0.1% NP-40. Lysates were clarified by three consecutive centrifugation steps at 20,000 × g for 3 min at 4°C. The supernatant was mixed 1:1 with 50 µl 5 x SDS loading buffer, heated at 95°C for 5 min, and 10 µl was loaded onto a 4-12% NuPAGE™ Bis-Tris gel (Thermo Fisher Scientific). The proteins were transferred semi-dry using Towbin buffer (25 mM Tris-HCl pH 7.5, 192 mM glycine, 20% methanol) on nitrocellulose membrane (GE Healthcare). The membrane was stained with 0.5% (w/v) Ponceau S (Roth) in 1% acetic acid, followed by several washes with water. The membrane was blocked in TN-Tween buffer (20 mM Tris-HCl pH 7.5, 150 mM NaCl, 0.05 % Tween) supplemented with 5% milk powder (Roth) for 30 min and then incubated with mouse anti-GFP (1:1000; Roche, #11814460001), mouse anti-Tubulin (1:50000; Sigma Aldrich, #T5168), rabbit anti-Oskar (1:2000) or mouse anti-Actin (1:1000; abcam, #ab11003) first antibodies in blocking solution overnight at 4°C. After several washes with TN-Tween, the membrane was incubated with horseradish peroxidase (HRP)-coupled goat anti-mouse (1:5000; Invitrogen, #62-6520) or monoclonal mouse anti-rabbit IgG light chain (1:10000; abcam, #ab99697) secondary antibodies for 2 h at RT. The membrane was washed several times with TN-Tween followed by signal detection using ECL Western blotting substrate (Thermo Fisher Scientific). Images were taken with FujiFilm FLA-7000 (Oskar-transgene) and GE AmershamTM Imager 600 (Tejas transgene) and processed with Fiji (Schindelin *et al*, 2012).

### Protein expression and purification

Details about all purified proteins used in this study are provided in **Supplementary Table 4**. *Drosophila* protein expression and purification were performed as described previously (Kubíková *et al*, 2023). For expression, Rosetta 2 competent cells (Novagen) were transformed with the respective protein expression vector and cultured at 37°C in LB medium supplemented with appropriate antibiotics. On the following day, 600 mL of TB medium were supplemented with antibiotics, 2 mM MgSO_4_, solution M (1.25 M Na_2_HPO_4_-7H_2_O, 1.25 M KH_2_PO_4_, 2.5 M NH_4_Cl and 0.25 M Na_2_SO_4_) at a 1:50 ratio, trace elements solution (50 mM FeCl_3_, 20 mM CaCl_2_, 10 mM MnCl_2_, 10 mM ZnSO_4_, 2 mM CoCl_2_, 2 mM CuCl_2_, 2 mM NiCl_2_, 2 mM Na_2_MoO_4_, 2 mM Na_2_SeO_3_, and 2 mM H_3_BO_3_) at a 1:10000 ratio, and 2% lactose, and inoculated with 25 mL of the overnight preculture. Cells were grown at 23°C for 26 h or until an OD_600_ of at least 10 was reached, and then harvested by centrifugation at 4°C and 3500 x g for 10 min.

Protein purification of *Drosophila* proteins was performed by affinity chromatography, using either Ni^2+^-NTA or glutathione agarose, followed by size exclusion chromatography. For glutathione affinity purification of GST-tagged proteins, cells were lysed in a buffer containing 20 mM Tris-Cl (pH 7.5), 500 mM NaCl, 1 mM MgCl_2_, 20% (w/v) glycerol, and 2 mM β-mercaptoethanol supplemented with 10 μg/mL DNase I (PanReac AppliChem) and ½ tablet of SIGMAFAST™ Protease inhibitor Cocktail (Sigma Aldrich). The cleared lysate was incubated with 1 mL glutathione agarose (Pierce®) pre-equilibrated with lysis buffer for 2 h at 4°C. After washing with lysis buffer, proteins were eluted using a buffer containing 100 mM Tris-Cl (pH 8.0), 500 mM NaCl, 20 mM reduced glutathione, and 20% (w/v) glycerol. For Ni^2+^-NTA affinity chromatography of His-tagged proteins, a similar procedure was used with the following modifications: the lysis buffer contained 20 mM Tris-Cl (pH 7.5), 300 mM NaCl, 20 mM imidazole (pH 7), 1 mM MgCl_2_, 10% (w/v) glycerol, and 2 mM β-mercaptoethanol, the elution buffer contained 20 mM Tris-Cl, (pH 7.5), 150 mM NaCl, 250 mM imidazole (pH 7), and 10% (w/v) glycerol, and the cleared lysate was incubated for 30 - 60 min at 4°C with 1 mL HisPur™ Ni-NTA Resin (Thermo Fisher Scientific).

After affinity chromatography, proteins were further purified by ion-exchange chromatography. GST, GST- and His-tagged Oskar-eLOTUS variants, GST-Tejas-eLOTUS variants, and Vasa-eLOTUS fusions were applied to a Q column in 20 mM Tris-Cl (pH 9), 50 mM NaCl, and 10% (w/v) glycerol and eluted using a gradient up to 1 M NaCl. His-tagged Tejas-eLOTUS and Bm-TDRD7-eLOTUS variants were applied to an SP column in 20 mM MES-NaOH (pH 6), 50 mM NaCl, and 10% glycerol, with elution using a gradient up to 1 M NaCl. His-Vasa constructs applied on a heparin column in 20 mM Tris-Cl (pH 7.5), 50 mM NaCl, and 10% glycerol, and eluted using a gradient up to 1 M NaCl.

After concentration, all protein samples were further purified by size-exclusion chromatography. Depending on the molecular weight of the protein, samples were loaded onto either HiLoad 16/600 Superdex 75 pg or 200 pg column (Cytiva), equilibrated in a buffer containing 20 mM Tris-Cl (pH 7.5), 150 mM NaCl, and 10% (w/v) glycerol. Elution fractions containing the protein of interest were concentrated, flash-frozen in liquid nitrogen, and stored at -80°C.

^15^N-labeled Bm-Vasa-NTD (aa 167-397) and CTD (aa 403-564) were expressed and purified as described previously (Codutti *et al*, 2024). Briefly, *E. coli* BL21 RIL cells containing plasmids encoding either domain were grown in minimal medium (M9) supplemented with 1 g/L ^15^N-labeled ammonium chloride (Eurisotop) at 30°C to an OD_600_ of ∼0.8. Protein expression was induced with 1 mM IPTG, and cultures were incubated for an additional 36 h at 11°C before harvesting. Cell pellets were resuspended, treated with lysozyme for 30 min at 4°C, and lysed by sonication on ice. Lysates were clarified by centrifugation at 18,000 rpm for 45 min and filtration through a 0.45 µm membrane. Proteins were purified by Ni^2+^ NTA affinity purification, and the His_6_-tag removed by TEV protease digestion. Proteins were further purified by reverse HisTrap purification, anion exchange, and size-exclusion chromatography and stored in SEC buffer (50 mM Tris-HCl pH 7.6, 150 mM NaCl, 2.5 mM β-mercaptoethanol).

Bm-TDRD7-eLOTUS (aa 1-100) for the NMR experiments was expressed in *E. coli* BL21-RIL cells grown at 37°C to an OD_600_ of ∼0.6. Protein expression was induced with 1 mM IPTG and cultures were incubated overnight at 16°C. Cells were harvested and processed as described for the Bm-Vasa domains. Purification followed the same protocol until the initial HisTrap affinity step. Fractions containing the protein of interest were pooled and incubated overnight at room temperature with TEV protease (1:30) in low-salt buffer (50 mM Tris-HCl pH 8.0, 150 mM NaCl, 5 mM β-mercaptoethanol) to remove the His_6_-tag. Cleaved protein was recovered from the flow-through of a reverse HisTrap purification and buffer exchanged using a HiPrep desalting column (Sephadex G-25, Cytiva) into cation exchange buffer (20 mM MES pH 6.0, 50 mM NaCl, 10% glycerol, 5 mM β-mercaptoethanol). The sample was further purified by cation exchange chromatography on a HiTrap SP HP column (Cytiva) and eluted using a linear NaCl gradient up to 1 M. Final purification was performed by size-exclusion chromatography on a HiLoad 16/600 Superdex 75 pg column (Cytiva) in the same buffer used for the Bm-Vasa domains.

### Preparation of Halo-Vasa-RNA fusion

To couple His-Halo-Vasa 200-661 to RNA, a Cu^+^-catalyzed alkyne azide cycloaddition (CuAAC) reaction was performed. Stock solutions of 100 mM Tris(3-hydroxypropyltriazolylmethyl)amine (THPTA) solution (Lumiprobe), 50 mM copper (I) bromide dispersion (Fluorochem), and a 1:10 dilution (corresponding to 323 mM) of a 1-Azido-18-chloro-3,6,9,12-tetraoxaoctadecane (halo-PEG4-azide) emulsion (Iris Biotech) were prepared in a solvent mixture of water, DMSO (>99.5%, Roth) and *ter*-butanol (99%, Grussing GmbH) at a 4:3:1 (v/v/v) ratio and degassed by argon bubbling for 2-3 min. A reaction containing 20 mM THPTA, 20 mM copper (I) bromide, and 10 mM halo-PEG4-azide was assembled with lyophilized single-stranded hexynyl-R26 oligonucleotide ((GCUUUACGGUGCU)_2_; IDT, 100 nmol synthesis scale) to obtain a final RNA concentration of 1 mM and incubated for 2 h at 37°C. The resulting halo-PEG4-RNA was purified by size-exclusion chromatography (SEC) using a Superdex 75 10/300 GL (Cytiva) column on an Äkta Pure system in buffer containing 20 mM Tris-HCl pH 6.8 and 150 mM NaCl at 4°C. Fractions containing halo-PEG4-RNA were concentrated using Amicon-Ultra™ 0.5 mL centrifugal filters (3 kDa MWCO; Millipore). Halo-PEG4-RNA was then coupled to His-Halo-Vasa 220-661 at a 1:1.1 molar ratio for 30 min at room temperature. Prior to coupling, the protein was pre-treated for 10 min RiboLock RNase inhibitor (Thermo Fisher Scientific) at a final concentration of 1U/µL. To separate the RNA-Halo-Vasa 220-661 fusion from the uncoupled protein, the sample was adjusted to pH 9 using buffer containing 20 mM Tris-HCl pH 9, 350 mM NaCl, 5 mM DTT, and 10% glycerol and loaded onto a HiTrap™ Q-XL 1 mL column (Cytiva) connected to an Äkta Pure system. The column was washed overnight with buffer containing 20 mM Tris-HCl pH 9, 350 mM NaCl, 10% glycerol, and the RNA-Halo-Vasa 220-661 fusion was eluted with 20 mM Tris-HCl pH 9, 600 mM NaCl, 10% glycerol. Eluted fractions were diluted to 150 mM NaCl using dilution buffer (Tris-HCl pH 7.5, 10% glycerol), concentrated using 20 mL centrifugal filters (30 kDa MWCO; Pierce™), flash-frozen in liquid nitrogen, and stored at -80°C.

### Protein-protein interaction assays

Isothermal titration calorimetry (ITC) was performed as described previously (Kubíková *et al*, 2023). His-Tejas 1-100 and His-Vasa 463-661 proteins were co-dialyzed in separate dialysis cassettes in a buffer containing 20 mM Tris-HCl pH 7.5 and 150 mM NaCl over night at 4°C. A total volume of 36.4 μL of 490 µM His-Tejas 1-100 was injected into a total volume of 250 μL of 40 µM His-Vasa 463-661. Initially, 0.4 μL were injected, followed by 18 injections in 2 μL steps.

GST pull down assays were performed in 100 µl reactions as described previously (Kubíková *et al*, 2023; Salgania *et al*, 2025). Specifically, proteins indicated in the figure legends were incubated for 30 minutes at 25°C in incubation buffer containing 20 mM Tris-Cl (pH 7.5), 150 mM NaCl, 10% (w/v) glycerol, 5 mM DTT, and 0.1% Tween-20. Subsequently, 40 µL of a 1:2 slurry of glutathione agarose resin (Pierce™, Thermo Fisher Scientific) in incubation buffer was added, and the mixture was incubated for an additional 90 minutes with intermittent mixing. The resin was then washed five times with 1 mL of incubation buffer. Bound proteins were eluted with SDS sample buffer, separated by SDS - PAGE, and visualized using PageBlue Protein Staining Solution. Protein concentrations used are indicated in the figure legends. To induce the closed conformation of His-Vasa 200-623-WT, 20 µM of the protein was preincubated for 5 min with 40 µM of the R13 ssRNA oligo and 2 mM ATP at RT prior to mixing with the GST fusion proteins.

ReLo assays were performed as described previously (Salgania *et al*, 2024). S2R+ cells were cultured at 25°C in Schneider’s *Drosophila* medium+ (L)-glutamine (Thermo Fisher Scientific) supplemented with 10% fetal bovine serum (Sigma Aldrich) and 1 x Gibco™ Antibiotic-Antimycotic (Thermo Fisher Scientific). Cells were seeded into four-well polymer µ-slides (Ibidi) and cotransfected using jetOPTIMUS (Polyplus Transfection) according to the user manual. Specifically, 600 µl of cells at a density of 1 x 10^6^ cells/ml were seeded per well and 61 µl of transfection reagent mixture was added. This mixture contained 1 µl of transfection reagent and a total of 600 ng of DNA for cotransfection of two (300 ng DNA each) plasmids diluted in jetOPTIMUS buffer. After 48 hours of incubation at 25°C, images of live cells were acquired using a Nikon Plan Apoλ 100 × NA 1.45 oil objective and a Nikon Ti2 microscope equipped with a Yokogawa CSU-W1 confocal scanner unit. NIS-Elements AR software was used for image acquisition. Images were processed with Fiji software (Schindelin *et al*, 2012).

### NMR

Two-dimensional ^1^H-^15^N correlation spectra of ^15^N-labeled Bm-Vasa-CTD or NTD were acquired in the absence and presence of increasing concentrations of Bm-TDRD7-eLOTUS (aa 1-100) at 298 K (ca. 25°C) in a buffer containing 50 mM Tris pH 7.5, 150 mM NaCl, and 2.5 mM β-mercaptoethanol. The initial concentration of Vasa-CTD or - NTD was 150 µM. All titration points were recorded on a 900 MHz Bruker Avance III HD spectrometer using 3 mm NMR tubes and a standard HSQC pulse sequence (Bodenhausen & Ruben, 1980). In the indirect ^15^N dimension, the carrier frequency was set to 91.25 MHz, the spectral width was 25 ppm (2281 Hz), and 180 increments were collected. Data were processed using NMRPipe (Delaglio *et al*, 1995), and phase corrections were applied manually in both dimensions to generate the final HSQC spectra.

### RNA-protein interaction assay

For biolayer interferometry (BLI) analysis, Octet^®^ SA (Streptavidin) Biosensors (Sartorius) were pre-incubated for at least 10 minutes in reaction buffer containing 20 mM Tris (pH 7.5), 150 mM NaCl, 5 mM MgCl_2_, 5 mM DTT, 2 mM ATP, 20 mM creatine phosphate (Roche), 80 ng/mL creatine kinase (Roche), 0.02% Tween 20 (Roth), and 50 ng/µL heparin (sodium salt from porcine intestinal mucosa, Sigma-Aldrich). The assay was carried out in a black, non-binding F-bottom 96-well plate (Greiner, #655900) using the Octet® RED96e System (Sartorius) with a total volume of 200 µL per well. Following an initial baseline measurement in reaction buffer, biosensors were loaded with a 5’-biotinylated single-stranded R13 oligonucleotide (GCUUUACGGUGU) at a concentration indicated in the figure legends for 300 seconds. Association with Vasa was measured by transferring the loaded biosensors into wells containing reaction buffer supplemented with different concentrations of His-Vasa 200-661* or His-Vasa-LOTUS fusion proteins as indicated for 100 seconds. Dissociation was monitored for 200 s by transferring the biosensors to wells containing reaction buffer only. Binding kinetics were analyzed using Octet® Data Analysis Software (Sartorius) with a 1:1 global fitting model.

### ATPase assays

Thin layer chromatography (TLC) assays were performed as described previously (Jeske *et al*, 2017). Proteins were incubated at 23°C with 0.1 µL (corresponding to 8.3 nM) of 10 µCi/µL [γ-^32^P]-ATP (Hartmann Analytic, SRP-301) in a volume of 20 µL in the presence or absence of single-stranded R26 RNA oligo of the sequence (GCUUUACGGUGU)_2_. The protein and RNA concentrations used are indicated in the figure legends. Per time point, 4 µL of the reaction was transferred into 50 µL of 5 mM EDTA (pH 7) to stop the reaction. The mixture was subjected to phenol/chloroform extraction, and 2 µL of the aqueous phase were analyzed by TLC on polyethyleneimine-cellulose (Merck) using 1 M formic acid and 0.5 M LiCl as mobile phase. The TLC plates were analyzed by phosphorimaging, ATP hydrolysis was quantified using Fiji (Schindelin *et al*, 2012).

Steady-state ATPase activity was measured using an NADH-coupled assay, as previously described (Guo and Pyle, 2022). A 5 x "enzyme cocktail" stock solution was freshly prepared containing 100 U/mL lactate dehydrogenase (Sigma Aldrich), 500 U/mL pyruvate kinase (Sigma Aldrich), 2.5 mM phosphoenolpyruvate (Sigma Aldrich), 1 mM NADH (Sigma Aldrich), 7.5 mM MgCl_2_, and filled up with ATPase reaction buffer (25 mM HEPES, pH 7.4, 150 mM NaCl, 5 mM DTT). Reactions were assembled in UV-Star^®^ 96-well half-area microplates (Greiner Bio-One) with a final volume of 25 µL.

For ATPase activity measurements as a function of RNA concentration, the reaction mixture contained 1 x enzyme cocktail, 1 U/µL RiboLock (Thermo Fisher Scientific), His-Vasa 200-661, 100 µM Tejas eLOTUS, and increasing concentrations of single-stranded R26 oligonucleotide ((GCUUUACGGUGCU)_2_). For ATPase activity measurements as a function of ATP concentration, the reactions contained 1 x enzyme cocktail, 1 U/µL RiboLock, and RNA-HT-Vasa or HT-Vasa at concentrations indicated in the figure legends. All reactions were filled up with ATPase reaction buffer and initiated by adding a MgCl_2_-ATP mixture at constant (5 mM) (RNA titration) or increasing (ATP titration) concentrations. The stock solution of the MgCl_2_-ATP mixture contained 50 mM ATP and 50 mM MgCl_2_ and was prepared by mixing equal volumes of 100 mM ATP (Roche; dissolved in 100 mM Tris-Cl pH 7.5) and 100 mM MgCl_2_. Immediately after setting up the reactions, plates were centrifuged at 3,000 rpm for 3 min, and the NADH absorbance at 340 nm was recorded every 30 seconds for at least 30 minutes using a Tecan Spark^®^ multimode microplate reader (the decrease in NADH absorbance corresponds to the rate of ATP hydrolysis). Reaction rates were plotted against substrate concentrations (ATP or RNA), and enzymatic parameters were calculated using GraphPad Prism 9 by fitting the data to the Michaelis-Menten model.

### Radioactive labeling of RNA substrates

All RNA oligos were purchased from IDT. ^32^P 5’ labeling of RNA was carried out in a 50 µL reaction containing 10 µM RNA oligo, 25 U T4 Polynucleotide kinase (Thermo Fisher Scientific), 1x reaction buffer (Thermo Fisher Scientific), and 2.5 µL of 10 µCi/µL [γ-^32^P] ATP (Hartmann Analytic), corresponding to 80 nM, for 1 h at 37°C. The labeled RNA was purified using MicroSpin™ G-25 columns (illustra™).

### dsRNA unwinding assays

To generate the dsRNA substrate for the unwinding assays, 250 nM ^32^P-labeled R13 oligonucleotide (GCUUUACGGUGU (Rogers *et al*, 2001)) and unlabeled complementary R13C oligonucleotide (AGCACCGUAAAGC) were mixed in a 1:1.2 molar ratio in a buffer containing 200 mM Tris-HCl (pH 7.5), 100 mM MgCl_2_, and 250 mM NaCl. The mixture was incubated at 95°C for 5 min and then slowly cooled to room temperature over approximately 90 min by switching off the heating block.

For the dsRNA unwinding assays, His-Vasa 200-661 and varying concentrations of eLOTUS were mixed in a 100 µL reaction volume containing 20 mM Tris-HCl (pH 7.5), 1 mM DTT, 80 ng/µL creatine kinase (Roche), 20 mM creatine phosphate (Roche), 1 U/µL RiboLock RNase inhibitor (Thermo Fisher Scientific), 5 mM MgCl_2_-ATP mixture, and 2.5 nM ^32^P-labeled R13 dsRNA. The protein concentrations used are indicated in the figure legends. The reaction was incubated on ice for 5 min, then 500 nM unlabeled competitor R13 ssRNA was added. The reaction was carried out at 25°C, and at the indicated time points, 20 µL aliquots were taken and mixed with 14 µL stop solution (140 mM Tris-HCl (pH 8.0), 70 mM EDTA, 1.4% SDS, 0.02% bromophenol blue, 0.02% xylene cyanol, 28% glycerol, and 0.6 mg/mL Proteinase K) and incubated for 15 min at 25°C. 20 µl of the samples were loaded onto a 20% native polyacrylamide gel (37.5:1; 0.5 x TBE) and run at 260 V for 3 hours at room temperature. The gel was then dried and exposed overnight on a phosphorimaging screen, which was subsequently scanned using a FujiFilm FLA-7000 or an Azure Sapphire Biomolecular Imager. dsRNA unwinding was quantified using Fiji (Schindelin *et al*, 2012).

### Structural modeling

For molecular dynamics (MD) simulations, a complex comprising Tejas-eLOTUS (aa 1-99), ssRNA (CGGGCCCG), and the Vasa core (aa 202-627) in an open conformation was constructed. To generate the open conformation of Vasa, the crystal structure (PDB ID: 2DB3) was separated into the NTD (including bound AMP-PNP), the CTD, and the interdomain linker (456-464). The linker conformation was replaced with that of the human DDX19 open conformation structure (PDB ID: 3EWS), and residues were subsequently mutated back to the Vasa sequence. The domains and linker were then reassembled and energy-minimized. RNA was positioned by alignment to the Vasa-RNA complex (PDB ID: 2DB3) and placed in contact with the CTD. The resulting Vasa-Tejas-RNA complex was used to initiate MD simulations using the Desmond module of the Schrödinger suite (release 2021-1) (Bowers *et al*, 2006) with the default force field. Systems were prepared using standard protocols, including assignment of protonation states and addition of hydrogen atoms, followed by energy minimization. Complexes were solvated in SPC water with 150 mM NaCl and simulated at 300 K for 25 ns. For clarity, the Vasa NTD, which does not contact RNA or eLOTUS, was omitted from the figure.

For AlphaFold3 (Abramson *et al*, 2024) predictions, the web interface with standard settings was used. The PAE plots were visualized using UCSF ChimeraX (Pettersen *et al*, 2021). All structures were visualized with PyMOL (Schrödinger, LLC. (2015). The PyMOL Molecular Graphics System, Version 3.0).

## ACKNOWLEDGMENTS AND FUNDING SOURCES

We thank Rebecca Reinig, Jana Kubikova, Harpreet Kaur Salgania, Alina Schneider, Marius Müldner, and Annika Klau for technical assistance, and Patricia Giselle Cipriani for discussion. We are grateful to Sherif Ismail and Aleš Obrdlik for constructive feedback on the manuscript. We thank Anne Ephrussi for providing anti-Oskar and anti-Vasa antibodies, Frank Wippich for the *osk*-STOP-cTRICK fly line, and Toshie Kai for the *tejas^48-5^* fly line. We also acknowledge the Fly Facility at the University of Cambridge, UK, for microinjection services, and the Nikon Imaging Center at the University of Heidelberg for access to microscopes. We thank the data storage service SDS@hd, supported by the Ministry of Science, Research and the Arts Baden-Württemberg (MWK) and the German Research Foundation (DFG) through the grants INST 35/1314-1 FUGG and INST 35/1503-1 FUGG. This work was funded by the Emmy Noether Program of the DFG (JE-827/1-1 and JE-827/1-2 to M.J.), by the Chica and Heinz Schaller Foundation (CHS; 2024 Award to M.J.), by the Swiss National Science Foundation (SNF; PCEFP3_187052 to M.H.), by the European Research Council (ERC-2020-STG 950262 to M.H.), and by the Leverhulme Trust (LIP-2020-017 to T.C.). This work was also supported in part through the NYU IT High Performance Computing resources, services, and staff expertise.

## REFERENCES

Abramson J, Adler J, Dunger J, Evans R, Green T, Pritzel A, Ronneberger O, Willmore L, Ballard AJ, Bambrick J, et al (2024) Accurate structure prediction of biomolecular interactions with AlphaFold 3. Nature 2024 630:8016 630: 493–500

Anantharaman V, Zhang D & Aravind L (2010) OST-HTH: A novel predicted RNA-binding domain. Biol Direct 5: 1–8

Andersen CBF, Ballut L, Johansen JS, Chamieh H, Nielsen KH, Oliveira CLP, Pedersen JS, Séraphin B, Hir H Le & Andersen GR (2006) Structure of the exon junction core complex with a trapped DEAD-Box ATPase bound to RNA. Science (1979) 313: 1968–1972

Balaratnam S, Hettiarachchilage M, West N, Piontkivska H & Basu S (2019) A secondary structure within a human piRNA modulates its functionality. Biochimie 157: 72–80

Bodenhausen G & Ruben DJ (1980) Natural abundance nitrogen-15 NMR by enhanced heteronuclear spectroscopy. Chem Phys Lett 69: 185–189

Bohnsack KE, Yi S, Venus S, Jankowsky E & Bohnsack MT (2023) Cellular functions of eukaryotic RNA helicases and their links to human diseases. Nature Reviews Molecular Cell Biology 2023 24:10 24: 749–769

Bono F, Ebert J, Lorentzen E & Conti E (2006) The Crystal Structure of the Exon Junction Complex Reveals How It Maintains a Stable Grip on mRNA. Cell 126: 713–725

Bowers KJ, Chow E, Xu H, Dror RO, Eastwood MP, Gregersen BA, Klepeis JL, Kolossvary I, Moraes MA, Sacerdoti FD, et al (2006) Scalable Algorithms for Molecular Dynamics Simulations on Commodity Clusters. International Conference on Software Composition

Breitwieser W, Markussen FH, Horstmann H & Ephrussi A (1996) Oskar protein interaction with vasa represents an essential step in polar granule assembly. Genes Dev 10: 2179–2188

Busa VF, Rector MJ & Russell R (2017) The DEAD-Box Protein CYT-19 Uses Arginine Residues in Its C-Tail to Tether RNA Substrates. Biochemistry 56: 3571–3578

Callebaut I & Mornon JP (2010) LOTUS, a new domain associated with small RNA pathways in the germline. Bioinformatics 26: 1140–1144

Del Campo M & Lambowitz AM (2009) Structure of the Yeast DEAD Box Protein Mss116p Reveals Two Wedges that Crimp RNA. Mol Cell 35: 598–609

Cao W, Coman MM, Ding S, Henn A, Middleton ER, Bradley MJ, Rhoades E, Hackney DD, Pyle AM & De La Cruz EM (2011) Mechanism of Mss116 ATPase reveals functional diversity of DEAD-Box proteins. J Mol Biol 409: 399–414

Carrera P, Johnstone O, Nakamura A, Casanova J, Jäckle H & Lasko P (2000) VASA mediates translation through interaction with a Drosophila yIF2 homolog. Mol Cell 5: 181–187

Caruthers JM, Johnson ER & McKay DB (2000) Crystal structure of yeast initiation factor 4A, a DEAD-box RNA helicase. Proc Natl Acad Sci U S A 97: 13080–13085

Chen W, Hu Y, Lang CF, Brown JS, Schwabach S, Song X, Zhang Y, Munro E, Bennett K, Zhang D, et al (2020) The dynamics of P granule liquid droplets are regulated by the caenorhabditis elegans germline RNA helicase GLH-1 via its ATP hydrolysis cycle. Genetics 215: 421–434

Chen Y, Boland A, Kuzuoǧlu-Öztürk D, Bawankar P, Loh B, Chang C Te, Weichenrieder O & Izaurralde E (2014) A DDX6-CNOT1 Complex and W-Binding Pockets in CNOT9 Reveal Direct Links between miRNA Target Recognition and Silencing. Mol Cell 54: 737–750

Chen Y, Potratz JP, Tijerina P, Del Campo M, Lambowitz AM & Russell R (2008) DEAD-box proteins can completely separate an RNA duplex using a single ATP. Proc Natl Acad Sci U S A 105: 20203–20208

Chiappetta A, Liao J, Tian S & Trcek T (2022) Structural and functional organization of germ plasm condensates. Biochemical Journal 479: 2477–2495

Cipriani PG, Bay O, Zinno J, Gutwein M, Gan HH, Mayya VK, Chung G, Chen J-X, Fahs H, Guan Y, et al (2021) Novel LOTUS-domain proteins are organizational hubs that recruit C. elegans Vasa to germ granules. Elife 10: 1–34

Codutti L, Kirkpatrick JP, zur Lage S & Carlomagno T (2024) Long-range conformational changes in the nucleotide-bound states of the DEAD-box helicase Vasa. Biophys J 123: 3884–3897

Collins R, Karlberg T, Lehtiö L, Schütz P, van den Berg S, Dahlgren LG, Hammarström M, Weigelt J & Schüler H (2009) The DEXD/H-box RNA Helicase DDX19 Is Regulated by an α-Helical Switch. Journal of Biological Chemistry 284: 10296–10300

Dahanukar A, Walker JA & Wharton RP (1999) Smaug, a novel RNA-binding protein that operates a translational switch in Drosophila. Mol Cell 4: 209–218

Dehghani M & Lasko P (2017) Multiple functions of the DEAD-box helicase vasa in Drosophila oogenesis. Results Probl Cell Differ 63: 127–147

Delaglio F, Grzesiek S, Vuister GW, Zhu G, Pfeifer J & Bax A (1995) NMRPipe: a multidimensional spectral processing system based on UNIX pipes. J Biomol NMR 6: 277–293

Ding D, Wei C, Dong K, Liu J, Stanton A, Xu C, Min J, Hu J & Chen C (2020) LOTUS domain is a novel class of G-rich and G-quadruplex RNA binding domain. Nucleic Acids Res 48: 9262–9272

Doetsch M, Schroeder R & Fürtig B (2011) Transient RNA–protein interactions in RNA folding. FEBS J 278: 1634–1642

Donsbach P & Klostermeier D (2021) Regulation of RNA helicase activity: Principles and examples. Biol Chem 402: 529–559

Dunker AK, Brown CJ, Lawson JD, Iakoucheva LM & Obradović Z (2002) Intrinsic disorder and protein function. Biochemistry 41: 6573–82

Ephrussi A, Dickinson LK & Lehmann R (1991) Oskar Organizes the Germ Plasm and Directs Localization of the Posterior Determinant Nanos. Cell 66: 37–50

Ephrussi A & Lehmann R (1992) Induction of germ cell formation by oskar. Nature 358: 387–392

Fairman-Williams ME, Guenther UP & Jankowsky E (2010) SF1 and SF2 helicases: Family matters. Curr Opin Struct Biol 20: 313–324

Gavis ER & Lehmann R (1994) Translational regulation of nanos by RNA localization. Nature 369: 315–318

Gorbalenya AE, Koonin E V., Donchenko AP & Blinov VM (1988) A conserved NTP-motif in putative helicases. Nature 333: 22

Granneman S, Lin CY, Champion EA, Nandineni MR, Zorca C & Baserga SJ (2006) The nucleolar protein Esf2 interacts directly with the DExD/H box RNA helicase, Dbp8, to stimulate ATP hydrolysis. Nucleic Acids Res 34: 3189–3199

Gray S, Cao W, Montpetit B & De La Cruz EM (2022) The nucleoporin Gle1 activates DEAD-box protein 5 (Dbp5) by promoting ATP binding and accelerating rate limiting phosphate release. Nucleic Acids Res 50: 3998–4011

Grohman JK, Del Campo M, Bhaskaran H, Tijerina P, Lambowitz AM & Russell R (2007) Probing the mechanisms of DEAD-box proteins as general RNA chaperones: The C-terminal domain of CYT-19 mediates general recognition of RNA. Biochemistry 46: 3013–3022

Guo R & Pyle AM (2022) Monitoring functional RNA binding of RNA-dependent ATPase enzymes such as SF2 helicases using RNA dependent ATPase assays: A RIG-I case study. Methods Enzymol 673: 39–52

Handler D, Olivieri D, Novatchkova M, Gruber FS, Meixner K, Mechtler K, Stark A, Sachidanandam R & Brennecke J (2011) A systematic analysis of Drosophila TUDOR domain-containing proteins identifies Vreteno and the Tdrd12 family as essential primary piRNA pathway factors. EMBO Journal 30: 3977–3993

Hay B, Jan LY & Jan YN (1988) A protein component of Drosophila polar granules is encoded by vasa and has extensive sequence similarity to ATP-dependent helicases. Cell 55: 577–587

Hay B, Jan LY & Jan YN (1990) Localization of vasa, a component of Drosophila polar granules, in maternal-effect mutants that alter embryonic anteroposterior polarity. Development 109: 425–433

Hazelrigg T, Levis R & Rubin GM (1984) Transformation of white locus DNA in Drosophila: Dosage compensation, zeste interaction, and position effects. Cell 36: 469–481

Henn A, Cao W, Hackney DD & De La Cruz EM (2008) The ATPase Cycle Mechanism of the DEAD-box rRNA Helicase, DbpA. J Mol Biol 377: 193–205

Hilbert M, Kebbel F, Gubaev A & Klostermeier D (2011) eIF4G stimulates the activity of the DEAD box protein eIF4A by a conformational guidance mechanism. Nucleic Acids Res 39: 2260–2270

Hondele M, Sachdev R, Heinrich S, Wang J, Vallotton P, Fontoura BMA & Weis K (2019) DEAD-box ATPases are global regulators of phase-separated organelles. Nature 573: 144–148

Jankowsky E (2011) RNA helicases at work: Binding and rearranging. Trends Biochem Sci 36: 19–29

Jankowsky E & Fairman ME (2007) RNA helicases - one fold for many functions. Curr Opin Struct Biol 17: 316–324

Jarmoskaite I & Russell R (2014) RNA helicase proteins as chaperones and remodelers. Annu Rev Biochem 83: 697–725

Jeske M, Bordi M, Glatt S, Müller S, Rybin V, Müller CW & Ephrussi A (2015) The crystal structure of the Drosophila germline inducer Oskar identifies two domains with distinct Vasa Helicase- and RNA-binding activities. Cell Rep 12: 587–598

Jeske M, Moritz B, Anders A & Wahle E (2011) Smaug assembles an ATP-dependent stable complex repressing nanos mRNA translation at multiple levels. EMBO J 30: 1–9

Jeske M, Müller CW & Ephrussi A (2017) The LOTUS domain is a conserved DEAD-box RNA helicase regulator essential for the recruitment of Vasa to the germ plasm and nuage. Genes Dev 31: 939–952

Johnstone O & Lasko P (2004) Interaction with elF5B is essential for Vasa function during development. Development 131: 4167–4178

Khemici V & Linder P (2016) RNA helicases in bacteria. Curr Opin Microbiol 30: 58–66

Kubíková J, Reinig R, Salgania HK & Jeske M (2020) LOTUS-domain proteins - Developmental effectors from a molecular perspective. Biol Chem 402: 7–23

Kubíková J, Ubartaitė G, Metz J & Jeske M (2023) Structural basis for binding of *Drosophila* Smaug to the GPCR Smoothened and to the germline inducer Oskar. Proceedings of the National Academy of Sciences 120: e2304385120

Kuramochi-Miyagawa S, Watanabe T, Gotoh K, Takamatsu K, Chuma S, Kojima-Kita K, Shiromoto Y, Asada N, Toyoda A, Fujiyama A, et al (2010) MVH in piRNA processing and gene silencing of retrotransposons. Genes Dev 24: 887–892

Lasko P (2013) The DEAD-box helicase Vasa: Evidence for a multiplicity of functions in RNA processes and developmental biology. Biochim Biophys Acta Gene Regul Mech 1829: 810–816

Lasko PF & Ashburner M (1988) The product of the Drosophila gene vasa is very similar to eukaryotic initiation factor-4A. Nature 335: 611–617 doi:10.1038/335611a0 [PREPRINT]

Leitão AL, Costa MC & Enguita FJ (2015) Unzippers, resolvers and sensors: a structural and functional biochemistry tale of RNA helicases. Int J Mol Sci 16: 2269–93

Liang L, Diehl-Jones W & Lasko P (1994) Localization of vasa protein to the Drosophila pole plasm is independent of its RNA-binding and helicase activities. Development 120: 1201–1211

Linder P & Jankowsky E (2011) From unwinding to clamping - the DEAD box RNA helicase family. Nat Rev Mol Cell Biol 12: 505–16

Linder P, Lasko P, Aschburner M, Leroy P, Nielsen P, Nishi K, Schnier J & Slonimski P (1989) Birth of the DEAD box. Nature 337: 121–122

Liu F, Putnam A & Jankowsky E (2008) ATP hydrolysis is required for DEAD-box protein recycling but not for duplex unwinding. Proc Natl Acad Sci U S A 105: 20209–20214

Liu N, Han H & Lasko P (2009) Vasa promotes Drosophila germline stem cell differentiation by activating mei-P26 translation by directly interacting with a (U)-rich motif in its 3’ UTR. Genes Dev 23: 2742–2752

Los G V., Encell LP, McDougall MG, Hartzell DD, Karassina N, Zimprich C, Wood MG, Learish R, Ohana RF, Urh M, et al (2008) HaloTag: A novel protein labeling technology for cell imaging and protein analysis. ACS Chem Biol 3: 373–382

Mallam AL, Jarmoskaite I, Tijerina P, Del Campo M, Seifert S, Guo L, Russell R & Lambowitz AM (2011) Solution structures of DEAD-box RNA chaperones reveal conformational changes and nucleic acid tethering by a basic tail. Proc Natl Acad Sci U S A 108: 12254–12259

Malone CD, Brennecke J, Dus M, Stark A, McCombie WR, Sachidanandam R & Hannon GJ (2009) Specialized piRNA Pathways Act in Germline and Somatic Tissues of the Drosophila Ovary. Cell 137: 522–535

Marintchev A, Edmonds KA, Marintcheva B, Hendrickson E, Oberer M, Suzuki C, Herdy B, Sonenberg N & Wagner G (2009) Topology and Regulation of the Human eIF4A/4G/4H Helicase Complex in Translation Initiation. Cell 136: 447–460

Mathys H, Basquin JÔ, Ozgur S, Czarnocki-Cieciura M, Bonneau F, Aartse A, Dziembowski A, Nowotny M, Conti E & Filipowicz W (2014) Structural and Biochemical Insights to the Role of the CCR4-NOT Complex and DDX6 ATPase in MicroRNA Repression. Mol Cell 54: 751–765

Von Moeller H, Basquin C & Conti E (2009) The mRNA export protein DBP5 binds RNA and the cytoplasmic nucleoporin NUP214 in a mutually exclusive manner. Nature Structural & Molecular Biology 2009 16:3 16: 247–254

Mohr G, Del Campo M, Mohr S, Yang Q, Jia H, Jankowsky E & Lambowitz AM (2008) Function of the C-terminal Domain of the DEAD-box Protein Mss116p Analyzed in Vivo and in Vitro. J Mol Biol 375: 1344–1364

Montpetit B, Thomsen ND, Helmke KJ, Seeliger MA, Berger JM & Weis K (2011) A conserved mechanism of DEAD-box ATPase activation by nucleoporins and InsP6 in mRNA export. Nature 472: 238–244

Nishida KM, Iwasaki YW, Murota Y, Nagao A, Mannen T, Kato Y, Siomi H & Siomi MC (2015) Respective functions of two distinct siwi complexes assembled during PIWI-interacting RNA biogenesis in bombyx germ cells. Cell Rep 10: 193–203

Nott TJ, Petsalaki E, Farber P, Jervis D, Fussner E, Plochowietz A, Craggs TD, Bazett-Jones DP, Pawson T, Forman-Kay JD, et al (2015) Phase Transition of a Disordered Nuage Protein Generates Environmentally Responsive Membraneless Organelles. Mol Cell 57: 936–947

Oberer M, Marintchev A & Wagner G (2005) Structural basis for the enhancement of eIF4A helicase activity by eIF4G. Genes Dev 19: 2212–2223

Ozata DM, Gainetdinov I, Zoch A, O’Carroll D & Zamore PD (2019) PIWI-interacting RNAs: small RNAs with big functions. Nat Rev Genet 20: 89–108

Ozgur S, Buchwald G, Falk S, Chakrabarti S, Prabu JR & Conti E (2015) The conformational plasticity of eukaryotic RNA-dependent ATPases. FEBS Journal 282: 850–863

Patil VS, Anand A, Chakrabarti A & Kai T (2014) The Tudor domain protein Tapas, a homolog of the vertebrate Tdrd7, functions in the piRNA pathway to regulate retrotransposons in germline of Drosophila melanogaster. BMC Biol 12: 1–15

Patil VS & Kai T (2010) Repression of Retroelements in Drosophila Germline via piRNA Pathway by the Tudor Domain Protein Tejas. Current Biology 20: 724–730

Pettersen EF, Goddard TD, Huang CC, Meng EC, Couch GS, Croll TI, Morris JH & Ferrin TE (2021) UCSF ChimeraX: Structure visualization for researchers, educators, and developers. Protein Sci 30: 70–82

Pontius BW (1993) Close encounters: why unstructured, polymeric domains can increase rates of specific macromolecular association. Trends Biochem Sci 18: 181–186

Price IF, Hertz HL, Pastore B, Wagner J & Tang W (2021) Proximity labeling identifies lotus domain proteins that promote the formation of perinuclear germ granules in c. Elegans. Elife 10

Prudêncio P & Guilgur L (2015) Protein Extraction from Drosophila Embryos and Ovaries. Bio Protoc 5

Pühringer T, Hohmann U, Fin L, Pacheco-Fiallos B, Schellhaas U, Brennecke J & Plaschka C (2020) Structure of the human core transcription-export complex reveals a hub for multivalent interactions. Elife 9: 1–65

Rajkowitsch L, Chen D, Stampfl S, Semrad K, Waldsich C, Mayer O, Jantsch MF, Konrat R, Bläsi U & Schroeder R (2007) RNA chaperones, RNA annealers and RNA helicases. RNA Biol 4: 118–130

Raz E (2000) The function and regulation of vasa-like genes in germ-cell development. Genome Biol 1: 1–6 doi:10.1186/gb-2000-1-3-reviews1017 [PREPRINT]

Rogers GW, Lima WF & Merrick WC (2001) Further characterization of the helicase activity of eIF4A. Substrate specificity. Journal of Biological Chemistry 276: 12598–12608

Sahayasheela VJ & Sugiyama H (2024) RNA G-quadruplex in functional regulation of noncoding RNA: Challenges and emerging opportunities. Cell Chem Biol 31: 53–70

Salgania HK, Metz J & Jeske M (2024) ReLo is a simple and rapid colocalization assay to identify and characterize direct protein–protein interactions. Nature Communications 2024 15:1 15: 1–12

Salgania HK, Metz J, Lingren E, Bleischwitz C, Hauser D, Máté KO, Bollack D, Lahr F, Bruckmann A, Garbelyanski A, et al (2025) Molecular insights into the Drosophila piRNA pathway via systematic ReLo protein interaction screening and structure prediction. Nucleic Acids Res 53: 13–14

Schindelin J, Arganda-Carreras I, Frise E, Kaynig V, Longair M, Pietzsch T, Preibisch S, Rueden C, Saalfeld S, Schmid B, et al (2012) Fiji: An open-source platform for biological-image analysis. Nat Methods 9: 676–682

Schuller SK, Schuller JM, Prabu JR, Baumgärtner M, Bonneau F, Basquin J & Conti E (2020) Structural insights into the nucleic acid remodeling mechanisms of the yeast tho-SUB2 complex. Elife 9: 1–51

Schüpbach T & Wieschaus E (1986) Maternal-effect mutations altering the anterior-posterior pattern of the Drosophila embryo. Rouxs Arch Dev Biol 195: 302–317

Schütz P, Bumann M, Oberholzer AE, Bieniossek C, Trachsel H, Altmann M & Baumann U (2008) Crystal structure of the yeast eIF4A-eIF4G complex: an RNA-helicase controlled by protein-protein interactions. Proc Natl Acad Sci U S A 105: 9564–9

Sengoku T, Nureki O, Nakamura A, Kobayashi S & Yokoyama S (2006) Structural Basis for RNA Unwinding by the DEAD-Box Protein Drosophila Vasa. Cell 125: 287–300

Sloan KE & Bohnsack MT (2018) Unravelling the Mechanisms of RNA Helicase Regulation. Trends Biochem Sci 43: 237–250

Sun Y, Atas E, Lindqvist LM, Sonenberg N, Pelletier J & Meller A (2014) Single-molecule kinetics of the eukaryotic initiation factor 4AI upon RNA unwinding. Structure 22: 941–948

Telley IA, Gáspár I, Ephrussi A & Surrey T (2012) Aster migration determines the length scale of nuclear separation in the Drosophila syncytial embryo. Journal of Cell Biology 197: 887–895

Vourekas A, Zheng K, Fu Q, Maragkakis M, Alexiou P, Ma J, Pillai RS, Mourelatos Z & Jeremy Wang P (2015) The RNA helicase MOV10L1 binds piRNA precursors to initiate piRNA processing. Genes Dev 29: 617–629

Vuzman D & Levy Y (2011) Intrinsically disordered regions as affinity tuners in protein–DNA interactions. Mol Biosyst 8: 47–57

Walker JE, Saraste M, Runswick MJ & Gay NJ (1982) Distantly related sequences in the alpha- and beta-subunits of ATP synthase, myosin, kinases and other ATP-requiring enzymes and a common nucleotide binding fold. EMBO J 1: 945–951

Wang C & Lehmann R (1991) Nanos is the localized posterior determinant in Drosophila. Cell 66: 637–647

Weis K & Hondele M (2022) The Role of DEAD-Box ATPases in Gene Expression and the Regulation of RNA–Protein Condensates. https://doi.org/101146/annurev-biochem-032620-105429 91: 197–219

Wenda JM, Homolka D, Yang Z, Spinelli P, Sachidanandam R, Pandey RR & Pillai RS (2017) Distinct Roles of RNA Helicases MVH and TDRD9 in PIWI Slicing-Triggered Mammalian piRNA Biogenesis and Function. Dev Cell 41: 623–637.e9

Wong E V., Cao W, Vörös J, Merchant M, Modis Y, Hackney DD, Montpetit B & De La Cruz EM (2016) Pi Release Limits the Intrinsic and RNA-Stimulated ATPase Cycles of DEAD-Box Protein 5 (Dbp5). J Mol Biol 428: 492–508

Xiol J, Spinelli P, Laussmann MA, Homolka D, Yang Z, Cora E, Couté Y, Conn S, Kadlec J, Sachidanandam R, et al (2014) RNA clamping by Vasa assembles a piRNA amplifier complex on transposon transcripts. Cell 157: 1698–1711

Yamazaki H, Namba Y, Kuriyama S, Nishida KM, Kajiya A & Siomi MC (2023) Bombyx Vasa sequesters transposon mRNAs in nuage via phase separation requiring RNA binding and self-association. Nat Commun 14: 1942

Yang Q, Del Campo M, Lambowitz AM & Jankowsky E (2007) DEAD-box proteins unwind duplexes by local strand separation. Mol Cell 28: 253–263

Yao Q, Cao G, Li M, Wu B, Zhang X, Zhang T, Guo J, Yin H, Shi L, Chen J, et al (2018) Ribonuclease activity of MARF1 controls oocyte RNA homeostasis and genome integrity in mice. Proc Natl Acad Sci U S A 115: 11250–11255

Zaessinger S, Busseau I & Simonelig M (2006) Oskar allows nanos mRNA translation in Drosophila embryos by preventing its deadenylation by Smaug/CCR4. Development 133: 4573–83

Zhang F, Wang J, Xu J, Zhang Z, Koppetsch BS, Schultz N, Vreven T, Meignin C, Davis I, Zamore PD, et al (2012) UAP56 couples piRNA clusters to the perinuclear transposon silencing machinery. Cell 151: 871–884

Zhang X, Yu L, Ye S, Xie J, Huang X, Zheng K & Sun B (2019) MOV10L1 Binds RNA G-Quadruplex in a Structure-Specific Manner and Resolves It More Efficiently Than MOV10. iScience 17: 36–48

