## Supplementary Data for "Localization-dependent activation of the DEAD-box ATPase Vasa by eLOTUS domains"

### Inventory:

Supplementary Figures 1-9
Supplementary Tables 1-4
Supplementary Methods
Supplementary References

### SUPPLEMENTARY FIGURES AND LEGENDS

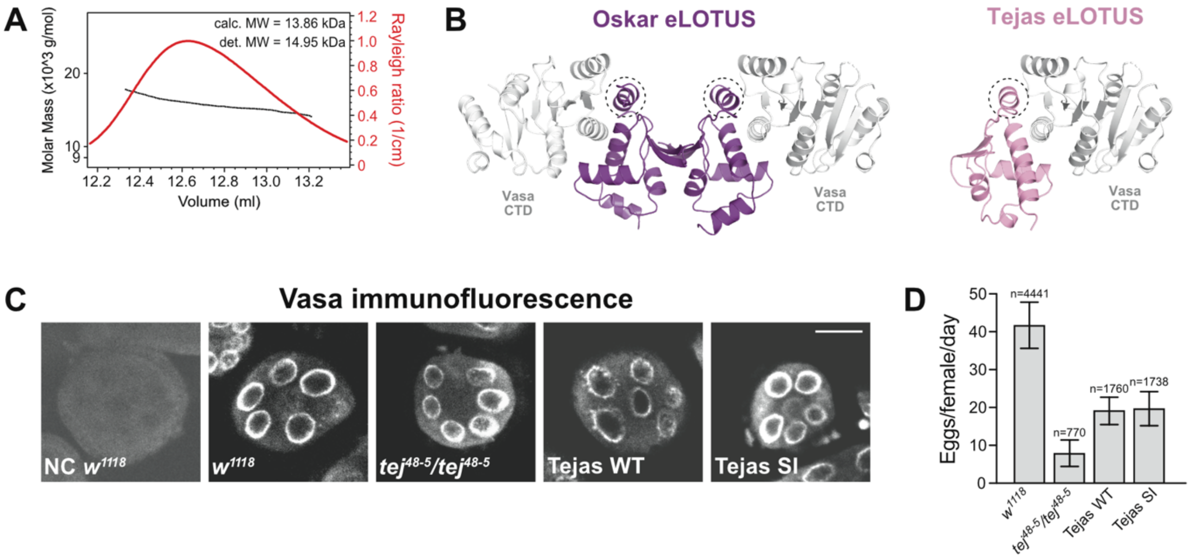

**Supplementary Figure 1.**
(A) MALS analysis of His-Tejas-eLOTUS (aa 1-100).
(B) Crystal structure of the Vasa-CTD-Oskar-eLOTUS complex (PDB ID: 5NT7) (left) and SWISS-MODEL-generated homology model of the Tejas-eLOTUS-Vasa-CTD complex (right). The C-terminal $\alpha$-helix specific to eLOTUS domains is highlighted with a dashed circle.
(C) Vasa immunostaining using confocal fluorescence microscopy of stage 3 egg chambers expressing the Tejas WT or Tejas SI transgene in the homozygous *tej*$^{48-5}$ null background. Scale bar is 20 µm. In the negative control (NC), the anti-Vasa antibody was omitted.
(D) Egg laying analyses of females with the genotype as indicated. Tejas WT or Tejas SI transgenes were expressed in the homozygous *tejas*$^{48-5}$ null background.

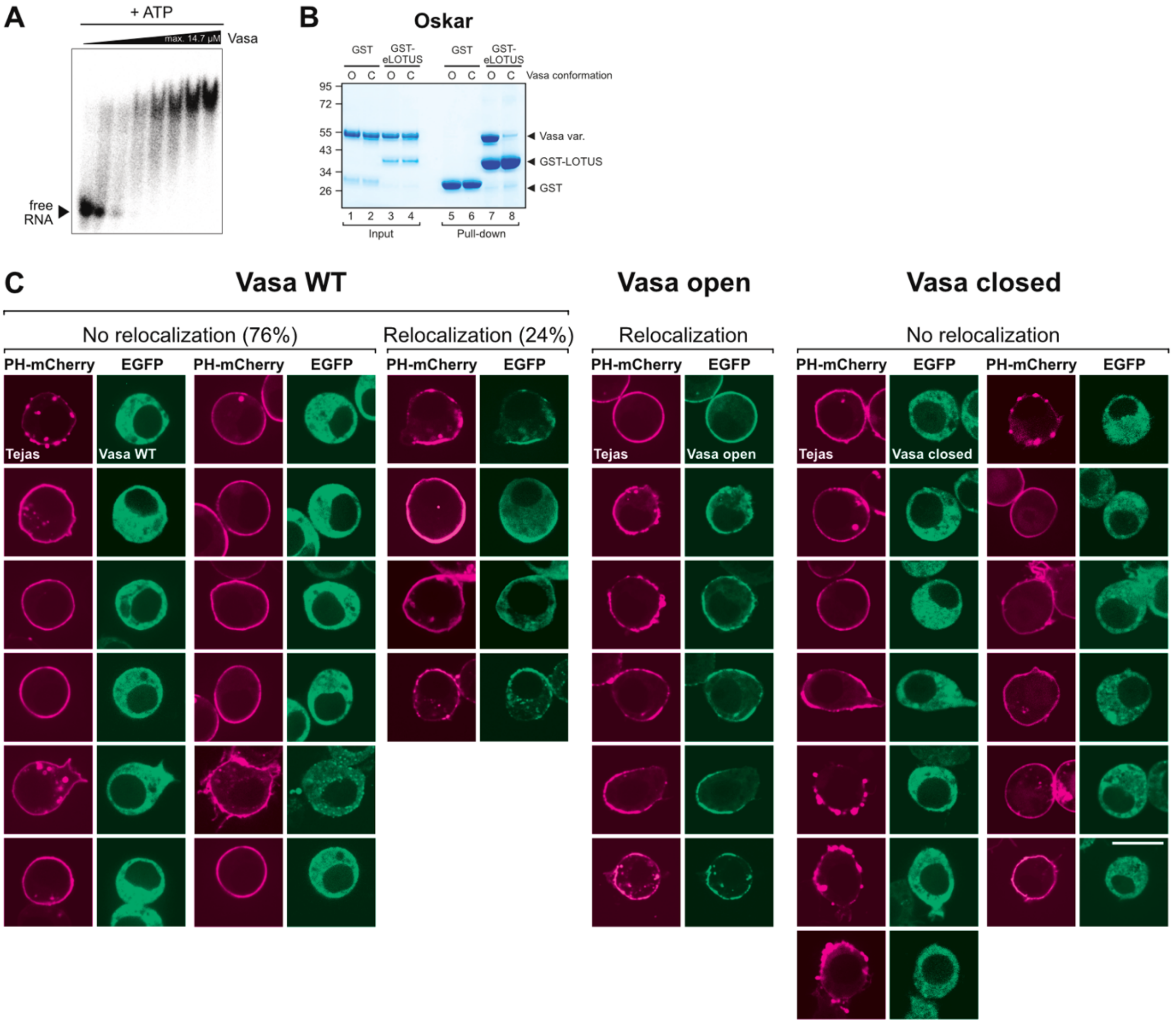

**Supplementary Figure 2.**
(A) Electrophoretic mobility shift assays (EMSAs) using 14.7 µM His-Vasa 200-623 and seven 2-fold serial dilutions incubated with 50 nM R26 ssRNA in the presence of 1 mM ATP and 5 mM $MgCl_2$.
(B) GST pull-down assay using 10 µM GST or GST-Oskar-eLOTUS (aa 139-240) and 20 µM His-Vasa 200-623. For the open conformation ('O'), the K295N mutant variant was used. For the closed conformation ('C'), the WT core was incubated with 40 µM of R13 ssRNA oligo and 2 mM ATP prior to incubation with the eLOTUS domain. Molecular weight marker (in kDa) is indicated at the left.
(B) ReLo assay showing several cells with the localization of Vasa WT, Vasa open (K295N), and Vasa closed (E400Q) in the presence of Tejas (related to **Figure 2D**). PH-mCherry and EGFP constructs were transiently coexpressed in S2R+ cells. After 26 hours, subcellular protein localization was examined by confocal live fluorescence microscopy. Interaction of Vasa WT with Tejas is only observed in a subset of cells (24%). In all of the cells, Vasa WT localizes exclusively in the cytoplasm in the presence of Tejas. Scale bar is 10 µm.

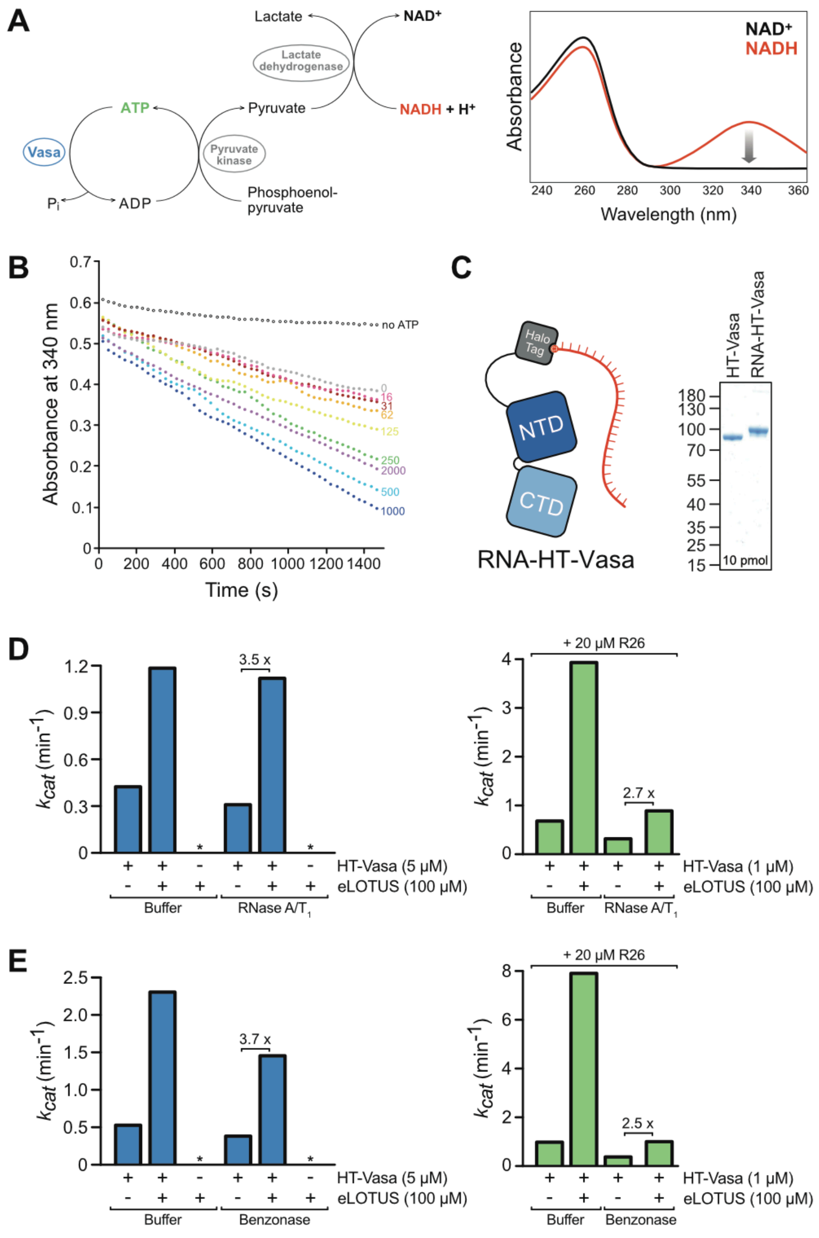

**Supplementary Figure 3.**
(A) Scheme of the NADH-coupled ATPase assay. ADP and inorganic phosphate produced during hydrolysis drive a two-step reaction involving pyruvate kinase and lactate dehydrogenase, resulting in NADH oxidation detectable at 340 nm.
(B) Absorbance at 340 nm measured over time during an NADH-coupled ATPase assay showing inhibition of ATPase activity in the presence of 2000 µM tRNA (violet dots). The reaction contained 5 µM His-Vasa core, 5 mM ATP, and the indicated concentrations of gel filtration-purified tRNA.
(C) Illustration of HaloTag-Vasa 200-661 (HT-Vasa) covalently linked to R26 RNA (RNA-HT-Vasa) (left panel). Coomassie-stained gel showing the completeness of the Halo-Vasa - RNA coupling. The molecular weight marker (kDa) is indicated on the left.
(D, E) RNA-independent ATPase activity of HT-Vasa measured by an NADH-coupled ATPase assay using 5 mM ATP in the presence or absence of His-Tejas-eLOTUS. Prior to the assay, Vasa and Tejas were incubated for 1 h with RNase A/T1 (D) or Benzonase (E), or with the corresponding buffers as controls (left panels). To verify RNase activity, an equivalent experiment was performed in parallel in the presence of 20 µM R26 RNA (right panels).

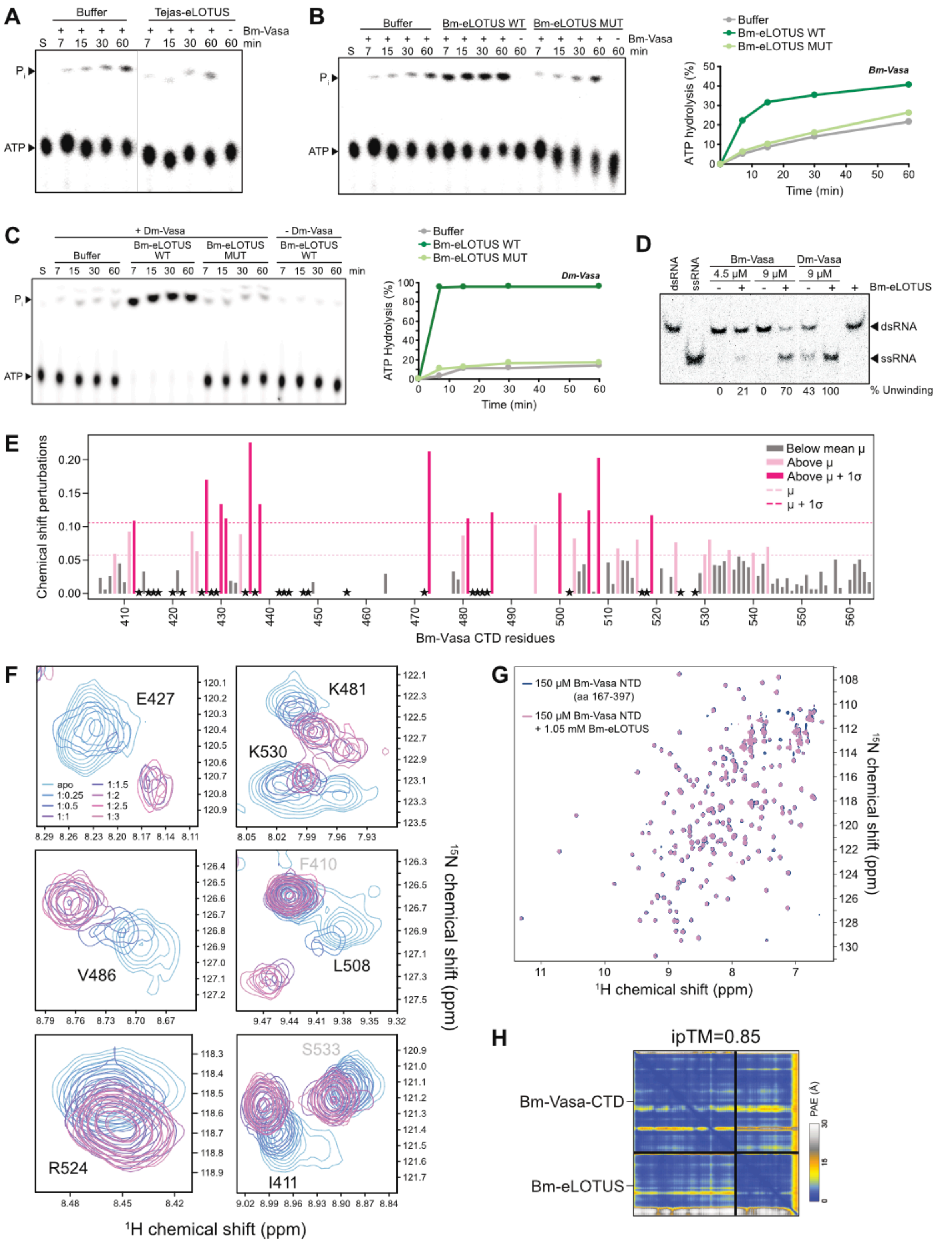

**Supplementary Figure 4.**

(A) ATPase assay using 2.5 μM of the Bm-Vasa core (aa 135-564), 10 μM R26 ssRNA, 8 nM [γ-$^{32}$P] ATP, in the presence of buffer or 100 μM Tejas-eLOTUS. All samples originated from one experiment but were loaded onto two thin layer plates. Lanes with irrelevant data were removed.

(B) ATPase assay using 2.5 μM of Bm-Vasa core, 10 μM R26 ssRNA, 8 nM [γ-$^{32}$P] ATP in the presence of buffer or 100 μM BmTDRD7-eLOTUS (1-100) WT or S13E/I81E mutant.

(C) ATPase assay using 2.5 µM of the His-Vasa 200-661*, 10 µM R26 ssRNA oligo, 8 nM [$\gamma$-$^{32}$P] ATP, in the presence of buffer, 100 µM wildtype or mutant Bm-TDRD7-eLOTUS (aa 1-100) domains.
(D) dsRNA unwinding assay performed in the presence of 3 mM ATP and an ATP-regenerating system, 5 nM labeled dsR13 oligonucleotide, and 500 nM unlabeled competitor R13 ssRNA. Bm-Vasa and Dm-Vasa cores were incubated for 1 h at concentrations indicated in the presence or absence of 100 µM Bm-TDRD7-eLOTUS. Lanes with irrelevant data were removed.
(E) Chemical shift perturbations (CSP) in Bm-Vasa CTD upon Bm-eLOTUS binding at 1:3 molar ratio. Per-residue CSP is calculated as $\sqrt{(\Delta\delta_H)^2 + (0.14 * \Delta\delta_N)^2}$, where $\Delta\delta_H$ and $\Delta\delta_N$ are the $^1$H and $^{15}$N chemical shift differences, respectively, in the presence and absence of the eLOTUS domain. The dashed lines in light pink and dark pink indicate the mean (µ) threshold and significantly perturbed residues (one standard deviation $\sigma$ from the mean µ). An asterisk (*) shows residues with overlapped chemical shifts or that disappeared upon adding the ligand. The unassigned residues in apo are left blank in the plot. Related to **Figures 3D and 3E**.
(F) Expanded views of selected amide resonances from **Figure 3D** during eLOTUS titration.
(G) Overlay of $^1$H-$^{15}$N HSQC spectra of the *Bombyx* Vasa-NTD in the absence (dark blue) and presence (pink) of the *Bombyx* TDRD7 eLOTUS domain, showing no significant CSPs.
(H) PAE plot of the structure prediction shown in **Figure 3E**.

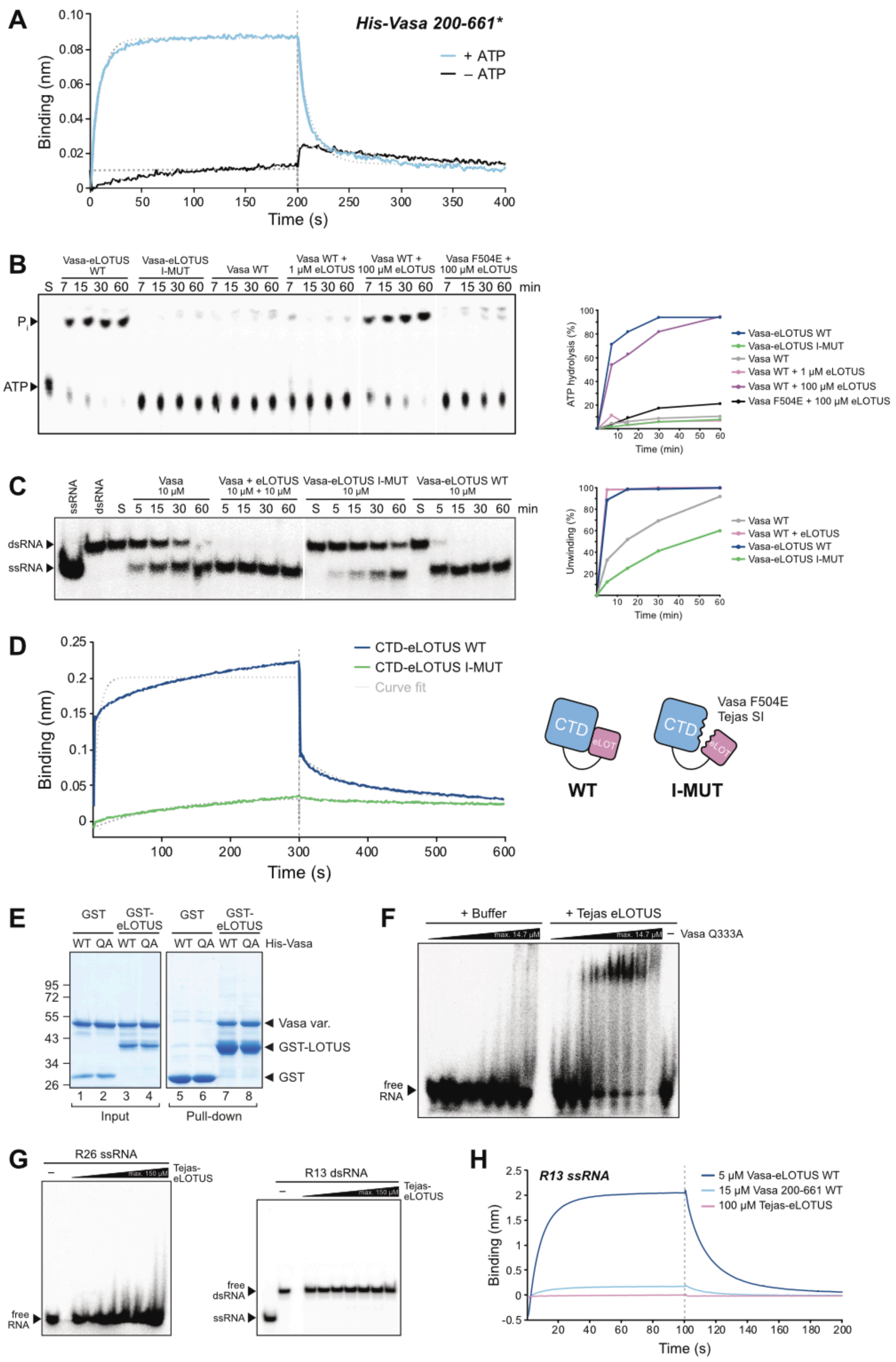

**Supplementary Figure 5.**
(A) Biolayer interferometry assay showing the binding of 50 μM His-Vasa 200-661* to sensor-coupled RNA in the presence of 5 mM ATP but not in its absence. Streptavidin biosensors were loaded in the presence of 10 nM biotinylated R13 ssRNA oligo. His-

Vasa 200-661* contains a short GS linker at its C-terminus not affecting Vasa function. Kinetic parameters in the presence of ATP: $k_{on}$ = 790 $M^{-1}s^{-1}$, $k_{off}$ = 4.3 $min^{-1}$, $K_D$ = 90.6 µM.

(B) ATPase assay using 10 µM R26 ssRNA oligo, 8 nM [$\gamma$-$^{32}$P] ATP, 1 µM of His-Vasa 200-661* WT, His-Vasa-200-661* MUT (F504E) or of the Vasa-eLOTUS WT or MUT fusion proteins in the absence or presence of the Tejas-eLOTUS domain. All samples originated from one experiment but were loaded onto two thin layer plates. His-Vasa 200-661* contains a short GS linker at its C-terminus not affecting Vasa function.

(C) dsRNA unwinding assay in the presence of 3 mM ATP, 25 nM labeled R13 dsRNA oligo, and 500 nM unlabeled competitor R13 ssRNA oligo using His-Vasa 200-623 ("Vasa") in the absence or presence of the Tejas-eLOTUS domain or using the Vasa-eLOTUS WT or MUT fusion proteins. All samples originated from one experiment but were loaded onto two gels.

(D) Biolayer interferometry sensograms comparing the RNA-binding affinities of 100 µM Vasa-CTD-Tejas-eLOTUS WT and I-MUT fusion proteins. Streptavidin biosensors were loaded in the presence of 10 nM biotinylated R13 ssRNA.

(E) GST pull-down assay using 1 nmol GST or GST-Tejas-eLOTUS (aa 1-100) and 2 nmol His-Vasa 200-623 WT or Q333A (QA). The data show that the QA mutation does not affect eLOTUS binding. Molecular weight marker (in kDa) is indicated at the left. Samples originated from one experiment but were loaded onto two gels.

(F) EMSAs using 50 nM R26 ssRNA and 14.7 µM and seven 2-fold serial dilutions of His-Vasa 200-623 Q333A in the absence ("Buffer") or presence of 150 µM His-Tejas-eLOTUS. Vasa QA alone does not efficiently bind to RNA in EMSAs (compare with Vasa WT in **Supplementary Figure 2A**). RNA binding becomes evident in the presence of eLOTUS.

(G) EMSAs using 50 nM R26 ssRNA (left panel) and 150 µM and five 2-fold serial dilutions of His-Tejas-eLOTUS or 50 nM R13 dsRNA (right panel) and 150 µM and seven 2-fold serial dilutions of His-Tejas-eLOTUS.

(H) Biolayer interferometry assay showing that the eLOTUS domain alone did not bind to the biosensor-coupled RNA oligo. RNA binding by His-Vasa 200-661* and the Vasa-eLOTUS WT fusion are shown for comparison. Streptavidin biosensors were loaded in the presence of 100 nM biotinylated R13 ssRNA oligo. His-Vasa 200-661* contains a short GS linker at its C-terminus not affecting Vasa function.

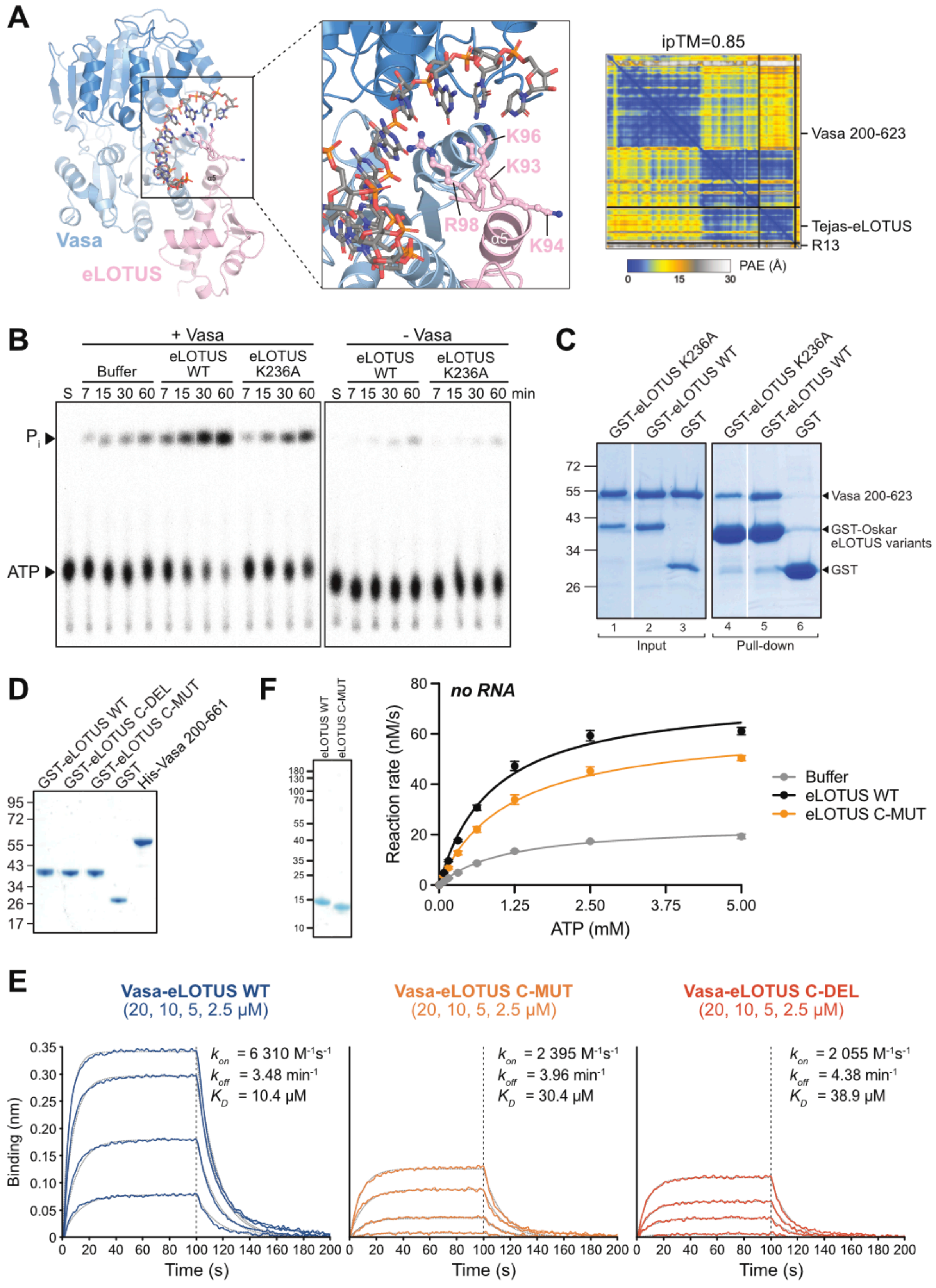

**Supplementary Figure 6.**
(A) AlphaFold3-generated structural model of the complex composed of the Vasa core, the Tejas-eLOTUS domain and an R13 ssRNA oligo.

(B) ATPase assay using 5 µM of the His-Vasa 200-661, 10 µM R26 ssRNA oligo, 8 nM [$\gamma$-$^{32}$P] ATP, in the presence of buffer, 25 µM wildtype or mutant Oskar-eLOTUS domain as indicated.
(C) GST pull-down assay using 1 nmol GST, GST-Oskar-eLOTUS WT or K236A mutant, and 2 nmol His-Vasa 200-623. Molecular weight marker (in kDa) is indicated at the left. Input and pull-down samples originated from one experiment but were loaded onto two gels. Lanes with irrelevant data were removed.
(D) Coomassie-stained gel showing the relative amounts (50 pmol) of the proteins used in the experiments shown in **Figures 4E and 4F**. Molecular weight marker (in kDa) is indicated at the left. Lanes with irrelevant data were removed.
(E) Full biolayer interferometry sensograms of the experiment shown in **Figure 4G**.
(F) NADH-coupled ATPase assay in the absence of RNA using 5 µM HT-Vasa 200-661 and increasing concentrations of ATP in the absence (buffer) or presence of 100 µM His-Tejas-eLOTUS WT or C-MUT.

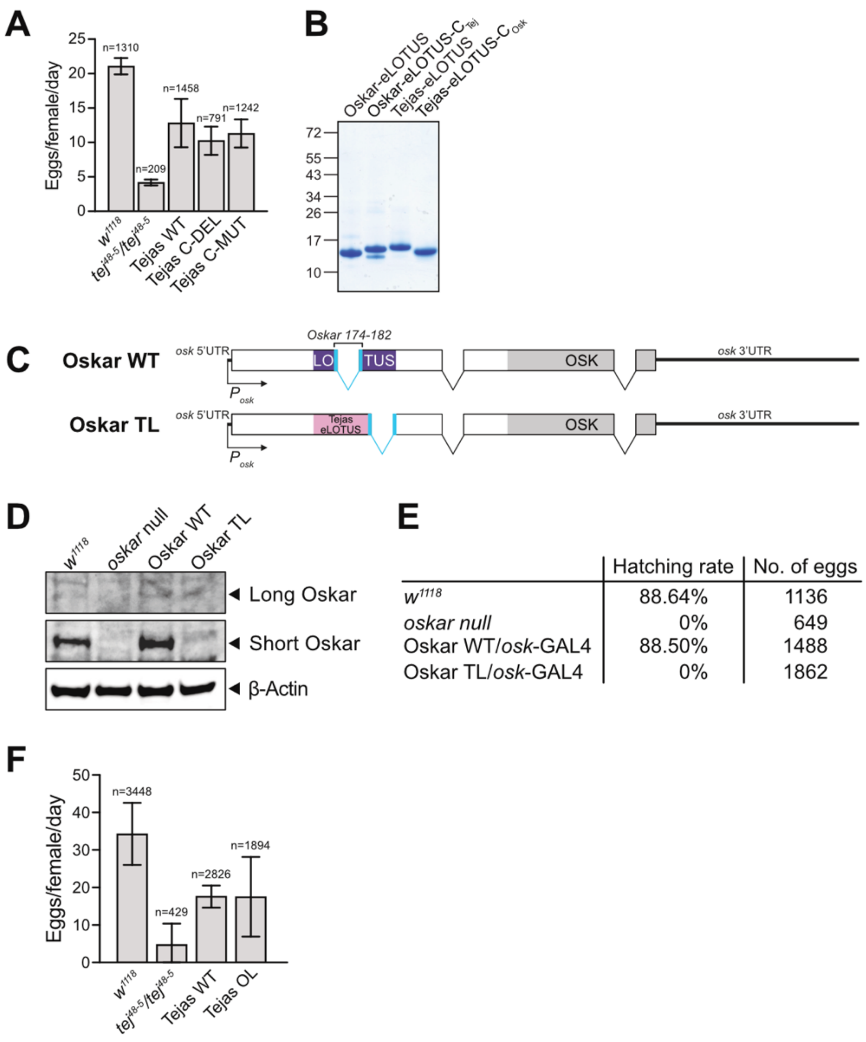

| | Hatching rate | No. of eggs |
|---|---|---|
| $w^{1118}$ | 88.64% | 1136 |
| *oskar null* | 0% | 649 |
| Oskar WT/*osk*-GAL4 | 88.50% | 1488 |
| Oskar TL/*osk*-GAL4 | 0% | 1862 |

**Supplementary Figure 7.**
(A) Egg laying by females with the genotype as indicated. Figure relates to experiment in **Figure 5D**.
(B) Coomassie-stained gel showing the relative amounts (250 pmol) of the proteins used in the experiments shown in **Figure 6C**. Molecular weight marker (in kDa) is indicated at the left.
(C) Scheme of the genomic Oskar-WT and Oskar-OL transgenes. The *oskar* mRNA is translated from two different in-frame start codons giving rise to the Long Oskar (aa 1-606) and Short Oskar (aa 139-606) protein isoforms (Markussen *et al*, 1995). For proper *oskar* mRNA localization and Oskar protein production *in vivo*, the *oskar* mRNA has to undergo a splicing event, which forms a specific stem-loop structure, the SOLE element, within the region that codes for the Oskar-eLOTUS domain (blue color) (Hachet & Ephrussi, 2004; Ghosh *et al*, 2012; Simon *et al*, 2015). To not interfere with *oskar* mRNA localization *in vivo*, we have introduced the Tejas eLOTUS domain upstream of the the SOLE element. For all studies, the Oskar-WT and Oskar-TL transgenes were expressed in the *oskar* null (*osk*-STOP-cTRICK) background.
(D) Western blot using anti-Oskar or anti-Tubulin antibodies showing that the Short Oskar isoform is specifically destabilized when it contains the Tejas-eLOTUS domain.
(E) Hatching rate analyses of females with the genotypes as indicated.
(F) Egg laying by females with the genotype as indicated. Figure relates to experiment in **Figure 7D**.

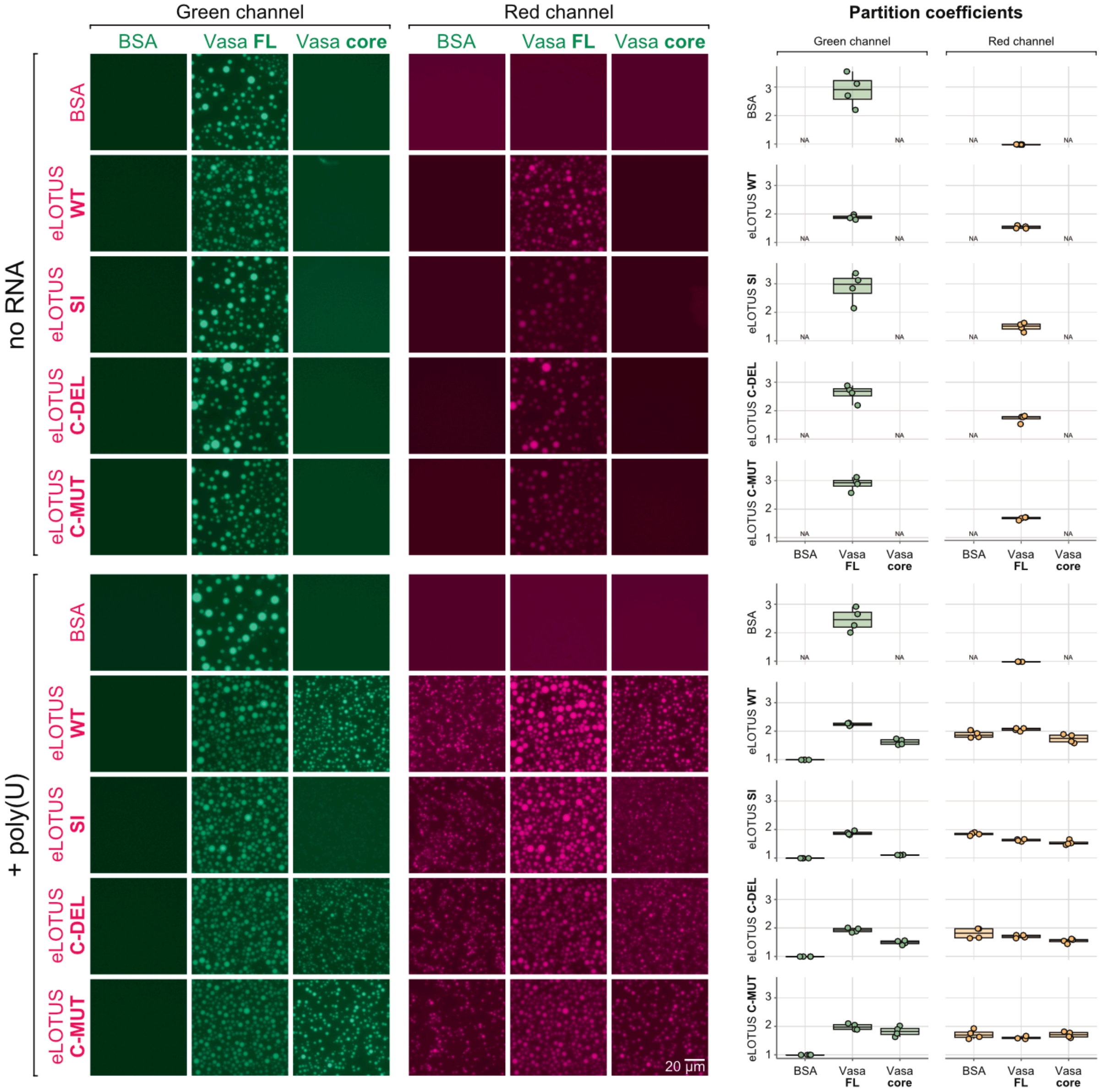

**Supplementary Figure 8.**
*In vitro* condensation assays were performed with 5 µM Vasa or BSA and 25 µM eLOTUS variants or BSA, with or without poly(U) RNA as indicated. For imaging, 565-ATTO-labeled eLOTUS variants (red) and 488-ATTO-labeled Vasa constructs (green) were spiked into reactions at 1% of the respective total protein concentration. The partition coefficient was calculated as the ratio of the mean fluorescence intensity inside droplets to that of the surrounding phase.

In the absence of RNA (upper panels), full length Vasa (Vasa FL) formed condensates independently of eLOTUS, and eLOTUS co-condensed with Vasa FL.

In the presence of poly(U) (lower panels), the eLOTUS domain alone formed condensates too. The Vasa core, lacking the N- and C-terminal intrinsically disordered tails, did not form condensates. However, eLOTUS WT, C-DEL, and C-MUT recruited Vasa core to condensates, whereas the interaction-deficient eLOTUS SI mutant did not.

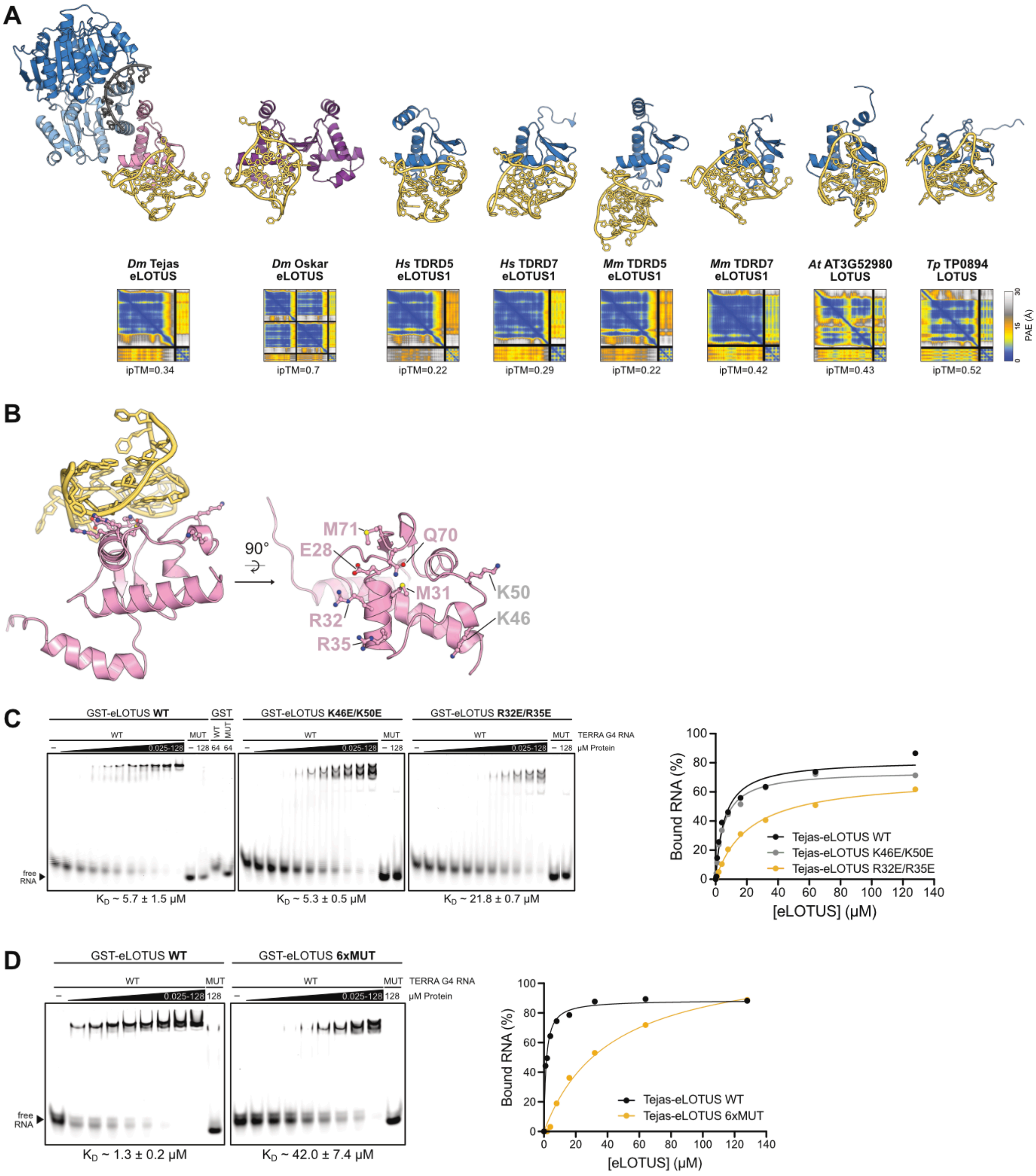

**Supplementary Figure 9.**

(A) AlphaFold3-generated structural models of different LOTUS domains interacting with TERRA G4 RNA. All domains were previously shown to bind to TERRA G4 (Ding *et al*, 2020). Domain boundaries and RNA sequences were derived from Ding et al. 2020. Only the G4 complexes with the Oskar and the bacterial LOTUS domains show high confidence in the prediction (ipTM>0.5).

(B) Residues mutated in subsequent experiments are highlighted in ball-and-stick representation. Residues that upon co-mutation show reduced G4 binding are shown in pink, and residues without effect in gray.

(C) EMSAs using 6-FAM-labeled TERRA G4 WT and MUT RNA. TERRA G4 MUT does not form a G-quadruplex structure (Ding *et al*, 2020). Binding was tested with 126 µM GST-Tejas-eLOTUS WT, K46E/K50E, and R32E/R35E, followed by ten 2-fold serial dilutions. GST alone (64 µM) was used as a control. Binding curves were fitted using a one-site binding model. $K_D$ values are indicated below the gels.

(D) EMSAs performed as in (E) using GST-Tejas-eLOTUS WT and the E28A/M31A/R32E/R35E/Q70A/M71A mutant (6xMUT), followed by eight 2-fold serial dilutions.

### SUPPLEMENTARY TABLES

#### Supplementary Table 1. DEAD-box protein ATPase activities

Protein concentrations, $K_M$ and $k_{cat}$ values are reported in the units as provided in the original publication. The protein domains used in the experiments are indicated in parentheses using the following abbreviations: DD, dimerization domain; core, N- and C-terminal RecA domains; C, C-terminal unstructured tail; CTE, C-terminal extension (structured); N, N-terminal unstructured tail; NTE, N-terminal extension (structured); RBD, RNA-binding domain; RRM, RNA recognition motif; SPRY, SPIa kinase and RYanodine receptor domain.

| | | *ATP titration in absence of RNA* | | | *ATP titration at saturating [RNA]* | | | *RNA titration at saturating [ATP]* | | | |
|---|---|---|---|---|---|---|---|---|---|---|---|
| **DEAD-box protein** | **[Protein]** | **$K_M$** (ATP) | **$k_{cat}$** (ATP) | **$k_{cat}/K_M$** (ATP) | **$K_M$** (ATP) | **$k_{cat}$** (ATP) | **$k_{cat}/K_M$** (ATP) | **$K_M$** (RNA) | **$k_{cat}$** (RNA) | **$k_{cat}/K_M$** (RNA) | **Reference** |
| **Vasa** 200-661 (core + C) *D. melanogaster* | 5 µM | 1310 µM | 0.43 $min^{-1}$ | 5.3 $M^{-1}s^{-1}$ | 961 µM | 0.29 $min^{-1}$ | 4.2 $M^{-1}s^{-1}$ | 429 µM | 2.18 $min^{-1}$ | 84 $M^{-1}s^{-1}$ | This study |
| **eIF4A** FL (core) *S. cerevisiae* | 0.25 µM | - | - | - | 470 µM | - | - | 18 µM | - | - | (Blum *et al*, 1992) |
| | 1 µM | - | - | - | - | - | - | 15 µM | 0.005 $s^{-1}$ | 333 $M^{-1}s^{-1}$ | (Andreou *et al*, 2017) |
| *M. musculus* | 4 µM | - | - | - | 80 µM | 3 $min^{-1}$ | 625 $M^{-1}s^{-1}$ | - | - | - | (Lorsch & Herschlag, 1998) |
| **eIF4AIII** FL (core) *H. sapiens* | 0.4 µM | - | - | - | 240 µM | 7 $s^{-1}$ | 29.1 x $10^3$ $M^{-1}s^{-1}$ | - | - | - | (Noble & Song, 2007) |
| **Dbp5** FL (NTE + core) *S. cerevisiae* | 0.6-1 µM | 1700 µM | 0.043 $s^{-1}$ | 25.3 $M^{-1}s^{-1}$ | - | - | - | 3.4 mM | 0.92 $s^{-1}$ | 270 $M^{-1}s^{-1}$ | (Wong *et al*, 2016) |
| **DbpA** FL (core + RRM) *E. coli* | 2-20 nM | - | not detectable | - | 65 µM | 5.2 $s^{-1}$ | 80 x $10^3$ $M^{-1}s^{-1}$ | 1 nM | 5.3 $s^{-1}$ | 5.3 x $10^9$ $M^{-1}s^{-1}$ | (Henn *et al*, 2008) |
| | 30 nM | - | - | - | 330 µM | 1.27 $s^{-1}$ | 3.9 x $10^3$ $M^{-1}s^{-1}$ | - | - | - | (Moore *et al*, 2017) |
| | 30 nM | - | - | - | 600 µM | 1 $s^{-1}$ | 1.7 x $10^3$ $M^{-1}s^{-1}$ | - | - | - | (López De Victoria *et al*, 2017) |
| | 5-10 nM | | | | 235 µM | 610 $min^{-1}$ | 43 x $10^3$ $M^{-1}s^{-1}$ | 24 nM | 580 $min^{-1}$ | 404 x $10^6$ $M^{-1}s^{-1}$ | (Tsu & Uhlenbeck, 1998) |

| | | *ATP titration in absence of RNA* | | | *ATP titration at saturating [RNA]* | | | *RNA titration at saturating [ATP]* | | | |
|---|---|---|---|---|---|---|---|---|---|---|---|
| **DEAD-box protein** | **[Protein]** | **$K_M$** (ATP) | **$k_{cat}$** (ATP) | **$k_{cat}/K_M$** (ATP) | **$K_M$** (ATP) | **$k_{cat}$** (ATP) | **$k_{cat}/K_M$** (ATP) | **$K_M$** (RNA) | **$k_{cat}$** (RNA) | **$k_{cat}/K_M$** (RNA) | **Reference** |
| **Mss116p** 73-597 (core + CTE) *S. cerevisiae* | 50 nm | 288 μM | 0.26 $s^{-1}$ | 903 $M^{-1}s^{-1}$ | 252 μM | 1.5 $s^{-1}$ | 62 x $10^3$ $M^{-1}s^{-1}$ | 205 nM | 1.8 $s^{-1}$ | 8.8 x $10^6$ $M^{-1}s^{-1}$ | (Cao *et al*, 2011) |
| **DDX1** FL (core + SPRY) *H. sapiens* | 1 μM | 1750 μM | 0.096 $s^{-1}$ | 54.9 $M^{-1}s^{-1}$ | 11.7 μM | 0.168 $s^{-1}$ | 14.4 x $10^3$ $M^{-1}s^{-1}$ | 70 nM | 0.141 $s^{-1}$ | 2.0 x $10^6$ $M^{-1}s^{-1}$ | (Kellner *et al*, 2015) |
| **Has1p** FL (core) *S. cerevisiae* | 390 nM | - | - | - | 440 μM | 5.4 $min^{-1}$ | 204.5 $M^{-1}s^{-1}$ | 160 nM | - | - | (Rocak *et al*, 2005) |
| **An3** FL (N + core) *X. laevis* | 190 nM | - | - | - | 65 μM | 9 $min^{-1}$ | 2.3 x $10^3$ $M^{-1}s^{-1}$ | - | - | - | (Askjaer *et al*, 2000) |
| **CrhR** FL (core) *Synechocystis sp.* | 8.75 nM | - | - | - | 580 μM | 500 $min^{-1}$ | 14.3 x $10^3$ $M^{-1}s^{-1}$ | - | - | - | (Chamot *et al*, 2005) |
| **CsdA** FL (core + DD + RBD) *E. coli* | 0.05-0.5 μM | - | - | - | - | 90 $min^{-1}$ | - | - | - | - | (Bizebard *et al*, 2004) |
| **SrmB** FL (core + CTE) *E. coli* | 20 nM | - | - | - | 180 μM | 20-40 $min^{-1}$ | 1.7-3.3 x $10^3$ $M^{-1}s^{-1}$ | 220 nM | 24 $min^{-1}$ | 1.8 x $10^6$ $M^{-1}s^{-1}$ | (Kossen *et al*, 2002) |
| **Dbp8** FL (core + C) *S. cerevisiae* | 1 μM | 194 μM | 17 $min^{-1}$ | 1460 $M^{-1}s^{-1}$ | - | - | - | - | - | - | (Granneman *et al*, 2006) |
| **Hera** FL (core + DD + RBD) *T. thermophilus* | 150 nM | - | - | - | - | - | - | 240 μM | 2.3 $s^{-1}$ | 9.6 x $10^3$ $M^{-1}s^{-1}$ | (Donsbach *et al*, 2025) |
| 1-419 (core + DD) | 300 nM | - | - | - | - | - | - | 871 μM | 0.9 $s^{-1}$ | 1033 $M^{-1}s^{-1}$ | |
| 1-365 (core) | 300 nM | - | - | - | - | - | - | not detectable | not detectable | not detectable | |
| **YxiN** FL (core + RRM) *B. subtilis* | 10 nM | - | 0.03 $min^{-1}$ | - | 230 μM | 163 $min^{-1}$ | 1.18 x $10^4$ $M^{-1}s^{-1}$ | 4.1 nM | 170 $min^{-1}$ | 6.9 x $10^8$ $M^{-1}s^{-1}$ | (Kossen & Uhlenbeck, 1999) |

### Supplementary Table 2. DEAD-box protein activators

| DEAD-box ATPase | Stimulator | Binding mode and effect on DEAD-box protein activity |
|---|---|---|
| Vasa | eLOTUS domain | Binds Vasa-CTD in open conformation and stimulates ATPase and dsRNA unwinding activity mainly by increasing RNA engagement. Stimulation is mediated by an unstructured positively charged stretch. Catalytic efficiency is enhanced ~135 to 465-fold (Tejas). |
| eIF4A | MIF4G domain of eIF4G | Interacts with both RecA domains of the core promoting 'half-open' conformation, thereby accelerating both core closing and opening (Schütz *et al*, 2008; Hilbert *et al*, 2011). Increases phosphate release (Hilbert *et al*, 2011) and ATP turnover but not dsRNA unwinding (Andreou & Klostermeier, 2014; Andreou *et al*, 2017). Catalytic efficiency is enhanced ~2.3 to 13-fold. |
| eIF4A | MIF4G domain of DAP5 | Interacts with both RecA domains and stimulates dsRNA unwinding (Virgili *et al*, 2013). |
| Dbp5 | MIF4G domain of Gle1 | Contacts both RecA domains and stimulates the RNA release (Montpetit *et al*, 2011). Stabilizes ATP binding and increases the phosphate release (Gray *et al*, 2022; Wong *et al*, 2016). Catalytic efficiency is enhanced ~300-fold (without RNA). |
| Fal1 | MIF4G domain of Sgd1 | Stimulates ATP hydrolysis (Davila Gallesio *et al*, 2020). |
| DDX6 | MIF4G domain of CNOT1 | Binds both Rec-A domains and stimulates ATPase activity by modulating conformation (Mathys *et al*, 2014; Chen *et al*, 2014). |
| eIF4A | eIF4B | eIF4B increases dsRNA unwinding and, in the presence of eIF4G, RNA binding. The 7-repeat region of eIF4B interacts with and stabilizes the RNA-bound closed conformation (Andreou & Klostermeier, 2014; Andreou *et al*, 2017; Rogers *et al*, 2001). In the presence of eIF4G, catalytic efficiency is enhanced by eIF4B ~14 to 52-fold. |
| eIF4AI | eIF4H | Stimulates dsRNA unwinding (Rogers *et al*, 2001; Sun *et al*, 2012). Binds the CTD in NMR experiments (Marintchev *et al*, 2009). Binds also the NTD in presence of RNA (Sun *et al*, 2014). |
| eIF4AIII | SELOR domain of MLN51 /Barentsz | RNA binding protein that interacts with both RecA domains (Andersen *et al*, 2006; Bono *et al*, 2006). Increases RNA binding and ATPase activity (Noble & Song, 2007). Catalytic efficiency is enhanced ~330-fold. |
| Dbp8 | Esf2 | Increases affinity for ATP (Granneman *et al*, 2006). Catalytic efficiency is enhanced ~7-fold (without RNA). |

**Supplementary Table 3. DNA constructs used in this study**
All constructs are *Drosophila melanogaster* sequences unless otherwise indicated. The specific protein isoforms (iso) used are indicated.

| **Vector** (insertion site) (code) | **Final DNA construct** | **DNA template information** | **Code** |
|---|---|---|---|
| **pGEX-6P-1** (Cytiva) | pGEX-**Oskar 139-240** (eLOTUS) | (Jeske *et al*, 2017), codon-optimized for expression in *E. coli* | A12-MJ |
| | pGEX-**Oskar 139-240 K236A** | Site directed mutagenesis of pGEX-Oskar 139-240 | RR99 |
| | pGEX-**Tejas 1-100** (eLOTUS) | (Jeske *et al*, 2017) | A59-MJ |
| | pGEX-**Tejas 1-100 SI** (S16E/I85E) | Site directed mutagenesis of pGEX-Tejas 1-100 | RR164 |
| | pGEX-**Tejas 1-100 C-MUT** (K94A/K96A/R98A) | Site-directed mutagenesis of pGEX-Tejas 1-100 | DH10 |
| | pGEX-**Tejas 1-93 / C-DEL** | Deletion PCR using pGEX-Tejas 1-100 | DH11 |
| | pGEX-**Tejas 1-100 K46E/K50E** | Site directed mutagenesis of pGEX-Tejas 1-100 | RR153 |
| | pGEX-**Tejas 1-100 R32E/R35E** | Site directed mutagenesis of pGEX-Tejas 1-100 | RR154 |
| | pGEX-**Tejas 1-100 6xMUT** (E28A/M31A/R32E/R35E/Q70A/M71A) | Site-directed mutagenesis of pGEX-Tejas 1-100 | ANG65 |
| **pHis-MCN** (NdeI/XbaI) (T18-MJ) (Jeske *et al*, 2015) | His-MCN-**BmTDRD7 1-100** (eLOTUS) *(Bombyx mori)* | (Jeske *et al*, 2017), codon-optimized for expression in *E. coli* | B66-MJ |
| | His-MCN-**BmTDRD7 1-100 MUT** (A13E/I81E) *(Bombyx mori)* | Site-directed mutagenesis of His-MCN-BmTDRD7 1-100 | ANG43 |
| | His-MCN-**Oskar 139-240** (eLOTUS) | (Jeske *et al*, 2015), codon-optimized for expression in *E. coli* | B7-MJ |
| | His-MCN-**Oskar 139-236** | Deletion PCR using His-MCN-Oskar 139-240 | ANG45 |
| | His-MCN-**Oskar-eLOTUS C-Tejas** (Oskar 139-236-Tejas-94-100) | PCR amplification of His-MCN-Osk 139-236 introducing Tejas 94-100 and religation | ANG42 |
| **pMJ-His** (ScaI) (T43-MJ) (Kubíková *et al*, 2023) | pMJ-His-**Tejas 1-100** (eLOTUS) | pGEX-Tejas 1-100 | RR292 |
| | pMJ-His-**Tejas 1-100 SI** (S16E/I85E) | pGEX-Tejas 1-100 SI | ANG48 |
| | pMJ-His-**Tejas 1-100 C-MUT** (K94A/K96A/R98A) | pGEX-Tejas 1-100 C-MUT | ANG6 |
| | pMJ-His-**Tejas 1-93 / C-DEL** | pGEX-Tejas C-DEL | ANG5 |

| | | | |
|---|---|---|---|
| | pMJ-His-**Tejas-eLOTUS C-Oskar**<br>(Tejas 1-93 - Oskar 237-240) | PCR amplification of pMJ-His-Tejas C-DEL introducing Osk 237-240 followed by religation | ANG36 |
| | pMJ-His-**Vasa 200-623** | His-MCN-Vasa 200-623 | RR60 |
| | pMJ-His-**Vasa 200-623 K295N** (Vasa open) | Site directed mutagenesis of pMJ-His-Vasa 200-623 | RR170 |
| | pMJ-His-**Vasa 200-623 Q333A** | Site directed mutagenesis of pMJ-His-Vasa 200-623 | RR172 |
| | pMJ-His-**Vasa 200-661***<br>(His-Vasa 200-661 GSGSGS) | PCR amplification of His-MCN-Vasa 200-661 introducing the GS linker followed by re-ligation | RR283 |
| | pMJ-His-**Vasa 200-661* F504E** | PCR amplification of His-MCN-Vasa 200-661 F504E introducing the GS linker followed by re-ligation | RR284 |
| | pMJ-**His-Vasa 200-661** | Deletion of the GS linker from pMJ-His-Vasa 200-661* | ANG37 |
| | pMJ-His-**Vasa-eLOTUS WT**<br>(Vasa 200-661_GSGSGS_Tejas 1-100) | PCR amplification and ligation from pMJ-His-Vasa 200-661* and pMJ-His-Tejas 1-100 | RR286 |
| | pMJ-His-**Vasa-eLOTUS I-MUT**<br>(Vasa 200-661 F504E_GSGSG_Tejas 1-100 S16E/I85E) | PCR amplification and ligation from pMJ-His-Vasa 200-661* F504E and pMJ-His-Tejas 1-100 SI | RR287 |
| | pMJ-His-**Vasa-eLOTUS C-MUT**<br>(Vasa 200-661_GSGSGS_Tejas 1-100 K94A/K96A/R98A) | Site-directed mutagenesis of pMJ-His-Vasa-eLOTUS WT | AK32 |
| | pMJ-His-**Vasa-eLOTUS C-DEL**<br>(Vasa 200-661_GSGSGS_Tejas 1-93) | Deletion PCR using pMJ-His-Vasa-eLOTUS WT | AK31 |
| | pMJ-His-**Vasa CTD-eLOTUS WT**<br>(Vasa 463-661_GSGSGS_Tejas 1-100) | Deletion PCR using pMJ-His-Vasa-eLOTUS WT | ANG1 |
| | pMJ-His-**Vasa CTD-eLOTUS I-MUT**<br>(Vasa 463-661 F504E_GSGSGS_Tejas 1-100 S16E/I85E) | Deletion PCR using pMJ-His-Vasa-eLOTUS I-MUT | ANG2 |
| | pMJ-His-**HaloTag-Vasa 200-661**<br>(HT-Vasa) | Gibson assembly using pET15b-His-Halo (Lolicato *et al*, 2022) and pMJ-His-Vasa 200-661 | ANG49 |
| **pMJ-His-MBP**<br>(ScaI)<br>(T46-MJ)<br>(Kubíková *et al*, 2023) | pMJ-His-MBP-**Vasa** FL | Synthetic sequence codon-optimized for *E. coli* | JK159 |
| | pMJ-His-MBP-**Vasa core** (aa 200-623) | Deletion PCR using pMJ-His-MBP-Vasa FL | pMH2194 |

| | | | |
|---|---|---|---|
| **pET-24a(+)** (NdeI/XhoI) (Novagen) | pET-24a-**Bm-Vasa NTD** (aa 167-397) | Synthetic sequence codon-optimized for *E. coli* | - |
| | pET-24a-**Bm-Vasa CTD** (aa 403-564) | Synthetic sequence codon-optimized for *E. coli* | - |
| **pAc5.1-EGFP** (EcoRV) (T5-MJ) (Tritschler *et al*, 2007) | pAc5.1-EGFP-**Vasa** iso A | (Salgania *et al*, 2024) | F15-MJ |
| | pAc5.1-EGFP-**Vasa open** (K295N) | (Salgania *et al*, 2024) | RR228 |
| | pAc5.1-EGFP-**Vasa closed** (E400Q) | (Salgania *et al*, 2024) | F16-MJ |
| | pAc5.1-EGFP-**Tejas** iso A | (Jeske *et al*, 2017) | F34-MJ |
| **pAc5.1-PH-mCherry** (FspAI) (HK49) (Salgania *et al*, 2024) | pAc5.1-PH-mCherry-**Tejas** iso A | (Salgania *et al*, 2024) | RR117 |
| | pAc5.1-PH-mCherry-**Vasa open** (K295N) | (Salgania *et al*, 2025) | RR132 |
| | pAc5.1-PH-mCherry-**Vasa closed** (E400Q) | (Salgania *et al*, 2025) | RR131 |
| **pBlue-script SK (-)** (Strata-gene) (subcloning vector) | pBSK-**Tejas WT** (*tejas* 5'UTR_CDS (Tejas-mEGFP)_*tejas* 3'UTR) | *Drosophila* cDNA (Tejas), synthetic DNA (mEGFP) | RR239 |
| | pBSK-**Tejas OL** (*tejas* 5'UTR_CDS (Oskar-139-240-Tejas-101-559-mEGFP)_*tejas*-3'UTR) | Oskar-eLOTUS amplified from *Drosophila* cDNA inserted into opened pBSK-Tejas-WT lacking Tejas-eLOTUS | RR294 |
| | pBSK-**Tejas SI** (*tejas* 5'UTR_CDS (Tejas S16E/I85E-mEGFP)_*tejas*-3'UTR) | Site directed mutagenesis of pBSK-Tejas-WT | RR236 |
| **attB** (Jeske *et al*, 2017) | attB-**Oskar WT** ($p_{osk}$_genomic oskar) | see **Supplementary Methods** | K7-MJ |
| | attB-**Oskar TL** ($p_{osk}$_genomic oskar containing Tejas-eLOTUS instead of Oskar-eLOTUS) | see **Supplementary Methods** | RR111 |
| | attB-pUAS-**Tejas WT** (pUAS_*tejas* 5'UTR_CDS (Tejas-mEGFP)_*tejas*-3'UTR) | pBSK-Tejas WT (AgeI and BglII) | RR108 |
| | attB-pUAS-**Tejas SI** (pUAS_*tejas* 5'UTR_CDS (Tejas S16E/I85E-mEGFP)_*tejas*-3'UTR) | pBSK-Tejas SI (AgeI and BglII) | RR233 |
| | attB-pUAS-**Tejas OL** (pUAS_*tejas* 5'UTR_CDS (Tejas-139-240-Oskar-101-559-mEGFP)_*tejas*-3'UTR) | pBSK-Tejas OL (AgeI and BglII) | RR109 |
| | attB-pUAS-**Tejas C-MUT** (pUAS_*tejas* 5'UTR_CDS (Tejas K94A/K96A/R98A/K102A/K104A -mEGFP)_*tejas*-3'UTR) | Gibson assembly using pMJ-His-Tejas 1-100 C-MUT, pBSK-Tejas WT, and attB-pUAS-Tejas WT | HK324 |
| | attB-pUAS-**Tejas C-DEL** (pUAS_*tejas* 5'UTR_CDS (Tejas ∆94-104-mEGFP)_*tejas*-3'UTR) | Gibson assembly using pAc5.1-mCherry-Tejas C_DEL and attB-pUAS-Tejas WT | ANG31 |

### Supplementary Table 4. Purification and application of recombinant proteins used in this study

| Protein | Purification steps (material) | Application |
|---|---|---|
| GST | GSH agarose, HiPrep Q, HiLoad 200 pg | GST pull-down assay, dsRNA unwinding assay |
| GST-**Oskar** eLOTUS | GSH agarose, HiPrep Q, HiLoad 200 pg | GST pull-down assay, ATPase assay (TLC) |
| GST-Oskar eLOTUS K236A | GSH agarose, HiPrep Q, HiLoad 200 pg | GST pull-down, ATPase assay (TLC) |
| His-Oskar-eLOTUS | $Ni^{2+}$-NTA agarose, HiPrep Q, HiLoad 200 pg | dsRNA unwinding assay |
| His-Oskar eLOTUS R238A/T239A | $Ni^{2+}$-NTA agarose, HiPrep Q, HiLoad 200 pg | ATPase assay, dsRNA unwinding assay, GST pull-down |
| His-Oskar-eLOTUS C-Tejas | $Ni^{2+}$-NTA agarose, HiPrep Q, HiLoad 200 pg | dsRNA unwinding assay |
| GST-**Tejas** eLOTUS | GSH agarose, HiPrep Q, HiLoad 200 pg | dsRNA unwinding assay, GST pull-down assay |
| GST-Tejas eLOTUS SI | GSH agarose, HiPrep Q, HiLoad 200 pg | dsRNA unwinding assay, GST pull-down assay |
| GST-Tejas eLOTUS C-MUT | GSH agarose, HiPrep Q, HiLoad 200 pg | GST pull-down, dsRNA unwinding assay |
| GST-Tejas eLOTUS C-DEL | GSH agarose, HiPrep Q, HiLoad 200 pg | GST pull-down, dsRNA unwinding assay |
| GST-Tejas eLOTUS K46E/K50E | GSH agarose, HiPrep Q, HiLoad 200 pg | EMSA |
| GST-Tejas eLOTUS R32E/K35E | GSH agarose, HiPrep Q, HiLoad 200 pg | EMSA |
| GST-Tejas eLOTUS 6xMUT | GSH agarose, HiPrep Q, HiLoad 200 pg | EMSA |
| His-Tejas eLOTUS | $Ni^{2+}$-NTA agarose, HiPrep SP, HiLoad 200 pg | ATPase assays (TLC, NADH), dsRNA unwinding assay, BLI, ITC, EMSA, condensation assay |
| His-Tejas eLOTUS SI | $Ni^{2+}$-NTA agarose, HiPrep SP, HiLoad 75 pg | ATPase assay (TLC), condensation assay |
| His-Tejas eLOTUS C-MUT | $Ni^{2+}$-NTA agarose, HiPrep SP, HiLoad 75 pg | ATPase assays (TLC, NADH), condensation assay |
| His-Tejas eLOTUS C-DEL | $Ni^{2+}$-NTA agarose, HiPrep SP, HiLoad 75 pg | ATPase assay (TLC), condensation assay |
| His-Tejas-eLOTUS C-Oskar | $Ni^{2+}$-NTA agarose, HiPrep SP, HiLoad 75 pg | ssRNA unwinding assay |
| His-**Vasa** 200-661 | $Ni^{2+}$-NTA agarose, HiPrep heparin, HiLoad 200 pg | ATPase assay (TLC NADH), dsRNA unwinding assay, BLI |
| His-Vasa 200-661* | $Ni^{2+}$-NTA agarose, HiPrep heparin, HiLoad 200 pg | ATPase assay (TLC), ds RNA unwinding assay, BLI, GST pull-down assay |
| His-Vasa 200-661 F504E | $Ni^{2+}$-NTA agarose, HiPrep heparin, HiLoad 200 pg | ATPase assay (TLC) |
| His-Vasa 200-623 open (K295N) | $Ni^{2+}$-NTA agarose, HiPrep heparin, HiLoad 200 pg | GST pull-down |

| | | |
|---|---|---|
| His-Vasa 200-623 Q333A | $Ni^{2+}$-NTA agarose, HiPrep heparin, HiLoad 200 pg | GST pull-down, EMSA |
| His-Vasa-eLOTUS WT | $Ni^{2+}$-NTA agarose, HiPrep Q, HiLoad 200 pg | ATPase assay (TLC), dsRNA unwinding assay, BLI |
| His-Vasa-eLOTUS I-MUT | $Ni^{2+}$-NTA agarose, Hi Prep heparin, HiLoad 200 pg | ATPase assay (TLC), dsRNA unwinding assay, BLI |
| His-Vasa-eLOTUS C-MUT | $Ni^{2+}$-NTA agarose, HiPrep Q, HiLoad 200 pg | BLI |
| His-Vasa-eLOTUS C-DEL | $Ni^{2+}$-NTA agarose, HiPrep Q, HiLoad 200 pg | BLI |
| His-Vasa-CTD-eLOTUS WT | $Ni^{2+}$-NTA agarose, HiPrep Q, HiLoad 200 pg | BLI |
| His-Vasa-CTD-eLOTUS I-MUT | $Ni^{2+}$-NTA agarose, HiPrep Q, HiLoad 200 pg | BLI |
| His-HaloTag-Vasa | $Ni^{2+}$-NTA agarose, HiPrep heparin, HiLoad 200 pg | ATPase assays (TLC, NADH) |
| Vasa FL (His-MBP) | $Ni^{2+}$-NTA Sepharose, 3C cleavage, HiLoad 200 pg | condensation assay |
| Vasa 200-623 (His-MBP) | $Ni^{2+}$-NTA Sepharose, 3C cleavage, HiLoad 200 pg | condensation assay |
| **Bm**-Vasa (His) | HisTrap, TEV cleavage, Heparin, S200 gel filtration | ATPase assay (TLC), dsRNA unwinding assay |
| Bm-Vasa NTD (His) | HisTrap, TEV cleavage, DEAE column, S200 gel filtration | NMR |
| Bm-Vasa CTD (His) | HisTrap, TEV cleavage, DEAE column, S200 gel filtration | NMR |
| His-Bm-TDRD7-eLOTUS | $Ni^{2+}$-NTA agarose, HiPrep SP, HiLoad 75 pg | ATPase assay, dsRNA unwinding assay, NMR |
| His-Bm-TDRD7-eLOTUS MUT | $Ni^{2+}$-NTA agarose, HiPrep SP, HiLoad 75 pg | ATPase assay (TLC) |

### SUPPLEMENTARY METHODS

#### Vasa immunofluorescence

One-day-old transgenic female flies were incubated for two days on a medium without yeast in the presence of $w^{1118}$ males. Ovaries were dissected and briefly stored in 1 x PBS. After removing PBS, the ovaries were fixed by incubation for 3 minutes in 500 µL preheated (92°C) 0.4% (w/v) NaCl, 0.3% Tween 20 solution. Then, 1 mL of the same solution at ice-cold temperature was added and incubation continued for 2 more minutes. Following fixation, ovaries were washed at room temperature with 1 mL of PBS-T (PBS supplemented with 0.1% Tween 20), and incubated for 1 hour at room temperature in PBS-T supplemented with 10% normal goat serum (GibcoBRL). The blocking solution was replaced with 200 µL of PBS-T supplemented with 10% Normal Goat Serum and guinea pig anti-Vasa antibody (1:2500) and samples were incubated overnight at 4°C on a rotary shaker. The next day, the ovaries were washed three times with 1 mL PBS-T. To protect the fluorophore group of the secondary antibodies from quenching, subsequent steps were performed using brown tubes to protect the sample from light. Ovaries were incubated for 2-3 hours at room temperature with Alexa Fluor™ 594-conjugated anti-guinea pig antibodies (Invitrogen) 1:1000 diluted in PBS-T supplemented with 10% normal goat serum, followed by two washing steps with PBS-T of 10 minutes each. During the first wash, nuclei were stained for 5 minutes with DAPI (Roth; 1:2500 in 1x PBS-T). After PBS-T removal, 100 µL of Vectashield® Antifade Mounting Media (VectorLabs) was added. Ovaries were either stored overnight or directly mounted onto a glass slide and sealed with a cover slip and nail polish. Imaging was performed using 20 x objective and a Zyla 4.2P camera on a Nikon Ti2-W1 spinning disk confocal microscope and the resulting images analyzed using Fiji (Schindelin *et al*, 2012).

#### Preparation of Oskar transgenes

The *osk*-STOP-cTRICK line is an Oskar protein null line, containing two stop codons at positions 159 (Glu) and 164 (Ile).

To generate attB-Oskar WT (K7-MJ), the BamHI/ApaI fragment of the genomic oskar sequence was amplified from the pDM30-g-osk vector (Ephrussi *et al*, 1991) using primers that introduced KpnI sites at both ends and subcloned into the PCR Blunt II-TOPO vector (Life Technologies) to generate the TOPO-Oskar WT (J1-MJ) subclone. The *oskar* promoter was excised from pDM30-g-osk using XhoI and BamHI. The K10 signal sequence in pUAS-K10attB (Koch *et al*, 2009) was removed by NdeI/XbaI digestion, filling in the 5′ overhangs, and re-ligating the blunt ends, and the UAS promoter sequence was removed by XhoI/KpnI digestion. Finally, the genomic *oskar* fragment was released from the TOPO-Oskar WT vector with BamHI/KpnI and ligated together with the XhoI/BamHI *oskar* promoter fragment into the XhoI/KpnI sites of the modified attB vector.

The attB-Oskar TL transgene was generated as follows. First, Oskar residues 139-173 and 183-240 were removed from the TOPO-Oskar WT subclone by two successive deletion PCR reactions. The remaining region encompassing residues 174-182 contains intron 1 and its flanking sequences, which upon splicing form the structured SOLE element required for *oskar* mRNA localization (Simon *et al*, 2015; Ghosh *et al*, 2012; Hachet & Ephrussi, 2004). Next, Tejas residues 1-100 were amplified from pBSK-Tejas WT (RR239) and inserted upstream of the retained *oskar* intron 1, generating TOPO-Oskar TL. Finally, TOPO-Oskar TL and attB-Oskar WT (K7-MJ) were digested with BamHI and MluI to release genomic *oskar* fragments containing either the Tejas-eLOTUS or Oskar-eLOTUS domain, respectively. The fragment containing the Tejas-eLOTUS domain was inserted into the attB-Oskar WT backbone lacking the Oskar-eLOTUS region, generating attB-Oskar TL.

The attB-Oskar WT and attB-Oskar TL transgenes were integrated into the attP40 site 25C6 on chromosome 2R using ΦC31 integrase via the *Drosophila* injection service of the University of Cambridge.

**MALS**

Multiangle-light-scattering (MALS) experiments were performed at 4°C using an ÅKTA purifier system (Cytiva) equipped with a Superdex 75 Increase 10/300 column (Cytiva) and connected to a DAWN 8+ light scattering detector (Wyatt Technology) and SEC-3010 RI refractive index detector (WGE Dr Bures GmbH, Germany). The system and the proteins were in buffer containing 20 mM Tris-Cl, pH 7.5 and 150 mM NaCl. 220 µl of His-Tejas 1-100 protein with a concentration of 1.2 mM was injected. The data were fitted and analyzed with the Astra 6 software (Wyatt Technology).

**ATPase assay in the presence of RNases**

In a total reaction volume of 50 µL, either 5 µL of 125 µM HT-Vasa or 16 µl of 1.56 mM His-Tejas eLOTUS were mixed with either 5 µL of Benzonase (250 U/µL; Millipore) and 40 µl Benzonase buffer (1 mM $MgCl_2$ and 20 mM Tris pH 8) or 5 µL of an RNase A/T1 mix (2,5 mg/mL RNase A, 250 U/µL RNase T1; both from Thermo Fisher Scientific) and 40 µl protein buffer (150 mM $NaCl_2$, 20 mM Tris pH 7.5). Samples were incubated for 1 h at 25°C. Control samples (“Buffer”) were treated identically but without addition of RNases. To verify RNase activity, 5 µl of 1 mM single-stranded R26 RNA oligonucleotide was added to the RNase and control samples before incubation. Following treatment, the proteins were used in the NADH-coupled ATPase assay at the concentrations indicated, using 5 mM 1:1 $MgCl_2$-ATP mixture.

**TERRA G4 substrates**

TERRA G4 substrates were prepared similar to as described (Ding *et al*, 2020). 6-FAM-labeled TERRA G4 (UUAGGGUUAGGGUUAGGGUUAGGG) and TERRA G4 MUT (UUACCGUUACCGUUACCGUUACCG) were purchased from IDT (100 nmol synthesis scale) and dissolved in 10 mM Tris-Cl pH 7.4 and 100 mM KCl to a final RNA concentration of 100 µM. The RNAs were incubated for 5 min at 95°C and slowly cooled in a thermal block to 16°C over approximately 2 h.

**EMSA**

To test RNA binding by His-Vasa WT, 50 nM 5'-$^{32}$P-labeled ssR26 ($(GCUUUACGGUGU)_2$; IDT) were incubated with 14.7 µM protein (and several 2-fold dilutions) for 30 min at 25°C in a buffer containing 150 mM NaCl, 20 mM Tris, pH 7.5, 10% glycerol, 1 mM ATP, 5 mM $MgCl_2$, and 1 mM DTT. The His-Vasa Q333A mutant was tested similarly in the absence or presence of 150 µM His-Tejas-eLOTUS.

To test RNA binding by the Tejas-eLOTUS domain, 50 nM 5'-$^{32}$P-labeled ssR26 or dsR26 were incubated with 150 µM His-Tejas-eLOTUS (and several 2-fold dilutions) for 30 min at 25°C in a buffer containing 150 mM NaCl, 20 mM Tris, pH 7.5, 10% glycerol, and 1 mM DTT.

EMSAs using G4 RNA were performed similarly as described (Ding *et al*, 2020). 250 nM TERRA G4 or TERRA G4 MUT RNA were mixed with GST-eLOTUS domains at different concentrations in a buffer containing 25 mM Tris-Cl pH 7.4, 150 mM KCl, 0.5 mM DTT, 5 mM EDTA, 0.5% NP-40 and 1 U/µl RiboLock (Thermo Fisher Scientific) for 30 min at 4°C.

All samples were separated on a 6% polyacrylamide (37.5:1) gel using 0.5 x TBE pH 9.5 as running buffer in a cold room. Radioactive gels were dried and analyzed using phosphorimaging, fluorescent gels were analyzed using a Sapphire FL Biomolecular Imager. All images were quantified using Fiji (Schindelin *et al*, 2012).

**Protein expression and purification for condensation assays**

Recombinant proteins were expressed in *E. coli* Lemo21(DE3) cells transformed with pET-MCN-based expression plasmids encoding full-length Vasa (Vasa FL) or the Vasa helicase core (aa 200-623) under kanamycin selection. Pre-cultures were grown overnight at 37°C in Terrific Broth (TB) medium supplemented with antibiotics and 1% (w/v) D-glucose. Expression cultures were inoculated into 2 L TB medium containing antibiotics and 1% (w/v) D-glucose to an initial OD600 of 0.05 and grown at 37°C to an OD600 of 0.4-0.6. Protein expression was induced with 250 µM IPTG, and cultures were incubated overnight at 20°C. Cells were harvested by centrifugation at 5,000 × g for 15 min at 4°C. Cell pellets were resuspended in 80 mL lysis buffer consisting of 50 mM Tris-HCl (pH 7.5), 1 M NaCl, 10% (w/v) glycerol, 100 µM PMSF (Roth), 1 x protease inhibitor cocktail (1000 x stock: 10 mM bestatin (Sigma), 2 mM E64 (Pepta Nova), 1 mM pepstatin A (Pepta Nova), 10 mM 1.10-phenanthrolin (Fisher Scientific) and 1mM phosphoramidon (Pepta Nova)), 2 µg/µL DNase I (Roche, Cat# 10104159001), and 60 µg/mL RNase A (Macherey-Nagel, Cat# 74050). Cells were lysed by pressure homogenization using an EmulsiFlex (Avestin) (two passes), followed by sonication for 2 min at 40% amplitude using a 2 s ON/OFF cycle. Lysates were cleared by centrifugation at 40,000 × g for 30 min at 4°C and filtered through a 0.45 µm cut-off filter. His-MBP-3C-tagged proteins were purified by immobilized metal affinity chromatography (IMAC) on an Äkta System (GE Life Sciences) using self-packed 5 ml columns with $Ni^{2+}$ Sepharose High Performance resin (GE Healthcare). The solubility tag was removed by overnight cleavage with 3C protease at 4°C during dialysis into storage buffer containing 50 mM Tris-HCl pH 7.5, 1 M NaCl, 10% glycerol (w/v), 2 mM $MgCl_2$ and 3 mM β-mercaptoethanol. Proteins were further purified by size exclusion chromatography (SEC) using a 16/600 HiLoad Superdex 200 pg column (Cytiva) equilibrated in storage buffer. Clean SEC fractions corresponding to monomeric protein were pooled and concentrated to 300-500 µM using Amicon Ultra-4 centrifugal filters (Merck Millipore). Concentrated protein aliquots were snap-frozen in liquid nitrogen and stored at -80°C.

**Chemical labeling of proteins**

Untagged recombinant proteins Vasa-FL, Vasa-core, and BSA were chemically labeled with ATTO488 NHS-ester dye (ATTO-TEC GmbH, Cat# AD-488-31), and His-eLOTUS WT, the mutant variants (SI, C-DEL, C-MUT), and BSA were labeled with ATTO565 (ATTO-TEC GmbH, Cat# AD-488-31). To specifically label the primary amine of the N-terminus and not of the lysines, labeling reactions were performed at pH 6.5. Proteins and ATTO NHS-ester dyes were mixed at a molar ratio of 1:4, followed by 1 h incubation at room temperature, protected from light. Using Zeba™ Spin Desalting Columns (Thermo Fisher Scientific), free dye was removed and the labelled protein was exchanged into storage buffer (50 mM Tris-HCl pH 7.5, 1 M NaCl, 10% glycerol (w/v), 2 mM $MgCl_2$ and 3 mM β-mercaptoethanol). Finally, all NHS-labeled proteins were concentrated using Amicon Ultra-4 centrifugal filters (Merck Millipore) to 50-100 µM into storage buffer and snap-frozen in liquid nitrogen and stored at -80°C.

***In vitro* condensation assay**

Condensation assays were carried out in 384-well, black, optically clear, flat-bottom, ultra-low-attachment-coated PhenoPlates (Revvity, Cat# 6057800) as previously described (Dörner *et al*, 2026). Protein stocks were diluted in protein storage buffer to a concentration of 10 x of the final assay concentration. For visualization of untagged protein variants, ATTO-labeled protein was spiked into the 10 × protein stocks at a final labeling fraction of 1%. The 10 × protein stocks were transferred to individual wells and mixed with the assay master mix at a ratio of 1:10 to yield the final assay conditions: 5 µM Vasa or BSA and 25 µM LOTUS domain variant or BSA in reaction buffer containing 50 mM MES (pH 5.7), 50 mM NaCl, 2 mM ATP, 0.5 µg/µL BSA, and, where indicated, 50 ng/µL polyU RNA (Sigma, Cat# P9528-25MG). After reaction assembly, the plate was centrifuged at 100 × *g* for 1 minute, and incubated at 25°C on the stage of the temperature-controlled microscope prior imaging. The time between reaction assembly and imaging was timed and controlled for each well, resulting in a constant total incubation time for each reaction of 30 minutes, to allow for direct comparison. Images were acquired on a temperature-controlled, inverted Nikon Ti2 widefield microscope equipped with a 40 × Plan Apo Lambda air objective NA 0.95, a Lumencor SPECTRA light source and a Hamamatsu ORCA-Fusion cMOS camera. All conditions were imaged using Differential Interference Contrast (DIC) microscopy and the appropriate fluorescence channels. Four images were acquired in each well. Microscope operation and acquisition were controlled using Nikon NIS-Elements software with automated acquisition via the JOBS module. Illumination intensity and exposure times were optimized for each fluorescent label and kept constant within the experiment. Images were processed by adjusting brightness and contrast using the open microscopy environment (OMERO). For each panel, representative images are shown.

**Quantification of condensation assays**

Images were analyzed using Nikon NIS-Elements GA3 software (Nikon) with the integrated Segment.ai module. A representative subset of images from different LLPS experiments was used to train the AI-based segmentation model to identify droplets based on the raw GFP (488 nm) signal. The trained model was subsequently applied to all images to automatically segment droplets. Droplets touching the image borders were excluded from further analysis. For each segmented droplet, the mean fluorescence intensity inside the droplet and in the surrounding was extracted for the 488 nm and/or 565 nm channels using GA3 analysis tools. The partition coefficient was calculated as the ratio of mean fluorescence intensity inside droplets to the mean intensity of the surrounding. For each condition, four independent images were analyzed. The mean partition coefficient per image was calculated and used for downstream analysis. Samples containing fewer than 10 segmented droplets were excluded from the downstream analysis. This threshold was applied as a quality-control measure to ensure that segmented objects represented bona fide phase-separated detected by the segmentation algorithm. Total droplet number per condition was calculated by summing all segmented droplets across images. Data processing, filtering, and plotting of mean partition coefficients and droplet numbers were performed in R (RStudio) using custom scripts available upon request.

### SUPPLEMENTARY REFERENCES

Andersen CBF, Ballut L, Johansen JS, Chamieh H, Nielsen KH, Oliveira CLP, Pedersen JS, Séraphin B, Hir H Le & Andersen GR (2006) Structure of the exon junction core complex with a trapped DEAD-Box ATPase bound to RNA. *Science (1979)* 313: 1968–1972

Andreou AZ, Harms U & Klostermeier D (2017) eIF4B stimulates eIF4A ATPase and unwinding activities by direct interaction through its 7-repeats region. *RNA Biol* 14: 113–123

Andreou AZ & Klostermeier D (2014) EIF4B and eIF4G jointly stimulate eIF4A ATPase and unwinding activities by modulation of the eIF4A conformational cycle. *J Mol Biol* 426: 51–61

Askjaer P, Rosendahl R & Kjems J (2000) Nuclear Export of the DEAD Box An3 Protein by CRM1 Is Coupled to An3 Helicase Activity. *Journal of Biological Chemistry* 275: 11561–11568

Bizebard T, Ferlenghi I, Iost I & Dreyfus M (2004) Studies on three E. coli DEAD-box helicases point to an unwinding mechanism different from that of model DNA helicases. *Biochemistry* 43: 7857–7866

Blum S, Schmid SR, Pause A, Buser P, Linder P, Sonenberg N & Trachsel H (1992) ATP hydrolysis by initiation factor 4A is required for translation initiation in Saccharomyces cerevisiae. *Proceedings of the National Academy of Sciences* 89: 7664–7668

Bono F, Ebert J, Lorentzen E & Conti E (2006) The Crystal Structure of the Exon Junction Complex Reveals How It Maintains a Stable Grip on mRNA. *Cell* 126: 713–725

Cao W, Coman MM, Ding S, Henn A, Middleton ER, Bradley MJ, Rhoades E, Hackney DD, Pyle AM & De La Cruz EM (2011) Mechanism of Mss116 ATPase reveals functional diversity of DEAD-Box proteins. *J Mol Biol* 409: 399–414

Chamot B, Colvin KR, Kujat-Choy SL & Owttrim GW (2005) RNA Structural Rearrangement via Unwinding and Annealing by the Cyanobacterial RNA Helicase, CrhR. *Journal of Biological Chemistry* 280: 2036–2044

Chen Y, Boland A, Kuzuoğlu-Öztürk D, Bawankar P, Loh B, Chang C Te, Weichenrieder O & Izaurralde E (2014) A DDX6-CNOT1 Complex and W-Binding Pockets in CNOT9 Reveal Direct Links between miRNA Target Recognition and Silencing. *Mol Cell* 54: 737–750

Davila Gallesio J, Hackert P, Bohnsack KE & Bohnsack MT (2020) Sgd1 is an MIF4G domain-containing cofactor of the RNA helicase Fal1 and associates with the 5’ domain of the 18S rRNA sequence. *RNA Biol* 17: 539–553

Ding D, Wei C, Dong K, Liu J, Stanton A, Xu C, Min J, Hu J & Chen C (2020) LOTUS domain is a novel class of G-rich and G-quadruplex RNA binding domain. *Nucleic Acids Res* 48: 9262–9272

Donsbach P, Kwas C, Steimer L, Samatanga B, Andreou AZ & Klostermeier D (2025) Inter-domain communication in the dimeric DEAD-box helicase Hera from T. thermophilus and implications for the mechanism of RNA unwinding. *Nucleic Acids Res* 53: 80

Dörner K, Gut MJ, Overwijn D, Cao F, Siketanc M, Heinrich S, Beuret N, Meyer J, Sharpe T, Lindorff-Larsen K, *et al* (2026) Fluorescent protein and peptide tags alter condensate formation and dynamics in vivo and in vitro. *EMBO Rep* 27: 89–121

Ephrussi A, Dickinson LK & Lehmann R (1991) Oskar Organizes the Germ Plasm and Directs Localization of the Posterior Determinant Nanos. *Cell* 66: 37–50

Ghosh S, Marchand V, Gáspár I & Ephrussi A (2012) Control of RNP motility and localization by a splicing-dependent structure in oskar mRNA. *Nat Struct Mol Biol* 19: 441–449

Granneman S, Lin CY, Champion EA, Nandineni MR, Zorca C & Baserga SJ (2006) The nucleolar protein Esf2 interacts directly with the DExD/H box RNA helicase, Dbp8, to stimulate ATP hydrolysis. *Nucleic Acids Res* 34: 3189–3199

Gray S, Cao W, Montpetit B & De La Cruz EM (2022) The nucleoporin Gle1 activates DEAD-box protein 5 (Dbp5) by promoting ATP binding and accelerating rate limiting phosphate release. *Nucleic Acids Res* 50: 3998–4011

Hachet O & Ephrussi A (2004) Splicing of oskar RNA in the nucleus is coupled to its cytoplasmic localization. *Nature* 428: 959–963

Henn A, Cao W, Hackney DD & De La Cruz EM (2008) The ATPase Cycle Mechanism of the DEAD-box rRNA Helicase, DbpA. *J Mol Biol* 377: 193–205

Hilbert M, Kebbel F, Gubaev A & Klostermeier D (2011) eIF4G stimulates the activity of the DEAD box protein eIF4A by a conformational guidance mechanism. *Nucleic Acids Res* 39: 2260–2270

Jeske M, Bordi M, Glatt S, Müller S, Rybin V, Müller CW & Ephrussi A (2015) The crystal structure of the Drosophila germline inducer Oskar identifies two domains with distinct Vasa Helicase- and RNA-binding activities. *Cell Rep* 12: 587–598

Jeske M, Müller CW & Ephrussi A (2017) The LOTUS domain is a conserved DEAD-box RNA helicase regulator essential for the recruitment of Vasa to the germ plasm and nuage. *Genes Dev* 31: 939–952

Kellner JN, Reinstein J & Meinhart A (2015) Synergistic effects of ATP and RNA binding to human DEAD-box protein DDX1. *Nucleic Acids Res* 43: 2813–2828

Koch R, Ledermann R, Urwyler O, Heller M & Suter B (2009) Systematic functional analysis of Bicaudal-D serine phosphorylation and intragenic suppression of a female sterile allele of BicD. *PLoS One* 4

Kossen K, Karginov F V. & Uhlenbeck OC (2002) The Carboxy-terminal Domain of the DEx Protein YxiN is Sufficient to Confer Specificity for 23S rRNA. *J Mol Biol* 324: 625–636

Kossen K & Uhlenbeck OC (1999) Cloning and biochemical characterization of Bacillus subtilis YxiN, a DEAD protein specifically activated by 23S rRNA: Delineation of a novel sub-family of bacterial DEAD proteins. *Nucleic Acids Res* 27: 3811–3820

Kubíková J, Ubartaitė G, Metz J & Jeske M (2023) Structural basis for binding of *Drosophila* Smaug to the GPCR Smoothened and to the germline inducer Oskar. *Proceedings of the National Academy of Sciences* 120: e2304385120

Lolicato F, Saleppico R, Griffo A, Meyer A, Scollo F, Pokrandt B, Müller HM, Ewers H, Hähl H, Fleury JB, *et al* (2022) Cholesterol promotes clustering of PI(4,5)P2 driving unconventional secretion of FGF2. *J Cell Biol* 221

López De Victoria A, Moore AFT, Gittis AG & Koculi E (2017) Kinetics and Thermodynamics of DbpA Protein's C-Terminal Domain Interaction with RNA. *ACS Omega* 2: 8033–8038

Lorsch JR & Herschlag D (1998) The DEAD box protein eIF4A. 2. A cycle of nucleotide and RNA-dependent conformational changes. *Biochemistry* 37: 2194–2206

Marintchev A, Edmonds KA, Marintcheva B, Hendrickson E, Oberer M, Suzuki C, Herdy B, Sonenberg N & Wagner G (2009) Topology and Regulation of the Human eIF4A/4G/4H Helicase Complex in Translation Initiation. *Cell* 136: 447–460

Markussen FH, Michon AM, Breitwieser W & Ephrussi A (1995) Translational control of oskar generates short OSK, the isoform that induces pole plasm assembly. *Development* 121: 3723–3732

Mathys H, Basquin JÔ, Ozgur S, Czarnocki-Cieciura M, Bonneau F, Aartse A, Dziembowski A, Nowotny M, Conti E & Filipowicz W (2014) Structural and Biochemical Insights to the Role of the CCR4-NOT Complex and DDX6 ATPase in MicroRNA Repression. *Mol Cell* 54: 751–765

Montpetit B, Thomsen ND, Helmke KJ, Seeliger MA, Berger JM & Weis K (2011) A conserved mechanism of DEAD-box ATPase activation by nucleoporins and InsP6 in mRNA export. *Nature* 472: 238–244

Moore AFT, Gentry RC & Koculi E (2017) DbpA is a region-specific RNA helicase. *Biopolymers* 107: e23001

Noble CG & Song H (2007) MLN51 stimulates the RNA-helicase activity of eIF4AIII. *PLoS One* 2

Rocak S, Emery B, Tanner NK & Linder P (2005) Characterization of the ATPase and unwinding activities of the yeast DEAD-box protein Has1p and the analysis of the roles of the conserved motifs. *Nucleic Acids Res* 33: 999–1009

Rogers GW, Richter NJ, Lima WF & Merrick WC (2001) Modulation of the Helicase Activity of eIF4A by eIF4B, eIF4H, and eIF4F. *Journal of Biological Chemistry* 276: 30914–30922

Salgania HK, Metz J & Jeske M (2024) ReLo is a simple and rapid colocalization assay to identify and characterize direct protein–protein interactions. *Nature Communications 2024 15:1* 15: 1–12

Salgania HK, Metz J, Lingren E, Bleischwitz C, Hauser D, Máté KO, Bollack D, Lahr F, Bruckmann A, Garbelyanski A, *et al* (2025) Molecular insights into the Drosophila piRNA pathway via systematic ReLo protein interaction screening and structure prediction. *Nucleic Acids Res* 53: 13–14

Schindelin J, Arganda-Carreras I, Frise E, Kaynig V, Longair M, Pietzsch T, Preibisch S, Rueden C, Saalfeld S, Schmid B, *et al* (2012) Fiji: An open-source platform for biological-image analysis. *Nat Methods* 9: 676–682

Schütz P, Bumann M, Oberholzer AE, Bieniossek C, Trachsel H, Altmann M & Baumann U (2008) Crystal structure of the yeast eIF4A-eIF4G complex: an RNA-helicase controlled by protein-protein interactions. *Proc Natl Acad Sci U S A* 105: 9564–9

Simon B, Masiewicz P, Ephrussi A & Carlomagno T (2015) The structure of the SOLE element of oskar mRNA. *RNA* 21: 1444–1453

Sun Y, Atas E, Lindqvist L, Sonenberg N, Pelletier J & Meller A (2012) The eukaryotic initiation factor eIF4H facilitates loop-binding, repetitive RNA unwinding by the eIF4A DEAD-box helicase. *Nucleic Acids Res* 40: 6199–6207

Sun Y, Atas E, Lindqvist LM, Sonenberg N, Pelletier J & Meller A (2014) Single-molecule kinetics of the eukaryotic initiation factor 4AI upon RNA unwinding. *Structure* 22: 941–948

Tritschler F, Eulalio A, Truffault V, Hartmann MD, Helms S, Schmidt S, Coles M, Izaurralde E & Weichenrieder O (2007) A divergent Sm fold in EDC3 proteins mediates DCP1 binding and P-body targeting. *Mol Cell Biol* 27: 8600–8611

Tsu CA & Uhlenbeck OC (1998) Kinetic analysis of the RNA-dependent adenosinetriphosphatase activity of DbpA, an Escherichia coli DEAD protein specific for 23S ribosomal RNA. *Biochemistry* 37: 16989–16996

Virgili G, Frank F, Feoktistova K, Sawicki M, Sonenberg N, Fraser CS & Nagar B (2013) Structural Analysis of the DAP5 MIF4G Domain and Its Interaction with eIF4A. *Structure* 21: 517–527

Wong E V., Cao W, Vörös J, Merchant M, Modis Y, Hackney DD, Montpetit B & De La Cruz EM (2016) Pi Release Limits the Intrinsic and RNA-Stimulated ATPase Cycles of DEAD-Box Protein 5 (Dbp5). *J Mol Biol* 428: 492–508
